# From microbes to mammals: pond biodiversity homogenization across different land-use types in an agricultural landscape

**DOI:** 10.1101/2022.01.28.477988

**Authors:** D. Ionescu, M. Bizic, R. Karnatak, C. L. Musseau, G. Onandia, M. Kasada, S.A. Berger, J.C. Nejstgaard, M. Ryo, G. Lischeid, M. O. Gessner, S. Wollrab, H.-P. Grossart

## Abstract

Local biodiversity patterns are expected to strongly reflect variation in topography, land use, dispersal boundaries, nutrient supplies, contaminant spread, management practices and other anthropogenic influences. In contrast, studies focusing on specific taxa revealed a biodiversity homogenization effect in areas subjected to long-term intensive industrial agriculture. We investigated whether land use affects biodiversity and metacommunity structure in 67 kettle holes (KH) representing small aquatic islands embedded in the patchwork matrix of a largely agricultural landscape comprising grassland, forest, and arable fields. These KH, similar to millions of standing water bodies of glacial origin, spread across northern Europe, Asia, and North America, are physico-chemically diverse, differ in the degree of coupling with their surroundings. We assessed biodiversity patterns of eukaryotes, *Bacteria* and *Archaea* in relation to environmental features of the KH, using deep-amplicon-sequencing of eDNA. First, we asked whether deep sequencing of eDNA provides a representative picture of KH biodiversity across the three domains of life. Second, we investigated if and to what extent KH biodiversity is influenced by the surrounding land-use. Our data shows that deep eDNA amplicon sequencing is useful for in-depth assessments of cross-domain biodiversity comprising both micro- and macro-organisms, but, has limitations with respect to single-taxa conservation studies. Using this broad method, we show that sediment eDNA, integrating several years to decades, depicts the history of agricultural land-use intensification. The latter, coupled with landscape wide nutrient enrichment (including by atmospheric deposition), groundwater connectivity between KH and organismal movement in the tight network of ponds, resulted in a biodiversity homogenization in the KH water, levelling off today’s detectable differences in KH biodiversity between land-use types.

## Introduction

The cultural landscape of Central Europe was characterized by low-input farming till the 1950s and early 1960s, after which industrialized agriculture became dominant with greatly increased fertilizer and pesticide use (Bauerkämper 2004, Sommer et al. 2008). Concomitantly, crop diversity decreased by more than 30% while the total crop coverage of land increased (Meyer et al. 2013). These changes in agricultural practice had negative consequences on biodiversity, resulting in declining plant (Meyer et al. 2013, Altenfelder et al. 2014), bird (Donald et al. 2006), invertebrate (Wilson et al. 1999) and amphibian (Berger et al. 2011, 2018) diversity. Furthermore, plant communities became homogenized (Macdonald and Johnson 2000, Baessler and Klotz 2006), as has commonly been observed after land-use intensification (Smart et al. 2006).

Ponds are intimately linked to their terrestrial surroundings, both the riparian zones immediately adjacent to the water bodies and the entire watershed due to their small size and topographic position in landscape depressions (Søndergaard et al. 2005, Kayler et al. 2019). As a result, pond biodiversity tends to be particularly affected by land use (Declerck et al. 2006), resulting, for instance, in increased organic matter and nutrient supply; pesticide spread by aerial spray, run-off and groundwater flow (Pérez-Lucas et al. 2019), leading to changes in plant (Altenfelder et al. 2014) and animal (Berger et al. 2011) communities.

Kettle holes (KH) are small landscape depressions formed on the outwash plains in front of retreating glaciers at the end of the last ice age. Most fill with water, at least temporarily, which has resulted in parts of the post-glacial landscapes of northern Europe, northern North America, and northern Asia being sprinkled with these small water bodies (Downing et al. 2006). For example, more than 90,000 occur in northeastern Germany, with densities reaching up to 40 per km^2^ (Kalettka and Rudat 2006). KH can vary greatly in hydro-geomorphological and biological features, even when they are geographically close to one another (Attermeyer et al. 2017). Biological activity in KH is high (Nitzsche et al. 2017) and they also play a critical role as local biodiversity hotspots (Scheffer et al. 2006, Joniak et al. 2007, Lischeid and Kalettka 2012, Pätzig et al. 2012, Platen et al. 2016, Novikmec et al. 2016), serving as habitat for insects both with and without aquatic life stages, as refuge and breeding ground for many amphibians, and as feeding areas for aquatic as well as terrestrial species (Berger et al. 2013, Heim et al. 2018). Accordingly, KH host diverse communities, both aquatic and extending beyond aquatic boundaries.

Water filled KH are aquatic islands embedded in the terrestrial landscape, where local communities are connected via passive and active overland dispersal to form a metacommunity (Wilson 1992, Leibold et al. 2004). Such metacommunities are subjected to local dynamics for example via food-web interactions and regional dynamics by a multitude of mechanisms such as mass and rescue effects, colonization and deterministic and stochastic extensions (Leibold et al. 2004). The frequent occurrence of KH in the landscape suggests they serve as stepping stones between habitats located in different land use types within the landscape (Premke et al. 2016, Kayler et al. 2018). Accordingly, the biodiversity of small ponds such as KH is disproportionately high (Scheffer et al. 2006) compared to the terrestrial surroundings, and is directly linked to the degree of connectivity to other ponds (Van Geest et al. 2003).

Assessing biological diversity across taxa, from microbes to mammals, is challenging. However, a promising approach is the use of environmental DNA (eDNA) which provides a common denominator for all taxa independent of body size and other species traits. Therefore, the analysis of eDNA has been increasingly applied as a non-invasive, highly sensitive monitoring tool (Deiner et al. 2017, Harper et al. 2019). The approach is based on collecting samples of live, dead, or partially decomposed organisms containing DNA that can be amplified and taxonomically annotated. Consequently, in highly dynamic ecosystems such as KH, eDNA methods will show long-term changes in the environment but may miss immediate effects of the surrounding. Most eDNA approaches aim to detect a specific set of taxa such as mammals, fish or amphibians by making use of previously identified specific sequences, or omnipresent genomic markers, such as the small and large subunits of ribosomal RNA genes or the Cox genes (Deiner et al. 2017, Andújar et al. 2018, Bylemans et al. 2019, Beng and Corlett 2020). Despite several downsides that have been recognized, such as misinterpretation of sequence frequencies or of presence and absence of taxa (Roussel et al. 2015, Harper et al. 2019), the approach has proved particularly useful for analyses motivated by species-specific conservation or restoration efforts. Importantly, however, it is also possible to use eDNA to assess biological diversity as a whole, facilitated by the use of genomic markers that capture all life forms.

In the present study, we embarked on a multi-seasonal analysis of cross-taxa biodiversity patterns in contrasting KH using deep sequencing (>200,000 reads per sample separately for *Bacteria*, *Archaea*, eukaryotes) of eDNA. These KH are characterized by three different land-use types embedded in a agriculture dominated landscape: arable fields, grassland, and forest.

We defined three main goals of the study: 1) evaluate whether deep sequencing of eDNA is a reliable approach for broad qualitative biodiversity assessments, providing representative, in depth information on a wide range of organisms, 2) assess the magnitude of the effect land-use type has on aquatic biodiversity, and 3) test to what extent the observed landscape-scale patterns depend on land use type. We hypothesized that biodiversity in KH surrounded by grassland and particularly by forest represents a more natural state than kettle holes embedded in agriculture fields, resulting in richer local communities within individual KH (α diversity) and more heterogeneous communities across KH within the more natural land-use categories (β-diversity).

## Methods

### Study sites and sampling

Samples for eDNA analysis were collected during 5 sampling campaigns of 2-3 days each in December 2016, and March, May, June and October 2017. All samples were taken in a set of 67 kettle holes located in northeastern Germany (Fig. 1). The area, one of the least populated in Germany, has a long history of farming, with >90% of the land now being covered by arable fields (Kalettka and Rudat 2006), although some of that land has been reconverted to grassland nearly two decades ago (Serrano et al. 2017). Routine monitoring of the KH water and riparian vegetation in the area started in 1993, shortly after the reunification of Germany (Kalettka and Rudat 2006). Each KH was categorized based on land-use type within a perimeter of ca. 50 m around the kettle holes, i.e. distinguishing KH in arable fields, grasslands, and forest patches (Fig. 1).

**Figure 1.**
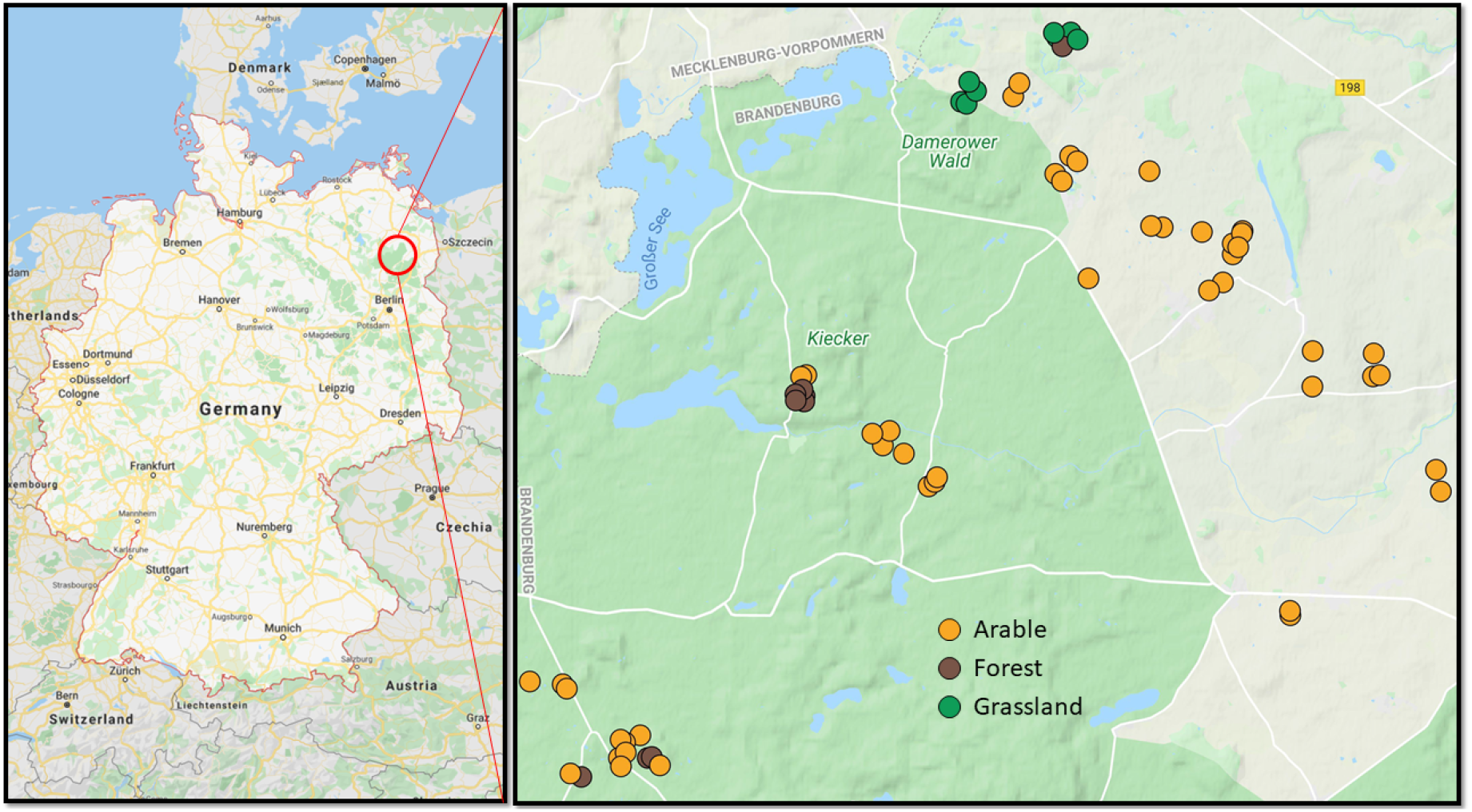
Map showing the location of the sampling area (125 km^2^) ca. 100 km north of the city of Berlin, Germany (left panel) and local distribution of three types of sampled kettle holes (KH) in arable fields (n = 47), forests (n = 11) and grassland (n = 9) (right panel). Map generated with Google Maps online tools.

Water samples were collected whenever enough water was present in the KH. Some occasionally fell dry, however (Nitzsche et al. 2017), and thus could not be sampled at all times, particularly in October 2017. To obtain representative samples, total volumes of *ca.* 20 L were collected at 5-15 locations selected within each KH, with the number of individual samples depending on KH size. The samples were pooled in cleaned buckets and 2 L were subsampled in the field, placed in ice chests containing a mixture of ice and table salt to lower the temperature during transport, and subsequently frozen at −80 °C in the laboratory for later eDNA analysis.

Sediment cores were taken at three time points (Table S1). In March 2017, sediment samples were collected from 54 of the 67 KH, both wet and dry. In some instances, a dense mat of belowground plant parts prevented sediment coring. Subsequently, sediment cores were only collected from wet KH that had recently dried out, or from previously dry KH that had refilled. Between 3-7 cores were taken per KH, depending on KH size, covering both littoral and central areas. The cores were sectioned into surface (upper 5 cm) and lower (5-20 cm depth) sediment layers to try and separate current benthic community from older resting stages and preserved eDNA. The sections were separately transferred into plastic bags were subsampled (1 g wet weight) for eDNA extraction. Both the complete samples and subsamples were stored at −80 °C for further processing. DNA extractions from multiple cores representing surface or lower sediment layers of a given KH at each sampling date were pooled. A compilation of the collected samples is given in Table S1.

### Analysis of physico-chemical properties

Temperature, conductivity, pH, redox potential, and oxygen concentration and saturation were measured on site during sampling using a multiparameter field probe (HI98194, Hanna Instruments, Vöhringen, Germany). Additional water (1 L) was collected to determine concentrations of nutrients and major ions. These samples were immediately frozen in an ice chest containing crushed ice mixed with table salt (NaCl) and analyzed within 48 h. Water analysis followed German standard methods (DIN 38405, 2018). Ca^2+^, Mg^2+^, K^+^, Na^+^, and total Fe were analyzed by inductively coupled plasma optical emission spectrometry (ICP-iCAP 6300 DUO, ThermoFisher Scientific GmbH, Dreieich, Germany). Br^−^, Cl^−^, NO_3_^−^, NO_2_^−^ and SO_4_^2−^ were analyzed using ion chromatography (882 Compact IC plus, Deutsche Metrohm GmbH & Co. KG, Filderstadt, Germany). Ammonium (NH_4_^+^) and soluble reactive phosphorus (orthophosphate; o-PO_4_^3−^-P) were measured spectrophotometrically (SPECORD 210 plus, Analytik Jena AG, Jena, Germany). Total phosphorus (TP) was measured as soluble reactive phosphorus after microwave digestion (Gallery™ Plus, Microgenics GmbH, Hennigsdorf, Germany). Dissolved organic carbon (DOC), total organic carbon (TOC) and total nitrogen (TN) were determined using an elemental analyzer (TOC-VCPH, Shimadzu Deutschland GmbH, Duisburg, Germany) with chemiluminescence detection. The specific absorption coefficient (SAC) was measured on a spectrophotometer (SPECORD 210 plus, Analytik Jena AG, Germany) as a proxy of dissolved aromatic carbon content (Weishaar et al., 2003). Finally, the SAC:DOC ratio was used as a rough measure of DOC quality.

### DNA extraction

The collected 2-L water samples were sequentially filtered (Nalgene filtration tower; ThermoFisher Scientific, Dreieich, Germany) to prevent clogging The filters used were polycarbonate membrane filters (pore size of 10 and 5 μm), combusted GF/F filters and finally polycarbonate filters with a pore size of 0.2 μm (47 mm diameter of all filters). The GF/F filter was included owing to its charge to capture naked eDNA and DNA released from cells lysed by freezing and thawing. All filters were rinsed twice with 50 mL autoclaved MilliQ water to remove salts, and subsequently flash frozen and stored at −80 °C.

Total (environmental) DNA was extracted from 329 samples consisting of 182 water samples, 75 surface sediment samples (< 5 cm), and 66 deeper sediment (5-20 cm) samples. To prevent analytical biases (Bálint et al. 2018), the different filtered subsamples were extracted in separate, randomly selected batches. DNA was extracted with phenol/chloroform according to a method modified by Nercessian et al. (2005). In brief, a CTAB extraction buffer containing SDS and N-laurylsarcosine was added to the samples together with an equal volume of phenol/chloroform/isoamylalcohol (25:24:1) solution. The samples were subject to bead-beating (FastPrep-24™ 5G Instrument, MP Biomedical, Eschwege, Germany), followed by centrifugation (14,000 *g*), a cleaning step with chloroform, and DNA precipitation with PEG-6000 (Sigma-Aldrich, Taufkirchen, Germany). The precipitated DNA was rinsed with 1 mL of 70% ethanol, dried and dissolved in water. Finally, all extracts from the same sample were pooled and kept at −80 °C till further processing.

### Sequencing

Sequencing was conducted separately for the SSU rRNA gene of *Archaea, Bacteria* and eukaryotes at MrDNA (Shallowater, TX, USA) using the following primers: Arch2A519F (5’ CAG CMG CCG CGG TAA 3’) and Arch1071R (5’ – GGC CAT GCA CCW CCT CTC - 3’) for archaea (Fischer et al., 2016); 341F (5’ CCT ACG GGN GGC WGC AG 3’) and 785R (5’ GAC TAC HVG GGT ATC TAA TCC 3’) for bacteria (Thijs et al. 2017); and Euk1560F (5’ TGG TGC ATG GCC GTT CTT AGT 3’) and Euk2035R (5’ CAT CTA AGG GCA TCA CAG ACC 3’) for eukaryotes (Hardy et al. 2010). The primers were barcoded on the forward primer and used in a 30-cycle PCR using the HotStarTaq Plus Master Mix Kit (Qiagen, Hilden, Germany) under the following conditions: 94 °C for 3 min, followed by 30 cycles at 94 °C for 30 s, 53 °C for 40 s and 72 °C for 1 min, followed by a final elongation step at 72 °C for 5 min. The PCR products were checked in 2% agarose gel to determine success of the amplification and relative band intensity. To ensure high coverage of rare taxa, batches of 20 samples were pooled for each sequencing run in equal proportions based on their molecular weight and DNA concentrations. The PCR products were purified using calibrated Ampure XP beads and then used to prepare an Illumina DNA library. Paired end 2 x 300 bp sequencing was performed on a MiSeq sequencer (Illumina, Inc., San Diego, CA. USA) following the manufacturer’s instructions. Sequence data are available at the NCBI Short Read Archive under project number PRJNA641761.

### Bioinformatic analysis

Paired end reads were merged using BBMerge from the BBMap package (part of JGI tools; https://sourceforge.net/projects/bbmap), after which the joined reads were quality trimmed and demultiplexed using cutadapt (V 1.16) to remove reads of low quality (q > 20) and shorter than 150 nt. Taxonomic annotation was performed for all reads from all samples without clustering based on the SILVA SSU NR99 data base (V132; Quast et al., 2013). This was accomplished by using PhyloFlash (V 3.3 b1; https://github.com/HRGV/phyloFlash; Gruber-Vodicka et al., 2019) and Kraken 2 (Wood et al. 2019). To improve the annotation of eukaryotic taxa, a new database was created consisting of all eukaryotic sequences in the SILVA SSU Parc database (V138). The SILVA Parc database also includes eukaryotic sequences shorter than 900 nucleotides and hence covers a much broader range of species than the SILVA NR99 database. The eukaryotic sequences from all samples were annotated using both PhyloFlash and the SINA aligner (Pruesse et al., 2012; V 1.6; https://github.com/epruesse/SINA) requiring a minimum consensus of 3 sequences for last common ancestor assignments. The resulting annotations were merged according to taxonomic names and presence/absence matrices were generated to account for the qualitative nature of the eDNA method especially when merging data from separate assays (i.e. separately targeting *Archaea*, *Bacteria*, and eukaryotes). Statistical analyses (see next section) using matrices generated by different annotation tools resulted in identical patterns.

### Statistical analysis

Multivariate (NMDS, PCA, CAP, Permanova) and diversity (richness and evenness) analyses were conducted using the Primer6 (V 6.1.1) + Permanova Package (V 1.0.1, Primer-E, Quest Research Limited, Auckland, New Zealand). NMDS was conducted using Bray-Curtis dissimilarity, retaining the ordination with the lowest calculated stress out of 1000 iterations. Permanova was used to test for the effect of land-use type, seasonality (i.e. time of sampling) or both. CAP (Canonical Analysis of Principal coordinates) was used to present the data according to factors found to have a significant effect by Permanova. Distance-Based Linear Models with Redundancy Analysis (DBLM-RDA) were used to test for the effects of water chemistry on community structure. Univariate analyses (ANOVA, Kruskal-Wallis and Dunn’s test), and diversity indices (Chao I, taxa richness, evenness) were calculated using the PAST3 software (Hammer et al. 2009). Since the data available in the sequence databases (e.g. SILVA) are not of uniform quality, not all sequences could be annotated to the same taxonomic resolution. Therefore, richness was assessed using the highest assignable taxonomic resolution

Ternary plots were generated using the *ggtern* package (Hamilton and Ferry 2018) in R V3.5 (The R Core Team 2018). Indicator species analysis was performed using the *indicspecies* R package (V.1.7.8; Cáceres and Legendre, 2009) testing for the IndVal index, as well as Pearson’s phi coefficient of association (Chytrý et al. 2002). The latter was used both on presence/absence data and sequence frequencies while considering the appropriate functions and required corrections as outlined in the *indicspecies* package manual (ver. 1.7.8). Indicator species analysis was conducted using the most elaborate annotation matrix (containing 50,000 taxa across the 3 domains *Archaea*, *Bacteria,* and eukaryotes). Additionally, the analysis was corrected for the greater number of sites in arable fields than in grasslands and forests. Data for ternary plots was generated as the percent presence of a specific taxon in each land-use group.

Rarefaction curves and cross-sample species accumulation curves were calculated using the specaccum function in the R Vegan library (Oksanen et al. 2006) and Primer6 (V 6.1.1, Primer-E, Quest Research Limited, Auckland, New Zealand), respectively.

## Results

### Water chemistry

Physical and chemical properties of the water (Fig. 2 and Table S2) highlight the temporal variability of parameters within kettle holes (KH) throughout the study. Variation among samples per land-use type as well as combined sampling campaign and land-use type are shown in Figs S1 and S2, respectively, along with information on statistically significant differences. Water chemistry of the surveyed KH varied among sampling dates and both within and among land-use types, but systematic differences among land-use types were small. Only KH surrounded by forest significantly differed from KH in arable fields and grasslands, and that only in some parameters such as conductivity and concentrations of DOC and most ions, but not nutrients (Fig. S1). In contrast, water chemistry and other environmental parameters did not differ between KH in arable fields and grasslands, even when data from different sampling campaigns were analyzed separately, the only exception being temperature (Fig. S2). The extent of seasonal variability and the timing when parameter-specific maxima or minima were observed differs among individual KH (Fig. 2A). Principle component analysis based on the maximal number of available parameters for the largest possible number of samples (143 of 182 water samples) did not separate KH according to land-use type (Fig. 2B). However, a seasonal pattern emerged between spring (March) and summer (June) with the smaller subset of autumn (October) samples being closer to spring samples, primarily driven by temperature, O_2_ saturation, DOC and nutrient concentrations (Fig. 2C). Conductivity and concentrations of several ions also significantly influenced the ordination, but reflected neither seasonality nor land-use type. All KH would be classified as eutrophic to hypereutrophic based on water chemistry data (Wetzel 2001). However, it is difficult to fully determine the trophic state of the KH in this study for two main reasons. First, no obvious relation was observed between O_2_ saturation and nutrient load at the time of sampling, except for TP in March and May (Fig. S3). Second, primary and secondary productivity, an essential part of measuring eutrophication (Khan and Ansari 2005) were not determined in this study.

**Figure 2.**
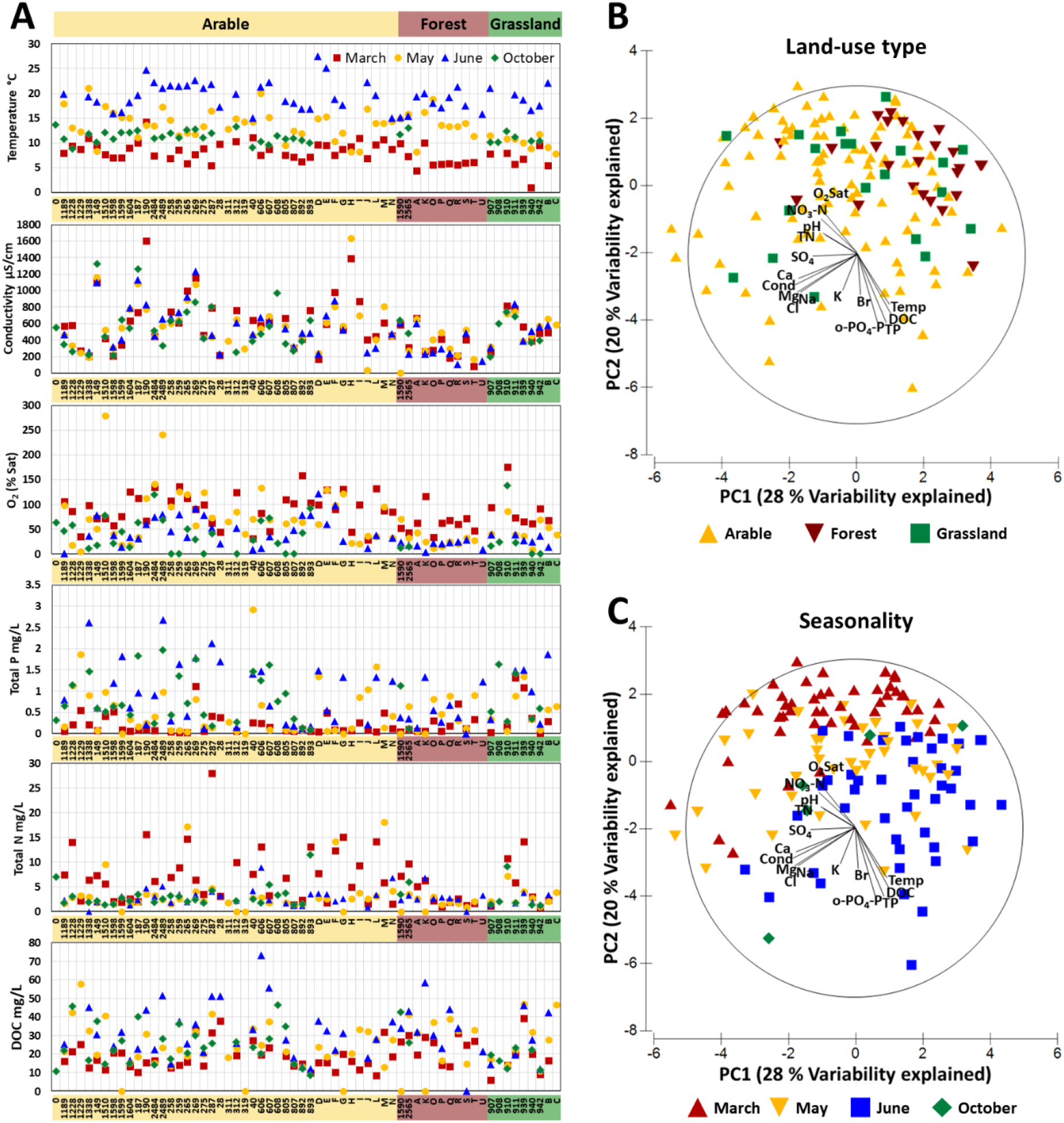
Variability among and within KH in terms of major physical and chemical variables determined during 5 sampling campaigns (A). Principle component analysis of water chemistry data. with samples labeled according to land-use type (B) or month of sampling (seasonality) (C).

### Sequencing effort

Separate sequencing assays resulted in 8.35×10^7^ archaeal, 11.6×10^7^ bacterial, and 11.4×10^7^ eukaryotic SSU rRNA gene sequences per assay, averaging 3.24×10^6^ sequences per sample (Table S3). Reads of eukaryotes were assigned to a large number of taxa (Supplementary dataset 1), including worms, mollusks, arthropods, amphibians, fish, birds and mammals, some of which were evidently rare or occasionally present in the KH.

### α-diversity

The Chao I index shows that overall organismal diversity was greater in sediments than in water. No significant differences among land-use types were observed in either sediment (Fig. 3A) or water (Fig. 3B) when the data were grouped according to land-use type (Fig. 3A). This holds when the data were analyzed together across all sampling campaigns and also when each campaign was inspected separately.

**Figure 3.**
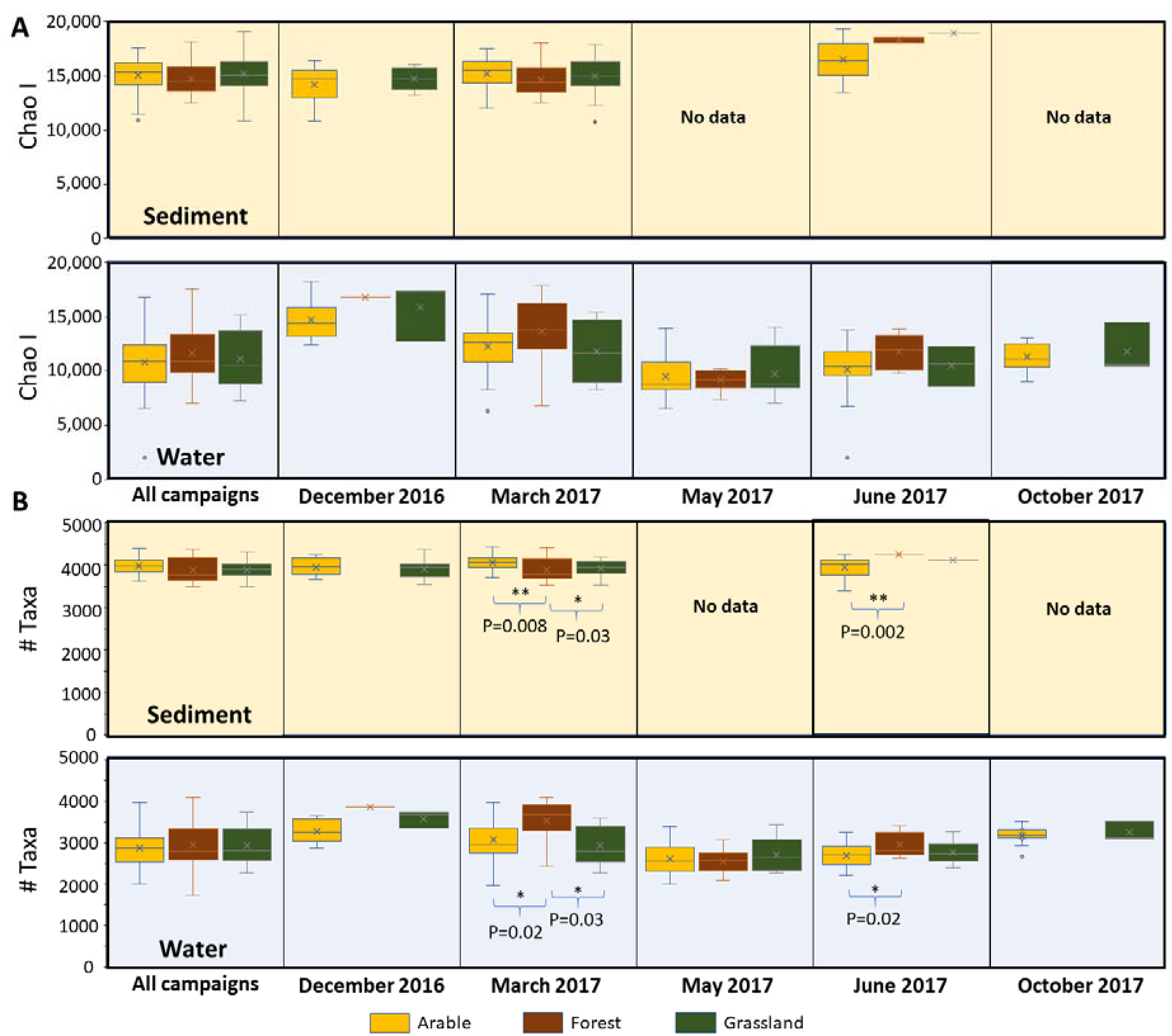
Richness assessment based on Chao I index accounting for the number of taxa for which singleton and doubleton sequences were obtained (A) and taxonomic richness which considers presence / absence alone (B). Whiskers mark the 25^th^ and 75^th^ percentile. Samples are grouped according to the assigned land use type and include *Archaea*, *Bacteria,* and eukaryotes data. Sediment and water samples are separated for both indices. In both cases sequences were grouped according to taxonomic annotations and were not clustered into distance-based operational taxonomic units. As not all sequences could be resolved to the same taxonomic depth (i.e. order, family, genus, species), these indices likely underestimate the true diversity. ANOVA and Kruskal – Wallis tests showed no overall difference between the land use types. However, when pairs of groups were compared using Mann-Whitney’s and Dunn’s tests significant differences were found as marked in the figure.

Depending on the annotation tool used to analyze the data, a total of 13,000 to 50,000 taxonomic entities were identified. Despite the large spread, trends similar to those shown in Fig. 3 were observed in all cases. Therefore, we chose a more stringent analysis which provided less taxonomic resolution by grouping sequences into broad taxonomic groups (e.g. into families rather than genera or genera rather than species). No statistically significant difference in taxon richness was observed among land-use types when all water or sediment samples were analyzed together (Fig. 3B). When analyzed according to sampling period, forest KH water samples collected in March 2017 harbored a higher diversity than samples from KH in grassland or arable fields (Fig. 3B). In contrast, sediment samples collected in March from KH in arable fields harbored more taxa than forest and grassland samples collected at the same time. Samples taken in June show a higher diversity in sediments of forest KH than in those of arable fields, and a similar pattern is apparent in the matching water samples. However, sediment data from the forest KH might be biased because the number of samples was low.

Both taxonomic richness and Chao I show that alpha diversity in water samples across all land-use types was higher in winter and early spring (i.e. December and March), reaching a minimum in mid-spring (May) and increasing again towards winter (Fig. 3B).

We further assessed richness at different time points within different functional groups across the different land-use types (Fig. 4). In most of the depicted groups, an increase in apparent richness (i.e. more species detected) was observed in spring (March, May). This increase and peak in richness occurred across all land use types, although at different magnitudes and not always in parallel. The largest number of species per land use was mostly found in arable fields, followed by forest and grasslands (ANOVA p<0.001). However, a comparison of species accumulation curves suggests there was no difference in richness among the land-use types and therefore the evidently higher number of species detected in arable fields is a result of having more sampling sites (Fig. S4). In contrast, forest KH harbored the largest number of different species per KH (ANOVA p=0.02) while KH from arable fields and grasslands were often similar (Mann-Whitney p=0.6).

**Figure 4.**
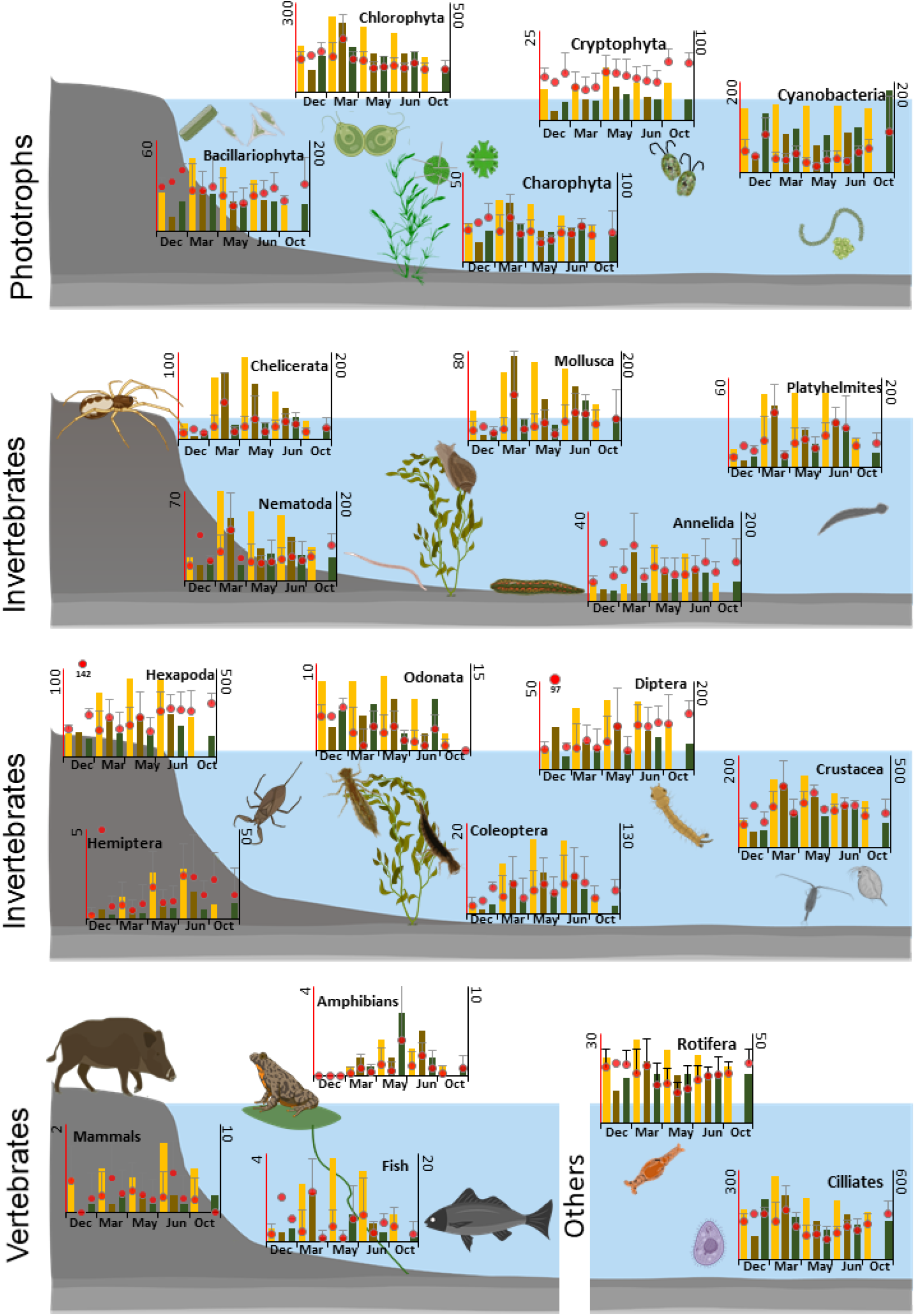
Average apparent species richness (no. of taxa) of selected functional groups per KH from each land-use type for the different sampling periods (left axis, red circles) alongside the summed richness (no. of taxa) per land-use for the same periods (right axis, bars). Yellow, brown, and green bars stand for arable, forest and grassland land use types, respectively. This figure was partially created with BioRender.com.

### β-diversity

Given that eDNA data are non-quantitative across the three domains of life (*Bacteria*, *Archaea*, eukaryotes), specifically with respect to multicellular taxa, the sequence frequency data was converted to presence / absence data. Separate analyses making use of sequence frequency as a proxy for abundance while excluding eukaryotic taxa did not notably alter the presented result (Fig. S5).

NMDS analysis shows a clear separation between community composition of sediment and water samples (Fig. 5A), which accounts for ca. 15 % of the variability across all samples (p=0.001). Water samples were separated according to sampling period (Fig. 5B), explaining *ca.* 11 % of the variability between samples (p=0.001). This percentage increases to 18 % when sequence frequencies are used instead of presence/absence data (p=0.001). In contrast, differentiating the samples according to land-use type shows no distinct pattern (Fig. 5C), although PERMANOVA analysis shows it is statistically significant (p=0.018), explaining only ca. 2 % of the variability between the samples. Breaking down the arable fields into the specific crops (at the time of sampling) did not improve explanatory power. Redundancy analysis using data from the 143 water samples for which all physical and chemical information is available reveals a separation based on sampling time point, similarly to the NMDS analysis, with a clear horizontal separation that appears to be mainly driven by temperature (Fig. 5D). Physical and chemical parameters cumulatively explain 23 % of the total variability among samples, with the contribution of most parameters being significant (p=<0.001 to 0.018), except for concentrations of Cl^−^ (p=0.08) and Br^−^ (p=0.35). Temperature, strongly correlating with the seasonal gradient, has the largest explanatory power among all parameters, accounting for 7 % of the total variability among samples.

**Figure 5.**
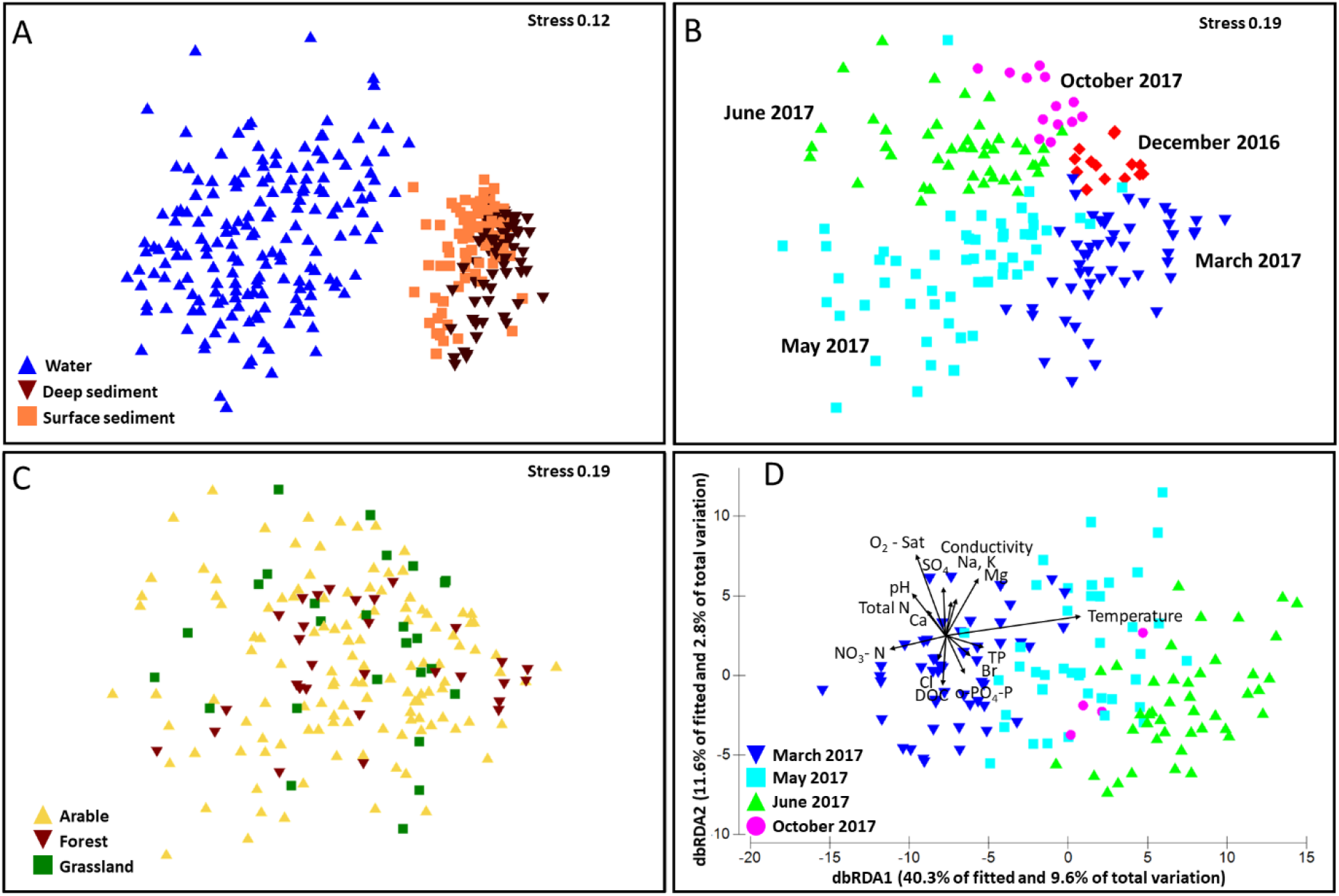
Nonmetric multidimensional scaling (NMDS) of sampled communities revealed a separation between the total community of water and sediment samples (A) and a seasonal clustering of the aquatic communities (B), but no land-use-based ordination patterns (C). A distance-based linear model and redundancy analysis (D) shows a seasonal separation of the water samples alongside the statistically significant environmental parameters, with temperature being the main driver. A three-dimensional NMDS analysis improves the fit, reducing the stress in panels A, B and C to 0.09, 0.12 and 0.12, respectively

To further explore the species distribution based on land-use type, we conducted an indicator species analysis of the water and sediment data using both a presence / absence matrix and sequence frequencies. The patterns were virtually identical (Fig. 6A). No taxa were restricted to a single land-use category. A larger number of taxa were associated with forest or grassland than with arable fields in both sediment and water samples. The top 5 bacterial and eukaryotic associated taxa per land-use type are presented in Table S4 and the complete results are given in Supplementary Dataset 2. In both analyses the number of taxa associated with arable fields was higher in sediment than in water samples. A similar pattern was observed when looking at species associated with two land-use types, one of which is arable fields (Fig. 6A). A graphical representation of taxa associated with different land-use types by means of ternary plots reveals a similar result (Fig. 6B). Most of the taxa appear to be neutral with regard to land-use type, as indicated by the dark blue to red color points clustered in the center of the plots. Only individual taxa, represented by the purple color, spread from the center towards specific land-use types. High correlations, indicated by taxa present in the colored triangles at the vertexes of each plot, are rare. However, overall sediment samples are more inclined towards arable fields than water samples, whereas the latter are more inclined towards forest and grassland. Separate analyses of bacteria (*Bacteria* and *Archaea*) and eukaryotes in sediments show the same distribution pattern. By contrast, eukaryotes in water samples appear to contribute more to the communities associated with forests, and bacteria to that associated with grasslands.

**Figure 6.**
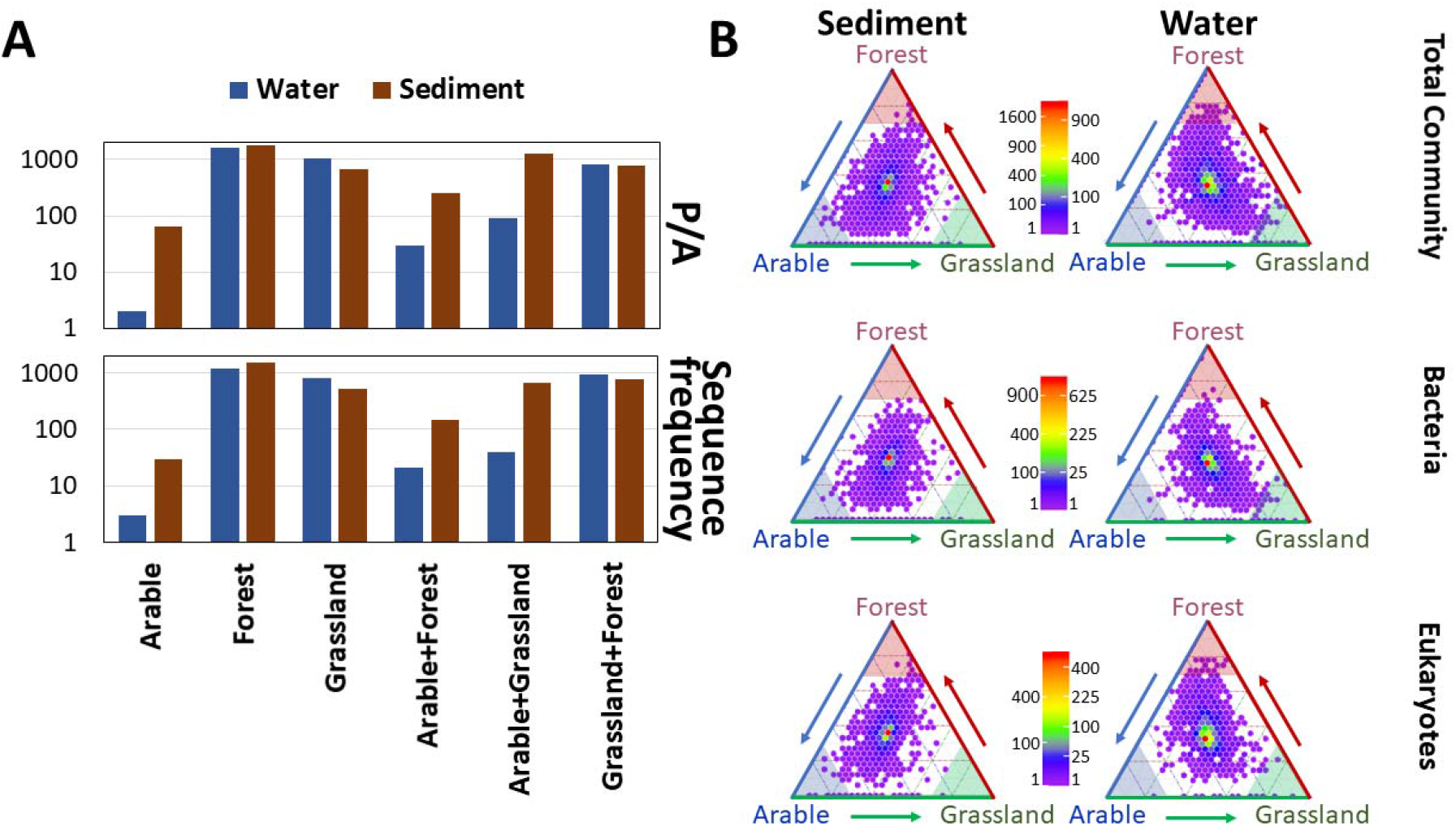
Indicator species analysis of taxa in all water and sediment samples calculated by using either a presence/absence (P/A) matrix as most suitable for eDNA data (upper A panel) or sequence frequency (lower A panel), which is possible because the analysis is based on single taxa. Taxa association with a single land-use type was exclusive as clearly shown in the ternary plots (panel B). The ternary plots depict the association of each taxon to specific land-use types which are represented by the three vertexes of each triangle. Individual taxa are pooled into hexagonal shapes for graphical purposes. The term “*Bacteria*” refers to both *Bacteria* and *Archaea*. The color scale refers to the square root of the number of taxa in each colored point with purple representing single taxa and dark red the maximum number. As indicated by the color code, most taxa appear in the middle of the plot and are hence generalists with respect to land-use type.

We tested whether land-use type influences the distribution of taxa in the entire community or specific taxonomic groups with different trophic functionality (i.e. primary producers and different level consumers), and whether the influence was more pronounced in sediment than in water (Table 1). This was compared to the effect of seasonality which had overall a greater effect on the total community. We defined land-use effect as a significant difference in biodiversity or community structure detectable when data from all sampling campaigns are pooled. For microorganisms we compared presence / absence data and sequence frequencies. For larger multicellular organisms that are unlikely to have been sampled intact, this quantitative data is likely biased because differently-sized body fragments could have been sampled falsely amplifying the amount of DNA without any change in number of organisms. Therefore, the comparisons were limited to presence / absence data. Breaking down the arable-field land-use type into specific crops grown during the sampling period, had no additional explanatory power in any of our analyses.

**Table 1:** Percent contribution of land use and seasonality to the β-diversity in water and sediment samples. Statistically non-significant results (PERMANOVA, p>0.05) are listed as 0 % contribution. Groups marked with an asterisk (*) were not present in all samples. Hemiptera were not detected in enough sediment samples to obtain statistically meaningful results. Quantitative data (i.e. sequence frequency) was only used for microorganisms, which are better represented given our sample size and more likely to have been sampled intact. Therefore, quantitative analysis of the eukaryotic community may be biased. Those percentages are marked with a superscript hashtag (^#^).

| Presence / Absence | Land Use |  | Seasonality |  |
| --- | --- | --- | --- | --- |
|  | Water | Sed. | Water | Sed. |
| **Total Community** | 3 | 4 | 11 | 5 |
| *Bacteria* | 2 | 0 | 9 | 3 |
| **Eukaryotes** | 4 | 5 | 16 | 9 |
| **Cyanobacteria** | 7 | 0 | 15 | 15 |
| **Euk. Phyto.** | 0 | 5 | 16 | 8 |
| **Bacillariophyta** | 0 | 0 | 17 | 13 |
| **Charophyta** | 0 | 0 | 13 | 4 |
| **Chlorophyta** | 0 | 5 | 15 | 6 |
| **Rhodophyta** | 0 | 0 | 12 | 8 |
| **Cryptophyta** | 0 | 7 | 13 | 0 |
| **Other algae** | 0 | 4 | 12 | 5 |
| **Fungi** | 5 | 0 | 16 | 6 |
| **Oomycetes** | 0 | 0 | 7 | 0 |
| **Labyrinthulomycetes** | 0 | 0 | 11 | 0 |
| **Ciliates** | 0 | 0 | 16 | 7 |
| **Rotifera** | 0 | 7 | 10 | 7 |
| **Alveolata** | 0 | 6 | 15 | 6 |
| **Rhizaria** | 0 | 0 | 12 | 0 |
| **Amoeba** | 0 | 9 | 17 | 9 |
| **Het. Flagellates** | 0 | 0 | 0 | 7 |
| **Insecta** | 6 | 0 | 26 | 17 |
| **Hexapoda** | 6 | 0 | 26 | 16 |
| **Odonata\*** | 10 | 23 | 35 | 0 |
| **Diptera** | 6 | 0 | 26 | 16 |
| **Hemiptera\*** | 10 | NA | 28 | NA |
| **Coleoptera** | 0 | 0 | 21 | 13 |
| **Chelicerata** | 0 | 0 | 21 | 16 |
| **Crustacea** | 6 | 0 | 13 | 9 |
| **Myriapoda** | NA | 0 | NA | 0 |
| **Annelida** | 0 | 0 | 17 | 9 |
| **Nematoda** | 0 | 7 | 19 | 19 |
| **Platyhelminthes** | 5 | 0 | 24 | 14 |
| **Mollusca** | 7 | 0 | 14 | 11 |
| **Porifera** | 0 | 0 | 7 | 0 |
| **Other Euk.** | 3 | 7 | 15 | 8 |

**Table 1:** Percent contribution of land use and seasonality to the β-diversity in water and sediment 432 samples.
| Abundance | Land Use |  | Seasonality |  |
| --- | --- | --- | --- | --- |
|  | Water | Sed. | Water | Sed. |
| **Total Community** | 6<sup>#</sup> | 10<sup>#</sup> | 17<sup>#</sup> | 8<sup>#</sup> |
| *Bacteria* | 5 | 0 | 16 | 8 |
| **Eukaryotes** | 6<sup>#</sup> | 14<sup>#</sup> | 20<sup>#</sup> | 10<sup>#</sup> |
| **Cyanobacteria** | 11 | 11 | 17 | 18 |
| **Euk. Phyto.** | 7 | 15 | 20 | 10 |
| **Bacillariophyta** | 0 | 0 | 17 | 17 |
| **Charophyta** | 0 | 11 | 18 | 8 |
| **Chlorophyta** | 0 | 17 | 20 | 8 |
| **Rhodophyta** | 0 | 0 | 10 | 0 |
| **Cryptophyta** | 8 | 14 | 18 | 0 |
| **Other algae** | 5 | 17 | 15 | 5 |
| **Fungi** | 7 | 15 | 18 | 7 |
| **Oomycetes** | 0 | 12 | 10 | 5 |
| **Labyrinthulomycetes** | 0 | 15 | 10 | 0 |
| **Ciliates** | 5 | 16 | 20 | 8 |
| **Rotifera** | 7 | 20 | 15 | 9 |
| **Alveolata** | 6 | 15 | 19 | 0 |
| **Rhizaria** | 0 | 14 | 16 | 0 |
| **Amoeba** | 11 | 17 | 16 | 11 |
| **Het. Flagellates** | 0 | 0 | 12 | 15 |
| **Insecta** | NA | NA | NA | NA |
| **Hexapoda** | NA | NA | NA | NA |
| **Odonata\*** | NA | NA | NA | NA |
| **Diptera** | NA | NA | NA | NA |
| **Hemiptera\*** | NA | NA | NA | NA |
| **Coleoptera** | NA | NA | NA | NA |
| **Chelicerata** | NA | NA | NA | NA |
| **Crustacea** | NA | NA | NA | NA |
| **Myriapoda** | NA | NA | NA | NA |
| **Annelida** | NA | NA | NA | NA |
| **Nematoda** | NA | NA | NA | NA |
| **Platyhelminthes** | NA | NA | NA | NA |
| **Mollusca** | NA | NA | NA | NA |
| **Porifera** | NA | NA | NA | NA |
| **Other Euk.** | NA | NA | NA | NA |

Land-use type had a minimal effect on the whole community, which was slightly larger in the sediment samples (Table 1). In contrast, seasonality had a much larger effect on the community as a whole and also for *Archaea*, *Bacteria* and eukaryotes separately, explaining 11, 9 and 16 % of the variability in taxa composition in the water samples, respectively. Accounting for the sequence frequencies increased the effects of both seasonality and land-use type for bacteria (from 9 to 16 % and 2 to 5 %, respectively). A similar effect was obtained for sequence frequencies of eukaryotes (from 4 to 6 % for land-use type and from 5 to 20 % for seasonality); however, because of possible differences in the representation of multicellular organisms in different samples, this data should be interpreted with caution.

For both total eukaryotic phytoplankton and *Cyanobacteria*, the same pattern was observed as for the total community. Land-use type had a stronger influence on biodiversity in the sediment, and seasonality on biodiversity in the water samples. Nevertheless, when separating the eukaryotic phytoplankton into taxonomic groups, land-use type had a stronger effect on *Chlorophyta* and *Charophyta* detected in the water column. The latter group was dominated by filamentous or single-celled planktonic species from the orders *Klebsormidiales*, *Desmidiales*, and *Zygnematales*. Land-use type had no effect on *Cryptophyta, Bacillariophyta* and *Rhodophyta*. Accounting for sequence frequency, significantly increased the percent of variability explained by land-use for eukaryotic phytoplankton in sediment samples and in some cases also in water samples. Land-use type had no effect on diatoms (*Bacillariophyta*) or *Rhodophyta* regardless of whether presence/absence data or sequence frequencies are analyzed.

Similarly, accounting for abundance, increases the percent variability explained by land-use type for other eukaryotic microorganisms such as fungi, *Oomycota*, and *Rotifera*, but not for heterotrophic flagellates. Overall, with few exceptions, seasonality remained the main explanatory factor of aquatic taxa diversity, while land-use explains best the diversity in sediments.

The species composition of larger multicellular organisms such as insects and major sub-phyla within crustaceans and molluscs is mainly explained by seasonality with the variability of only some of the groups partially explained by land-use type. Among these, the Odonata (dragonflies and damselflies) stand out with 23 % of the variability in taxonomic composition in sediment samples being explained by land-use type.

Using *Crustacea*, *Cyanobacteria,* and eukaryotic phytoplankton we evaluated whether the organisms in the sediment are of planktonic or benthic origin (Fig. 7A-C) and to what extent the sediment and water communities differed from one another. Planktonic copepods (*Calanoida* and *Cyclopida*) and benthic copepods (*Harpacticoida*) as well as ostracods (Podocopida) were present in the deep and shallow sediment as well as in the water. However, the crustacean community structure differed significantly between the water and sediment and between sediment samples from KH surrounded by different land-use types (Fig. 7A). Similarly, despite their dependence on light for photosynthesis, *Cyanobacteria* (Fig. 7B) and eukaryotic phytoplankton (Fig. 7C) occurred not only in water samples but also in sediments. The water and sediment communities differed from one another in both cases. However, while for *Cyanobacteria* land-use type clearly separated the sediment sample, eukaryotic phytoplankton communities from arable fields and grassland sediments were similar. Interestingly, eukaryotic phytoplankton groups from deep and shallow sediments are separated (Fig. 7C).

**Figure 7.**
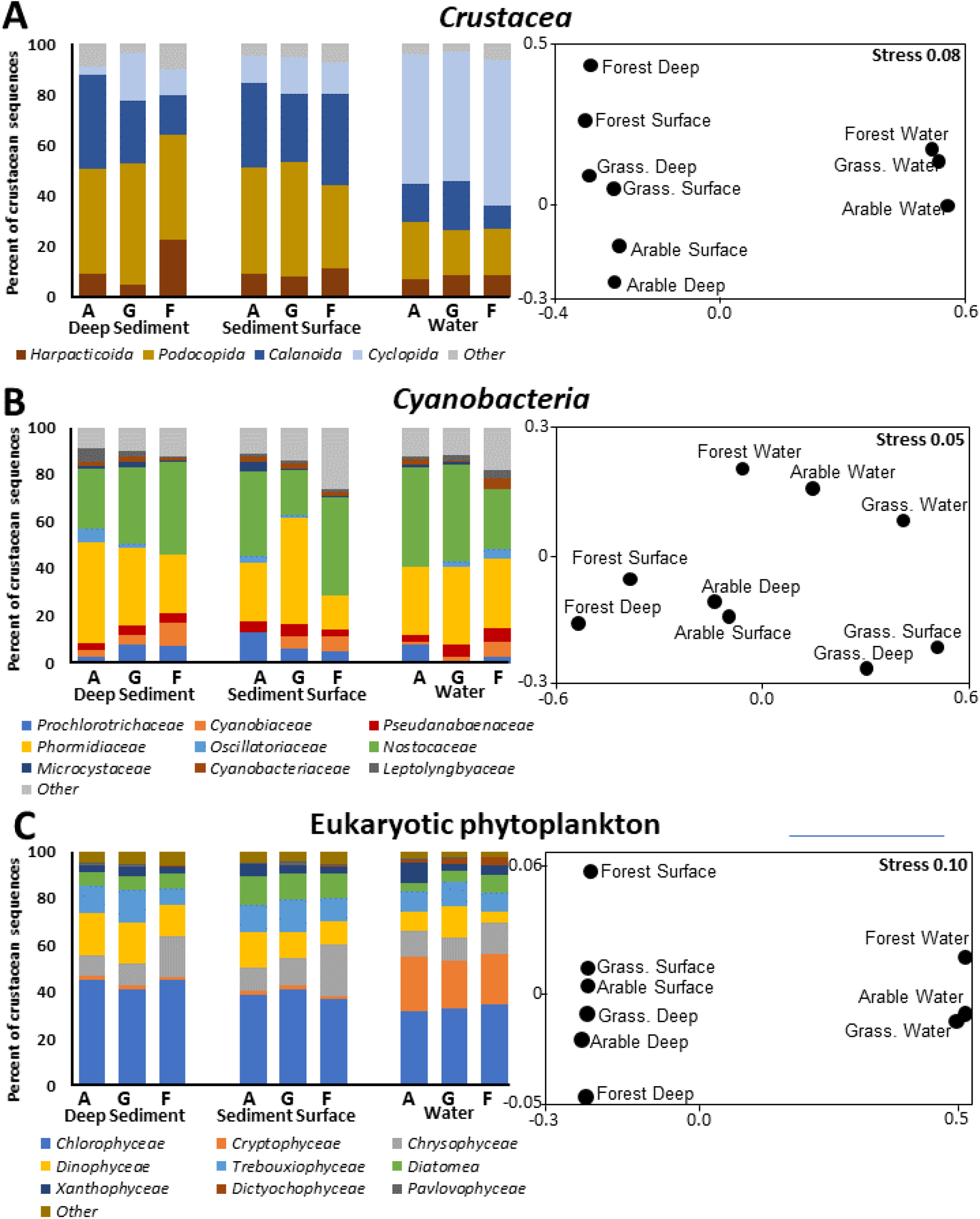
General community composition of crustaceans (A), *Cyanobacteria* (B), and eukaryotic phytoplankton (C) in deep sediments (5-15 cm), surface sediments (0-5 cm) and water samples. The NMDS figures for each group show projections of similarities among the different sample types. Land-use types: A = arable fields, G = grassland, F = forest.

**Figure 8.**
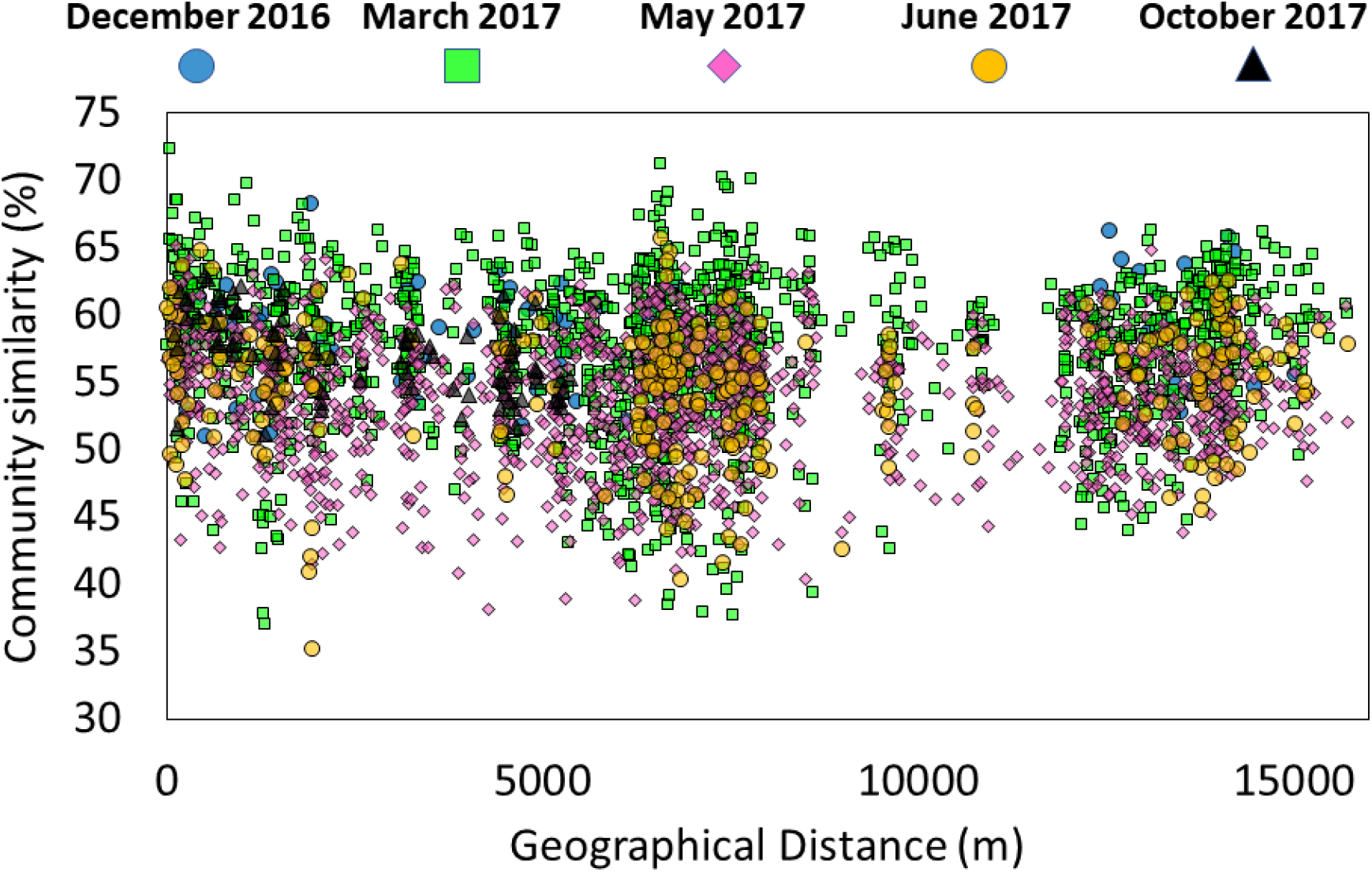
Bray-Curtis similarities between all sample pairs as calculated from a binary (presence absence) taxa matrix and plotted against the geographical distance between the two samples. Plots of the different sampling campaigns are overlaid and distinguished by color. A plot depicting all combinations of sample-pairs and hence accounting for possible lag effects in species dispersal does not reveal any significant correlation (data not shown).

In several cases, adjacent KH were attributed to different land-use types. Therefore, we sought to see if geographical proximity affected the similarity in community composition of KH. In none of the tested cases were KH close to each other (10s of meters apart) more similar than the more distant ones (up to 10 km apart). These results do not change when sequence frequencies were used as proxy instead of presence absence data (data not shown). Second, to verify this observation and to test whether geographical distance affects only certain taxa in our sampling area, a taxon-wise spatial autocorrelation test was conducted. This analysis found no correlation between the distribution of taxa and their geographical location (See section on Spatial Autocorrelation calculations in supplementary material).

## Discussion

In this study we addressed two main questions. First, we sought to evaluate whether a deep-amplicon-sequencing approach of eDNA provides a detailed, nearly complete, snapshot of the biodiversity in small water bodies, such as KH. For this purpose, we used the small subunit of the ribosomal RNA as a general marker, rather than searching for target organisms using taxa specific methods such as specific primers or microarrays (Deiner et al. 2017, Bylemans et al. 2019). Second, by using the above approach, we investigated the magnitude of the effect land-use type in the surroundings of small water bodies has on aquatic biodiversity.

### Deep sequencing of eDNA

Broad-target amplicon sequencing has been used for biodiversity studies for nearly four decades with ever-evolving taxa coverage, in particular as evolving databases allow for better design of new primers and sequencing depth increases as technology evolves. Therefore, we chose to couple this established approach with methods for capturing rare and naked DNA as utilized in eDNA studies. At the same time, we used a separate deep sequencing approach for each of the 3 domains: *Archaea*, *Bacteria*, and eukaryotes to improve the assay specificity and the chances of recovering rare taxa within each domain.

Our analysis focused on taxonomic entities and did not account for microdiversity (i.e. strain variability in marker gene sequence) as can be resolved by defining amplicon sequence variants. This choice, following the approach of the Silva NGS analysis pipeline (Ionescu et al. 2012), considers identical taxonomic entities as likely to have identical or similar functionality, though in the case of microorganisms these entities may be represent ecotypes coming from two adjacent yet separate microniches within one KH. Rarefaction curves separately calculated for each sample and for *Bacteria*, *Archaea* and eukaryotes (Fig. S6) show that more than 50 % of the total discovered taxa per sample were discovered in the first 25 % of the sequences and a clear decrease in discovery rate was observed already before. Given the sequencing depth and the large sample volume, it is not surprising that despite this decrease, new taxa were continuously discovered without apparently approaching an asymptote (Huber et al. 2007, Dethlefsen et al. 2008). A large portion of the reads is, therefore, due to the discovery of relatively rare taxa contributing to more than 50 % of the overall number of discovered taxa. Accordingly, following the rarefaction curve criteria defined in Schöler et al. (2017), our data are sufficient to cover most of the diversity. Sample-wise taxon accumulation plots show that 75 % of the total number of the observed taxa was represented by less than 25 % of the samples (Fig. S7), supporting the notion that overall diversity was well covered. Alternative taxonomic annotation pipelines (e.g. Kraken2) resulted in lower taxonomic diversity, i.e. sequences attributed to different organisms in the presented annotations are merged into single taxa by those alternative methods. Therefore, the results of our rarefaction analyses represent an upper boundary and perhaps an overestimation of taxonomic diversity, suggesting that the sequencing depth we used had even a greater coverage.

The bacterial and archaeal community composition is not informative regarding the coverage of rare species and overall diversity. This is due to the high abundance of these tiny cells in water, typically ranging between 10^5^ and 10^8^ mL^−1^ (Bižić-Ionescu et al. 2015) and the large volume of water concentrated for the sequence analyses. In contrast, our samples likely contained mostly microscopic eukaryotes as intact organisms, while larger taxa can be partly derived from decomposing cells and naked DNA in the water. Therefore, the discovery of multicellular taxa such as plants, insects, amphibians, fish, birds and mammals, whose DNA is expected to be rare in the volume sampled, demonstrates the success in capturing the nature of most permanent, and some transient, organisms in the specific water body. This is in line with the taxa-independent rarefaction curves discussed above, suggesting that most of the diversity in the collected samples was captured. The presence of vertebrates such as fish, birds and mammals could be confirmed either by direct observations or recent tracks on the KH shores, whereas the diversity of benthic macroinvertebrates and rotifers matches or exceeds those observed in parallel surveys using classical microscopic methods (G. Onandia and C. Musseau, unpublished data). This is consistent with previous studies. Although the taxonomic annotation of sequences largely depends on the quality and comprehensiveness of the databases used, the diversity coverage is independent. Accordingly, it has been repeatedly shown that the species detection and sensitivity of eDNA-based studies exceeds that of classical methods (Deiner et al. 2017, Emilson et al. 2017, Fernández et al. 2018, Kim et al. 2019, Yang and Zhang 2020).

We therefore conclude that our deep eDNA amplicon sequencing approach of general marker genes can capture the overall (but not absolute) biodiversity across the domains of life in small water bodies, providing a reliable qualitative overview of resident and transient organisms, including resting stages in the sediment. Nevertheless, even samples for which *ca.* 3 million reads were obtained, the coverage of the taxonomic diversity did not reach a plateau. Accordingly, the coverage obtained in the present study would be too low to analyze microdiversity. Studies targeting specific taxa or a single taxon will benefit from a more targeted approach using designated primers for one or more genes. However, deep sequencing of eDNA marker genes provides a reliable, rapid, and cost-effective method when a detailed cross-taxa overview and total-biodiversity assessment is desired, for instance in surveys motivated by conservation efforts.

### Land-use effects on biodiversity in water

Land use is expected to affect the composition of both permanent and transient members of aquatic communities. For example, intensive agriculture has been shown to result in the decrease in plant (Meyer et al. 2013, Altenfelder et al. 2014), bird (Donald et al. 2006), invertebrates (Wilson et al. 1999) and amphibian (Berger et al. 2011) diversity. Similarly, differences in communities have been documented between ponds in urban vs. rural environments (Joniak et al. 2007, Akasaka et al. 2010) and between lotic waters in forested and agricultural landscapes (Fasching et al. 2020). We tested for land-use effects on α- and β-diversity in the water column and sediment of the sampled KH. Taxonomic richness, Chao I and rarefaction curves all show sediments to be more diverse than water. As sequencing depth (Fig. S7) and sampled biomass are comparable between sediments and water, this may be a result of more niches being available for microorganisms in sediments, but could also be due to long-term (decades) accumulation of dead organisms and naked DNA. The difference between the sediment and water community is likely driven by the long-term accumulation of DNA from different periods of the KH, the presence of eggs and resting stages, the anoxic nature of submerged sediments selecting for specific organisms and the likely introduction of DNA from terrestrial organisms, in part during dry periods. In contrast, the water column samples merely represent snapshots of the current community.

Richness does not differ between land-use types, neither in sediment nor in water samples. This does not change when taxa are separated into the three domains of life (i.e. *Archaea*, *Bacteria*, eukaryotes; Fig. S8). Some significant differences are apparent when samples of different sampling campaigns are separately analyzed. Specifically, taxonomic richness is higher in forest water samples collected in March and June, suggesting a stronger seasonal than land-use effect. In several cases, forest samples also stand out with respect to environmental parameters (Fig. S1). This is in contrast to grassland and arable fields, which are generally not significantly different from one another. The latter may be a result from weak organismal dispersal barriers in the open land, whereas forest KH could be shielded by an arborous buffer zone. In addition, tree cover also results in reduced evaporation, alters the light regime and provides higher input of organic matter as plant litter. Additionally, land-use type is not a permanent feature with transitions of grasslands to arable fields being more common than the other way around (Nitsch et al. 2012, Serrano et al. 2017).

The effect of seasonality (time of sampling) is further evident in β-diversity where more of the variability between samples can be explained by the sampling period rather than land-use type. The percent variability of the water chemistry that can be explained by seasonality is much lower than that of β-diversity (2.5 % vs. 25 %). Thus, it is unlikely that the seasonality effect on the communities is driven by seasonal changes in chemical parameters. The γ-diversity of different taxa (with similar and different trophic function) (Fig. 4) further shows many groups of organisms that follow seasonal changes in richness. These changes are reflected in an increase in richness in early or late spring, often followed by a decrease in summer and autumn across all three land-use types. The latter could be the result of some organisms, such as insects, with life cycles that include aquatic stages and emergence in (late) spring.

Comparing the percent variability explained by seasonality to that explained by land-use type for the entire community, or for selected taxonomic groups, it becomes evident that seasonality is the main factor determining community composition. This suggests that species can inhabit or pass through (actively or passively) all land-use types according to their annual cycle. However, when inspecting the different distribution patterns of each taxon across the land-use types, it becomes evident that the land-use type influences the relative abundance of specific taxa, i.e. their ability to establish a local population. This is clearly apparent for microorganisms in our dataset, and may also have been the case for larger organisms as suggested by our sediment data (e.g. Crustaceans – Fig. 7B). However, for larger organisms this cannot generally be evaluated without a more targeted approach.

### Land-use effects on biodiversity in sediments

Incorporating sequence frequency in the analysis as a proxy for abundance of microorganisms enhanced the percent of variability explained for some of the analyzed taxonomic groups, specifically when applied to the sediment compartment. This suggests that sediment possess a ‘memory’ which in part document the response of aquatic biodiversity to the intensification of agriculture in the region since the early 1950s (Bauerkämper 2004). A previous analysis of carbon and nitrogen isotopes in relation to changes in land use suggests a long-term effect of agriculture can be traced in sediments of the KH in our study area. eDNA degradation in sediments is significantly slower than in the water column (Harrison et al. 2019, Sakata et al. 2020). The occurrence of planktonic phototrophs (eukaryotic algae and cyanobacteria) in sediment, particularly in deeper layers (>5 cm), is a direct evidence of the resulting accumulation of DNA from past communities in the sediments. This is further supported by DNA of planktonic crustaceans found also in both sediment layers distinguished in our study. This implies that our analysis of eDNA in sediments also reflects the distribution of organisms integrated over the sedimentation period. A corollary of this conclusion is that comparisons of the eDNA of such pelagic taxa between surface-water and sediment samples can inform about past communities and could thus be related to long-term changes in land use or other important environmental factors.

The sedimentation rates previously estimated from two KH that are part of the present study (KH258 and KH807; Kleeberg et al., 2016) correspond to an average age of 15-30 years prior to this study in the upper 5 cm and to 50-100 years at 20 cm depth. These values are in line with previous studies in the area, which concluded that the sedimentation rate increased from 1-2 mm y^−1^ prior to 1960, to 5 mm y^−1^ afterwards (Frielinghaus and Vahrson 1998). However, sedimentation rates determined in similar KH in Poland (Karasiewicz et al. 2014) based on ^14^C dating of organic matter, placed the upper 5 and 20 cm as old as 900 and 1500 years ago, respectively. Meij et al., (2019), while supporting the 100 year range, show large variability among the KH in the study area. Accordingly, depending on the sedimentation rate (and extent of sediment re-working) in the different KH, the increased percent of explained variability in the sediment fraction, either for the total community or for specific taxonomic groups with different trophic functionality can be differently interpreted. In case most KH in this study are characterized by rapid sedimentation rates, the sediment eDNA reflects the response of the KH communities to the agriculture intensification in the area since the 1950s (Sommer et al. 2008). In contrast, a slow sedimentation rate would mean sediments still include periods when low-input agriculture was the main practice in the area. However, several facts point to fast sedimentation as the more likely scenario. First, land-use in the area likely changed considerably over the last centuries (Kaplan et al. 2009, Nicolay et al. 2014). Second, only low-input agriculture was practiced in the area prior to the 1950s (Bauerkämper 2004, Sommer et al. 2008). Last, many of the taxa resulting in differences between the sediment and water are primary producers, conceivably responding to increased inputs of agrochemicals, notably P and N from agriculture (Table 1). Therefore, the sediment eDNA could well reflect community changes triggered by intensified agriculture. The separate clustering of eukaryotic phytoplankton from the upper and lower sediment, further suggests a recent change in communities in the last 20-30 years as dated by Kleeberg et al. (2016). Coupling thin-layer sediment eDNA analysis with sediment dating in a large number of KH are needed to reinforce this tentative conclusion.

### Land-use preferences of taxa

Intensive agricultural land use has been shown to result in biotic homogenization, erasing even subtle patterns resulting from other land-use types within a landscape dominated by arable fields (Smart et al. 2006, Buhk et al. 2017). Indicator species analysis reveals that no single taxon is uniquely associated with a specific land-use, although some taxa show a higher preference to one or two (out of three) land uses (Fig. 6A). As expected, based on the above discussion, this analysis also shows a higher number of taxa in sediment than in water - being associated with a particular land-use type. The lowest number of taxa that was specifically associated with a single land-use type was assigned to arable fields, whereas forests harbored the most, followed closely by grassland. This contrast between arable fields vs. forests and grasslands may suggest that the latter two represent more constant environments. The continuous mechanical processing (e.g. ploughing and harvesting) of fields results in constant morphological restructuring of niches and destruction of macro-and micro gradients, possibly influencing littoral organisms or those moving between the aquatic and terrestrial environments. Crop rotation and subsequent alteration in the type of agrochemicals may however affect the KH water, driving continuous changes in the overall community. The quantitative associations of different taxa to specific land-use types visualized through ternary plots support the results of the indicator species analyses suggesting that aquatic biodiversity was likely homogenized at the regional level during more than half a century of intensive agriculture practice, independent of land use immediately adjacent to the KH.

Opposite tendencies in sediment and water samples are evident in the community as a whole and the separately analyzed communities and in the three domains of life (*Bacteria*, *Archaea* and Eukarya). Sediment samples harbored more taxa with a higher affinity towards arable fields, reflecting likely the archived response to the early days of intensive agriculture in the area. In contrast, taxa in water samples are more inclined towards forest and grassland, suggesting that despite the overall homogenization, some taxa may still find refuge in non-arable areas. Interestingly, bacteria and eukaryotes in water samples exhibit a different pattern. *Bacteria* are more inclined towards grasslands, with *Cyanobacteria* dominating the grassland-associated taxa, and eukaryotes towards forests, with fungi being the largest group of forest-associated eukaryotic taxa. These differences probably reflect light availability in grassland and plant litter inputs in forest KH, respectively.

### Buffering of land-use effects are likely negligible

Our study shows that none of the organisms detected in water samples is associated with a specific land-use type. In addition, the community as a whole was not structured by land-use type. One likely explanation is the ubiquitous and long-lasting eutrophication in our study region resulting from intensive agricultural practices during the last decades (Lischeid et al. 2018). Indeed, the phosphorus concentration found in all ponds throughout the study period corresponds to eutrophic to hypertrophic conditions (Wetzel 2001). Additionally, while some individual chemical parameters did differ among land-use types (e.g. DOC), the latter does not explain the chemical variability among the KH, indicating a chemical homogenization effect. Lischeid et al. (2018) showed that KH in the study area are connected via groundwater. Therefore, the nutrient and mineral concentrations in a given KH may reflect an integral of the inputs in the area rather than a local chemical signature. These physical and chemical properties of the water play a major role in shaping aquatic communities. Therefore, the local biota likely reflects the water it resides in rather than mirrors the surrounding land-use type. Joniak et al. (2007) conducted a large study on zooplankton and macrophytes covering 165 ponds in Poland situated in a similar environmental setting as our KH, but covering a larger eutrophication gradient. Certain taxonomic groups tended to be associated with different eutrophication states of the ponds but like in our study, no correlation was found to land-use type. Vegetated buffer strips at least 5 m in width around the ponds were proposed to minimize or even eliminate the influences of surrounding land cover, in particular nutrient inputs from arable fields. In our area, however, the connection of KH via groundwater could have minimized such buffering effects of surrounding vegetation. This lack of notable land-use influences is consistent with analyses of other rural ponds in different land-use types (Declerck et al. 2006, Joniak et al. 2007, Pätzig et al. 2012), this is in contrast to a pronounced land-use effect on ponds found when comparing urban and rural ponds and lakes (Akasaka et al. 2010, Kraemer et al. 2020).

### Geographical distribution

Organisms belonging to all domains of life are transferred between KH via different abiotic and biotic vectors such as wind, insects, mammals and humans. The dense network and long existence of KH in the region likely presents transfer opportunities to all organisms over time. Choudoir et al. (2017) showed that dispersal area of bacteria lies in the order of thousands of km^2^. On the other end, species with elevated loyalty to birth ponds such as amphibians (Smith and Green 2005), who typically do not wander further than a few hundred meters from “home”, will encounter new ponds within this range due to the high density of KH in the area. Nevertheless, once these organisms arrive at a new location, they may be unsuccessful in establishing a new local population and are therefore eradicated or remain extremely low in abundance or dormant state until an opportunity arises for them to multiply (Ionescu et al., 2014). This may be the case for most bacteria and both unicellular and some multicellular eukaryotes. This explains how in a landscape with homogenized biodiversity where generally “everything is everywhere”, the difference among KH communities is not in par with geographical distance.

## Conclusions

We demonstrate that broad-scale eDNA analyses can serve to identify specific taxonomic groups that may be influenced by land-use change or land management. Such groups can be subsequently investigated in detail through targeted studies at high taxonomic resolution using additional or alternative genetic markers. The enhanced sensitivity of sequence frequencies as opposed to presence/absence data to detect land-use effects on biodiversity in water or sediment suggests that while species composition may not be influenced by land-use, their ability to establish identical populations everywhere may exist. Accordingly, following low-cost, deep sequencing broad eDNA surveys, quantitative studies should be employed to investigate specific conservation-prone taxa.

Our analysis of water and sediment eDNA resulted in opposing magnitudes of the effect of land-use type on the biodiversity of KH in an agricultural landscape. We propose that the higher land-use effect, noticeable in the deeper sediment data for the total community and specific taxonomic groups, likely reflects more than half a century of intensive agriculture. Sediment eDNA data reveals a response in the abundance of primary producers and planktonic consumers, possibly caused by the continuous eutrophication of KH as reflected in KH water chemistry today. This eutrophication process archived in the sediment could have led to the homogenization of biodiversity across the entire study area, resulting in a low detectable land-use effect on biodiversity patterns in the water samples, which represent the current situation. This broad-scale homogenization effect on aquatic biodiversity could have been reinforced by KH connectivity below and above ground. Below ground, groundwater can propagate effects of one land-use type such as arable fields to KH located elsewhere. Above ground, the dense network of KH facilitates tight coupling through active and passive species dispersal.

Our eDNA approach, capturing all taxa across the three domains of life are consistent with previous studies focusing on specific groups. Most KH included in those studies are located in areas with a long history of industrialized agriculture. Although the small size of KH implies a strong influence of the surroundings, we propose that the lack of pronounced land-use effects results from the homogenization, and possibly loss, of KH biodiversity during a long period of intensive agriculture and the concomitant KH eutrophication. Interestingly, similar to our results on KH, some recent lake studies also show minimal effects of land-use type (Abell et al. 2011, Catherine et al. 2016, Marmen et al. 2020).

This conclusion emphasizes the intricacies of aquatic biodiversity conservation in landscapes dominated by agriculture. In a broad survey summarizing the results of different conservation practices, Gonthier et al. (2014) conclude that the combination of decreased management intensity (i.e. less agrochemicals) coupled with increases in landscape complexity around arable fields and farms, is most suitable to prevent species loss. However, the effects of such measures in areas where biodiversity has been already homogenized over half a century is still unclear.

## Supporting information

Supplementary material - Spatial autocorrelation

Table S1 sampling summary

Table S2 Physico-Chemical paramteres

Table S3 Sequence counts

Table S4 Indicator Species

## Acknowledgments

We thank the farmers and landowners in the study area for their collaboration and making the sampled KH accessible. We also thank the Landwirtschaft- und Umweltamt - Landkreis Uckermark for assistance and support in sampling the forest KH in the Kiecker Nature reserve, and Dr. Günter Heise for accompanying us. Further thanks are due to Dr. Masahiro Ryo for analysis of spatial autocorrelation analyses, and to Mr. Gonzalo Idoate Santaolalla, Dr. Jason Woodhouse, Dr. Luca Zoccarato, Mr. Darshan Neubauer, Mr. Alberto Villalba, Mr. Justin Stranz and Mr. Ignaciao Rodanes Ajamil for sampling assistance. This study was funded by the German Federal Ministry of Education and Research (BMBF) within the Collaborative Project “Bridging in Biodiversity Science - BIBS” (funding number 01LC1501). M. Bizic was additionally funded through the DFG Eigene Stelle project (BI 1987/2-1). All responsibilities for the content of this publication are assumed by the authors.

*# This paper has been revised since this version but is still under review and therefore cannot be updated. This original version aims to allow access to the data and discussions presented in: Bizic et al., 2022, Land-use type temporarily affects active pond community structure but not gene expression patterns. Molecular ecology **https://doi.org/10.1111/mec.16348***

