## Supplementary Figures for "From microbes to mammals: pond biodiversity homogenization across different land-use types in an agricultural landscape"

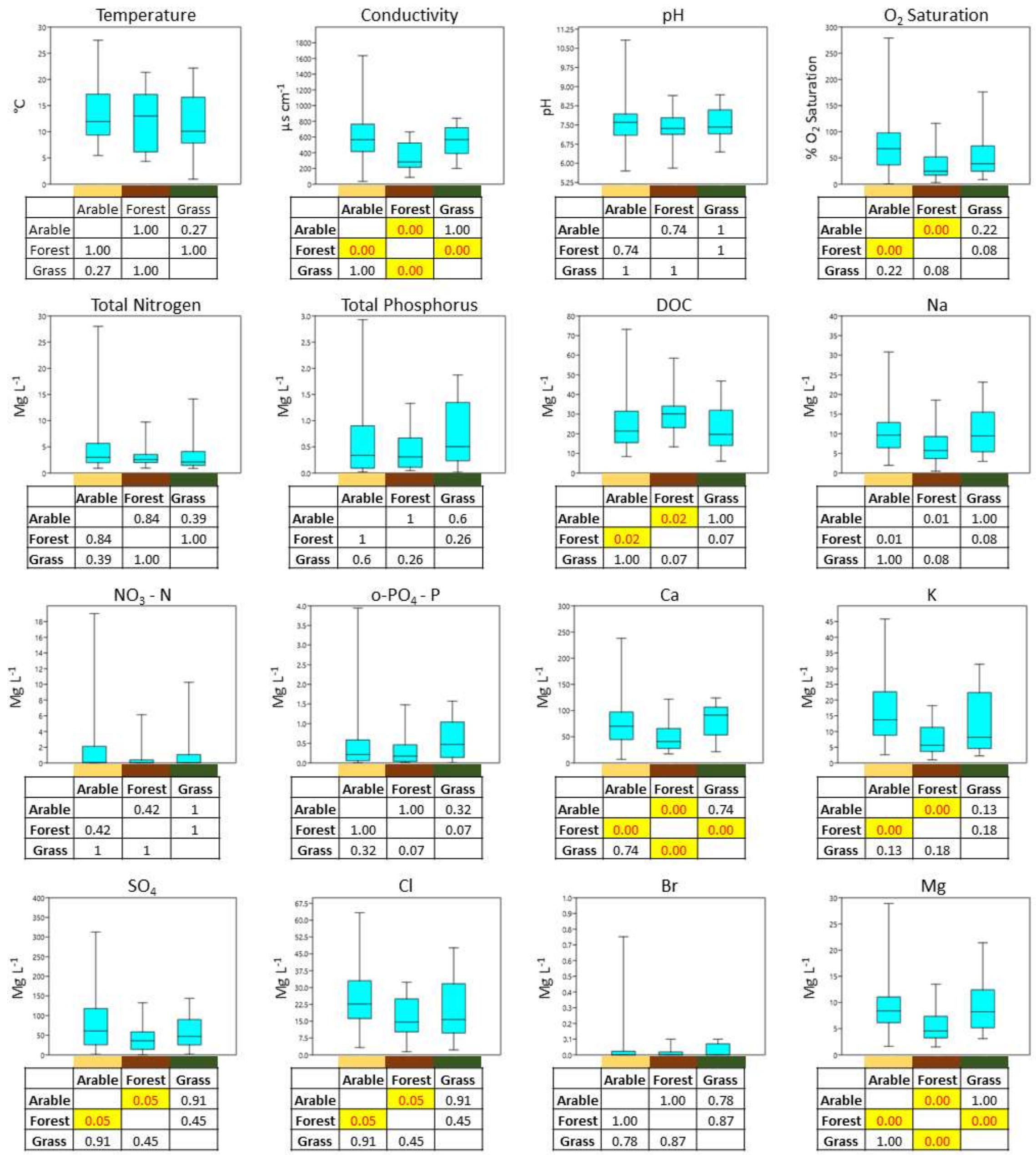


Temperature

| | Arable | Forest | Grass |
|---|---|---|---|
| Arable | | 1.00 | 0.27 |
| Forest | 1.00 | | 1.00 |
| Grass | 0.27 | 1.00 | |

Conductivity

| | Arable | Forest | Grass |
|---|---|---|---|
| Arable | | 0.00 | 1.00 |
| Forest | 0.00 | | 0.00 |
| Grass | 1.00 | 0.00 | |

pH

| | Arable | Forest | Grass |
|---|---|---|---|
| Arable | | 0.74 | 1 |
| Forest | 0.74 | | 1 |
| Grass | 1 | 1 | |

$O_2$ Saturation

| | Arable | Forest | Grass |
|---|---|---|---|
| Arable | | 0.00 | 0.22 |
| Forest | 0.00 | | 0.08 |
| Grass | 0.22 | 0.08 | |

Total Nitrogen

| | Arable | Forest | Grass |
|---|---|---|---|
| Arable | | 0.84 | 0.39 |
| Forest | 0.84 | | 1.00 |
| Grass | 0.39 | 1.00 | |

Total Phosphorus

| | Arable | Forest | Grass |
|---|---|---|---|
| Arable | | 1 | 0.6 |
| Forest | 1 | | 0.26 |
| Grass | 0.6 | 0.26 | |

DOC

| | Arable | Forest | Grass |
|---|---|---|---|
| Arable | | 0.02 | 1.00 |
| Forest | 0.02 | | 0.07 |
| Grass | 1.00 | 0.07 | |

Na

| | Arable | Forest | Grass |
|---|---|---|---|
| Arable | | 0.01 | 1.00 |
| Forest | 0.01 | | 0.08 |
| Grass | 1.00 | 0.08 | |

$NO_3$ - N

| | Arable | Forest | Grass |
|---|---|---|---|
| Arable | | 0.42 | 1 |
| Forest | 0.42 | | 1 |
| Grass | 1 | 1 | |

o-$PO_4$ - P

| | Arable | Forest | Grass |
|---|---|---|---|
| Arable | | 1.00 | 0.32 |
| Forest | 1.00 | | 0.07 |
| Grass | 0.32 | 0.07 | |

Ca

| | Arable | Forest | Grass |
|---|---|---|---|
| Arable | | 0.00 | 0.74 |
| Forest | 0.00 | | 0.00 |
| Grass | 0.74 | 0.00 | |

K

| | Arable | Forest | Grass |
|---|---|---|---|
| Arable | | 0.00 | 0.13 |
| Forest | 0.00 | | 0.18 |
| Grass | 0.13 | 0.18 | |

$SO_4$

| | Arable | Forest | Grass |
|---|---|---|---|
| Arable | | 0.05 | 0.91 |
| Forest | 0.05 | | 0.45 |
| Grass | 0.91 | 0.45 | |

Cl

| | Arable | Forest | Grass |
|---|---|---|---|
| Arable | | 0.05 | 0.91 |
| Forest | 0.05 | | 0.45 |
| Grass | 0.91 | 0.45 | |

Br

| | Arable | Forest | Grass |
|---|---|---|---|
| Arable | | 1.00 | 0.78 |
| Forest | 1.00 | | 0.87 |
| Grass | 0.78 | 0.87 | |

Mg

| | Arable | Forest | Grass |
|---|---|---|---|
| Arable | | 0.00 | 1.00 |
| Forest | 0.00 | | 0.00 |
| Grass | 1.00 | 0.00 | |

**Figure S1.** Distribution of environmental parameters across the different land uses summed up across all the sampling campaigns. Mustard, brown, and green colors represent arable fields, forests, and grassland, respectively. Significant differences between groups of samples according to the Mann-Whitney test and Bonferroni-corrected p-values are highlighted in yellow in the interaction matrix under each panel. P-values lower than 0.01 appear as 0.00.

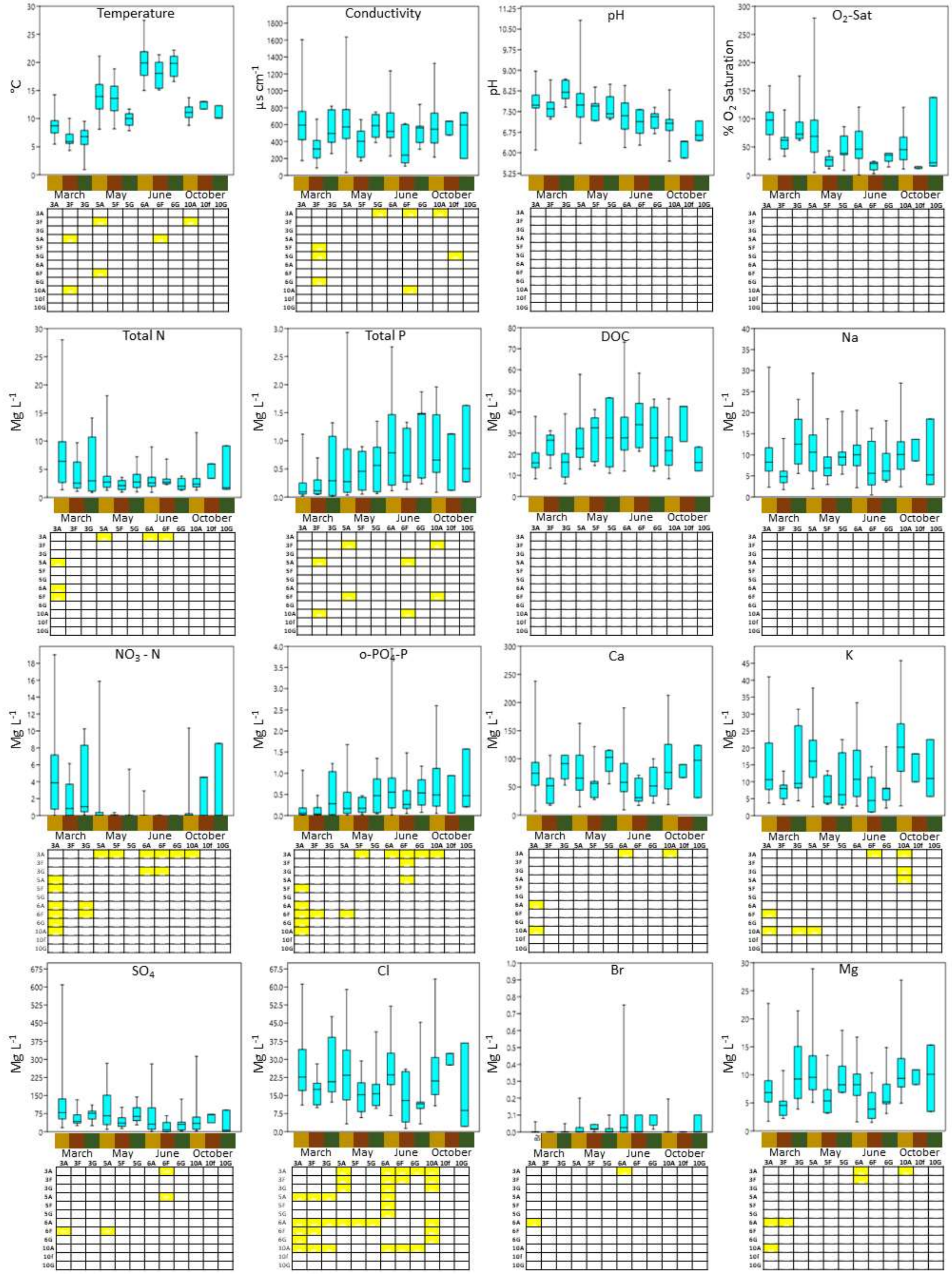


**Figure S2.** Distribution of environmental parameters across the different sampling campaigns and land uses. Mustard, brown, and green colors represent arable fields, forests, and grassland, respectively. Significant differences between groups of samples according to the Mann-Whitney test and Bonferroni-corrected p-values are highlighted in yellow in the interaction matrix under each panel.

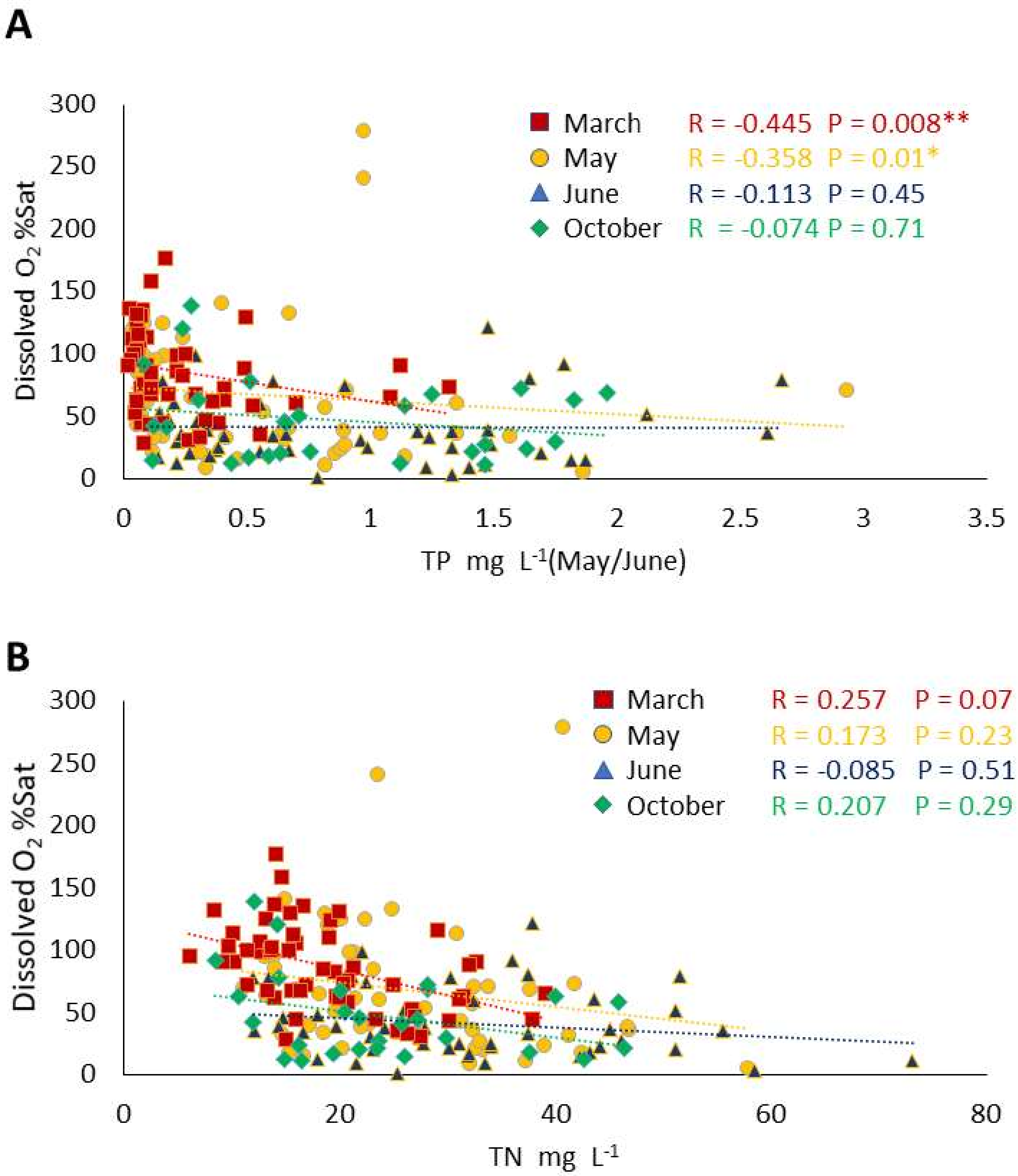


**Figure S3.** Oxygen saturation vs. total phosphorus (A) and total nitrogen (B) from different KH and the different campaigns. Spearman correlation factor (R) and correlation significance are shown in the legend of each plot. A significant decrease in oxygen saturation with increasing nutrients could be indicative of eutrophication.

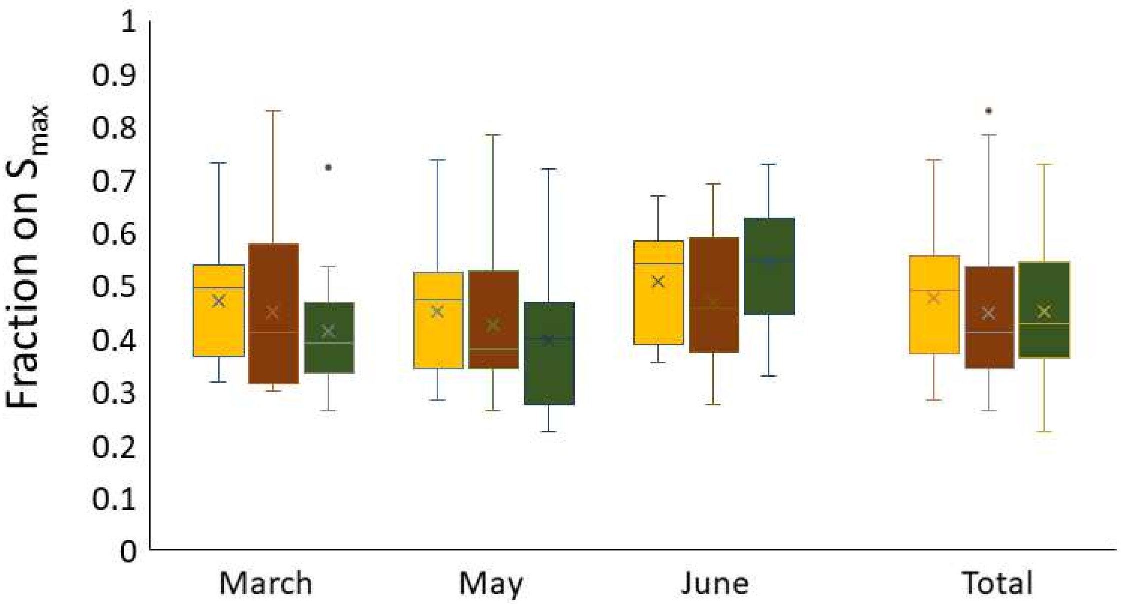


**Figure S4.** Averaged fraction of accumulated species per land use type for each campaign or in total considering the first 6 sites. The fraction was calculated out of the maximum number of species per campaign for the different functional groups detailed in Table 1. No statistically significant different was seen between the different land-use types suggesting that the larger number of taxa detected in arable fields (Fig. 4) is a result of a larger sampling effort (i.e. more sampled KH).

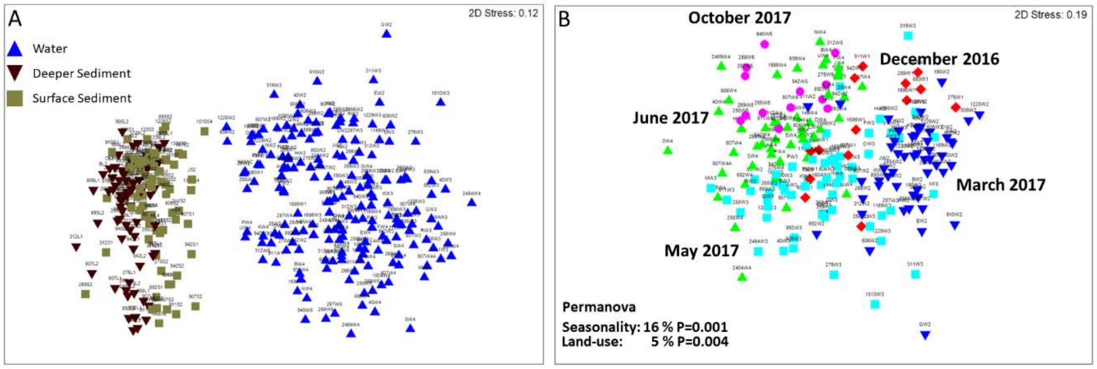


**Figure S5.** NMDS analysis of bacterial and archaeal community from sediment and water samples (A) or just water samples (B) calculated using Bray Curtis similarity and accounting for sequence frequency as a proxy for taxa abundance.

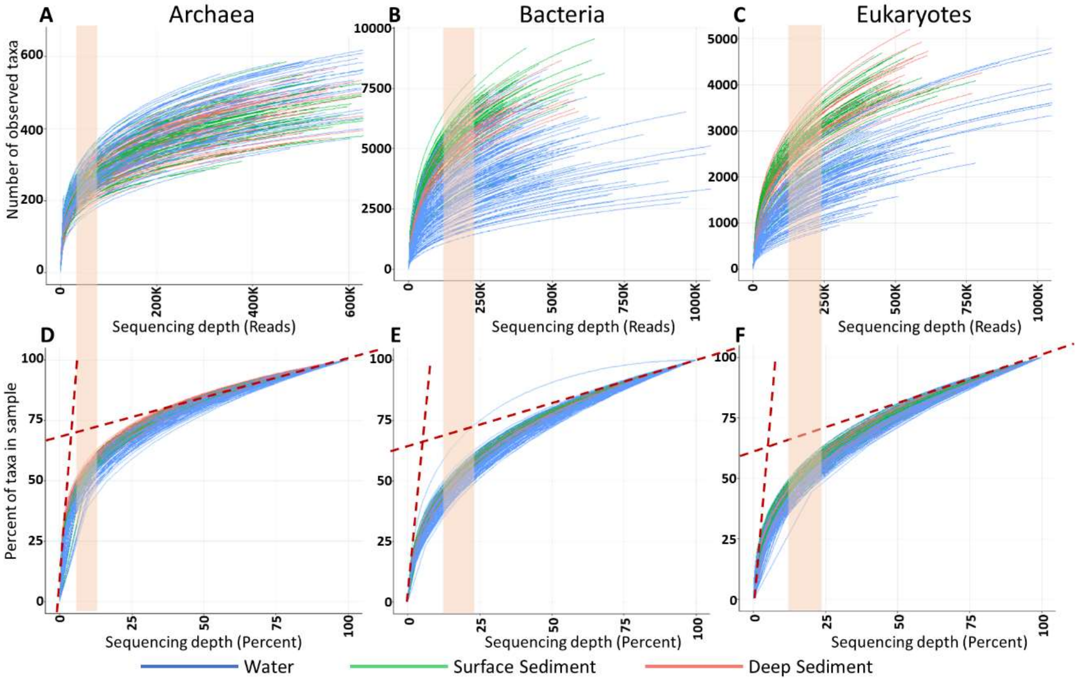


**Figure S6**. General rarefaction curves for *Archaea*, *Bacteria* and eukaryotes calculated for all samples (A-C) alongside the same data normalized to the maximum number of sequences and taxa available in each sample (D-E). The percent/number of sequences needed to discover 50 % of the taxa is highlighted by the colored bar. The triangle enclosed between the two dashed asymptotic lines and the rarefaction curves, show the region in which the number of taxa discovered by increased sequencing significantly decreases.

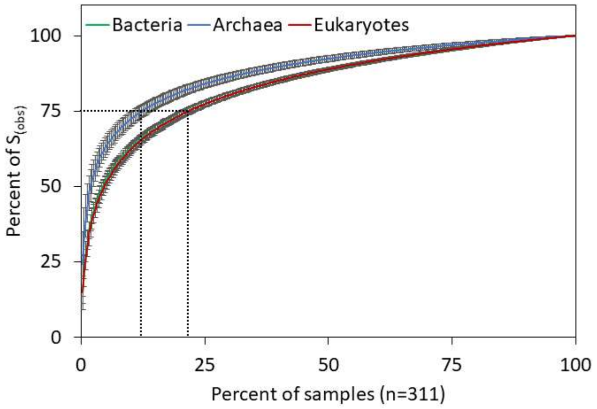


**Figure S7**. Species accumulation curve for *Archaea*, *Bacteria* and Eukaryotes showing that over 75 % of the taxa identified are found in the first 25 % of the samples (< 77 samples).

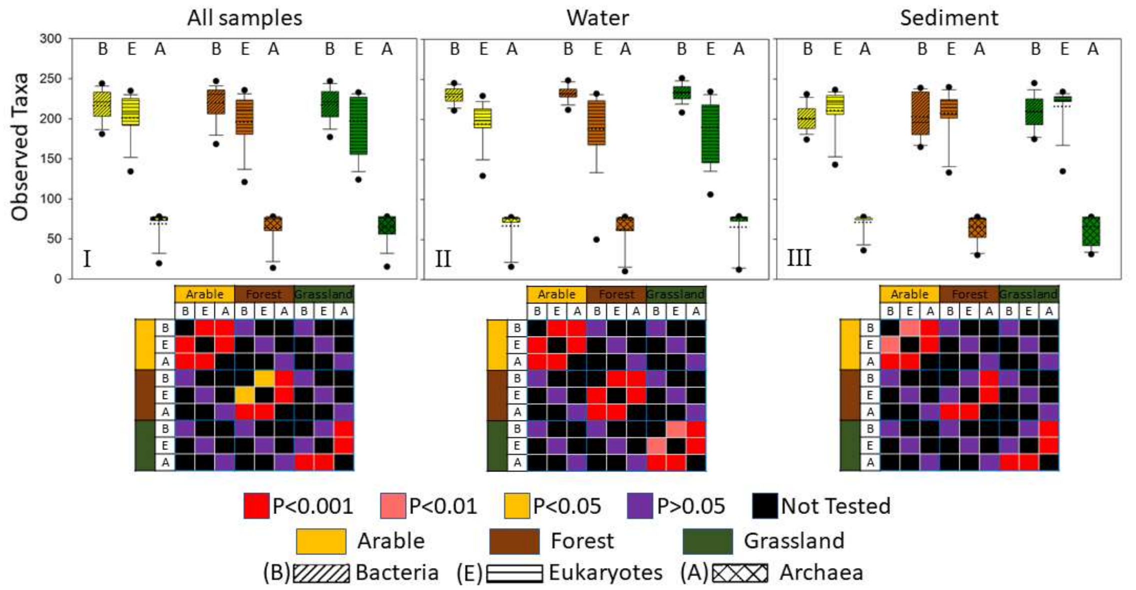


**Figure S8.** Taxonomic richness of the different domains of life, *Bacteria* (B), Eukaryotes (E) and *Archaea* (A) separated into the three types of studied land-uses (Arable, Forest, Grassland) for all samples (I), water samples (II) and sediment samples (III). The significance of differences between different sample groups was tested by ANOVA on Ranks and significant results are marked in the matrices below each panel. Cross land-use-type differences were tested per domain. Cross domain-cross land-use type (i.e. *Bacteria* in arable fields vs. Eukaryotes in forests, were not tested).
