## Supplementary material - Spatial autocorrelation for "From microbes to mammals: pond biodiversity homogenization across different land-use types in an agricultural landscape"

**Spatial autocorrelation scripts and calculations**

***Written by Masahiro Ryo***

***2019 03 03***

---

### Spatial analysis

Testing if any spatial autocorrelation structure appears in the distribution of each taxonomic group.

1. For each taxon, applying distance-based Moran's Eigenvector Mapping (dbMEM)
2. Opposingly, regressing each of the dbMEM patterns with the taxon data

#### 1. Settings
Set the working directory appropriately.

```
r
setwd("C:/Users/masah/Dropbox/i_share/Research_share/2019_Danny_KHmicrobes/R/Dann
y_KH") getwd()
```

```
##                                                                           [1]
"C:/Users/masah/Dropbox/i_share/Research_share/2019_Danny_KHmicrobes/R/Danny_KH"
```

Calling some packages. Packages are automatically installed if needed.

```
r # install devtools tmp.install = require("devtools", character.only =
TRUE)==FALSE if(tmp.install) install.packages("devtools") # install patchwork
tmp.install = require("patchwork", character.only = TRUE)==FALSE if(tmp.install)
devtools::install_github("thomasp85/patchwork") # install and require them
package.list = c("ggplot2", "patchwork", "adespatial", "data.table", "mlr",
"parallelMap") tmp.install = which(lapply(package.list, require, character.only =
TRUE)==FALSE)                                              if(length(tmp.install)>0)
install.packages(package.list[tmp.install],     repos     =     "http://cran.us.r-
project.org") lapply(package.list, require, character.only = TRUE)
```

### 2. Data handling

Let's read the dataset:

```
df = fread("../../Data/BIBS_DNA_kraken_Correct_Taxonomy_USEME.csv")


skip_data = 13
n_taxa = ncol(df) - skip_data
n_site = nrow(df)
```

The number of taxa: 1.352310^{4}.
The number of sites: 323

The data is mostly zero-inflated (many zero data). Visualizing the relationship between the proportion of zero-data and the number of taxa that exceeds the proportion.

```
cut_threshold = 0.9 # threshold for data occupation

n_count = numeric(0)
for(i in seq(0,1, by=0.05)){
  n_count = append(n_count, sum(apply(df[,(skip_data+1):ncol(df)], 2,
function(x) sum(x>0)>=(i*nrow(df)))==TRUE))
}
plot(seq(0,1, by=0.05), n_count/n_taxa, type = "l", xlab="proportion of
non-zero data", ylab="count")
```

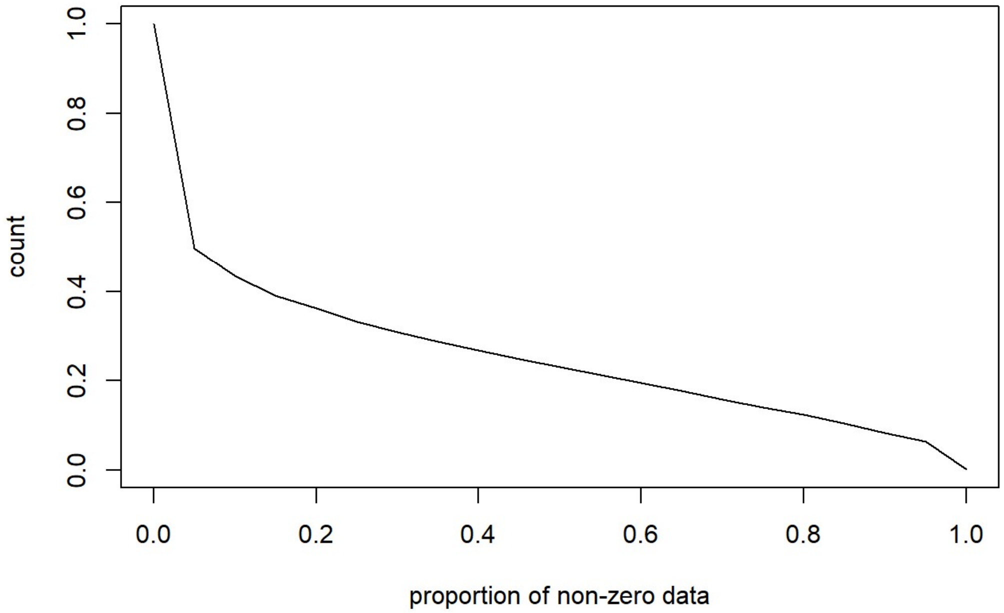


About the half of taxa (6761) are occupied by zero more than 95% of the sampling sites (n = 323).
Here, we discard the taxa having zero-data more than 10% for further analyses.

```
id_selected = which(apply(df[,(skip_data+1):(skip_data+n_taxa)], 2,
function(x) sum(x>0)>=(cut_threshold*nrow(df)))==TRUE) + skip_data

df_selected = df[,..id_selected]
```

**The number of taxa is 1131 after the screening.**

---

### 3. Spatial Autocorrelation Patterns across multiple scales

In this section, using distance-based Moran's Eigenvector Mapping, we generate spatial autocorrelation patterns across multiple scales from the geocoordinates of the sampling sites. MEM vectors are screened according to its power to explain the spatial variability. See Bauman D., Fortin M-J, Drouet T. and Dray S. (2018)
R package: adespatial

```
dbmem.data0 = dbmem(xyORdist=cbind(df$Lon,df$Lat))
dbmem.selected0 = mem.select(x=cbind(df$Lon,df$Lat), listw =
attributes(dbmem.data0)$listw)
## Procedure stopped (R2more criteria): variable 6 explains only 0.000620
of the variance.
print(dbmem.selected0$summary)
##   variables order          R2     R2Cum  AdjR2Cum pvalue
## 1      MEM1     1 0.593704130 0.5937041 0.5924384  0.001
## 2      MEM3     3 0.207731438 0.8014356 0.8001945  0.001
## 3      MEM2     2 0.134851233 0.9362868 0.9356876  0.001
## 4      MEM4     4 0.009006865 0.9452937 0.9446055  0.001
## 5      MEM5     5 0.002362262 0.9476559 0.9468303  0.002
dbmem.selected =
dbmem.selected0$summary$variables[which(dbmem.selected0$summary$R2>0.10)]
```

In total, 6 MEM vectors were produced.
Yet, we focus only on positive autocorrelation patterns of which R2 is higher than 10%.
We use MEM1, MEM3, MEM2 for further analyses.

```
dbmem.data = data.frame(cbind(df$Lon, df$Lat,
dbmem.data0[,dbmem.selected]))
colnames(dbmem.data) = c("Longitude", "Latitude",
colnames(dbmem.data0[,dbmem.selected]))

df = cbind(df, dbmem.data)
```

We plot the geocoordinates of the sampling sites and the selected MEM vectors.

```
# plot MEMs

g_0 = ggplot(data=dbmem.data, aes(x = Longitude, y=Latitude)) +
      geom_point()+
      xlim(c(53.3,53.45)) +
      ggtitle('Sites') +
      theme(text = element_text(size=9),
           axis.text.x = element_text(angle=90, hjust=1))

g = ggplot(data=dbmem.data, aes(x = Longitude, y=Latitude)) +
     scale_color_distiller(palette = "RdYlBu") +
     guides(size=FALSE) +
     xlim(c(53.3,53.45)) +
     theme(
           legend.title=element_blank(),
           legend.text = element_text(size=8),
           legend.key.width = unit(3, "point"),
           text = element_text(size=9),
           axis.text.x = element_text(angle=90, hjust=1),
```

```
             legend.justification = c(0.5, 0.5),
             legend.position = c(1, 0.5),
             legend.background = element_rect(fill=alpha('white', 0.0))
             )
for(i in 1:length(dbmem.selected)){
  eval(parse(text=(paste(
    "g_",i, "= g + geom_point(aes(size = abs(MEM",i,"), color = MEM",i,"))
+ ggtitle('MEM", i,"')",
    sep=""))))
}
print((g_0|g_1) / (g_2 | g_3))
```

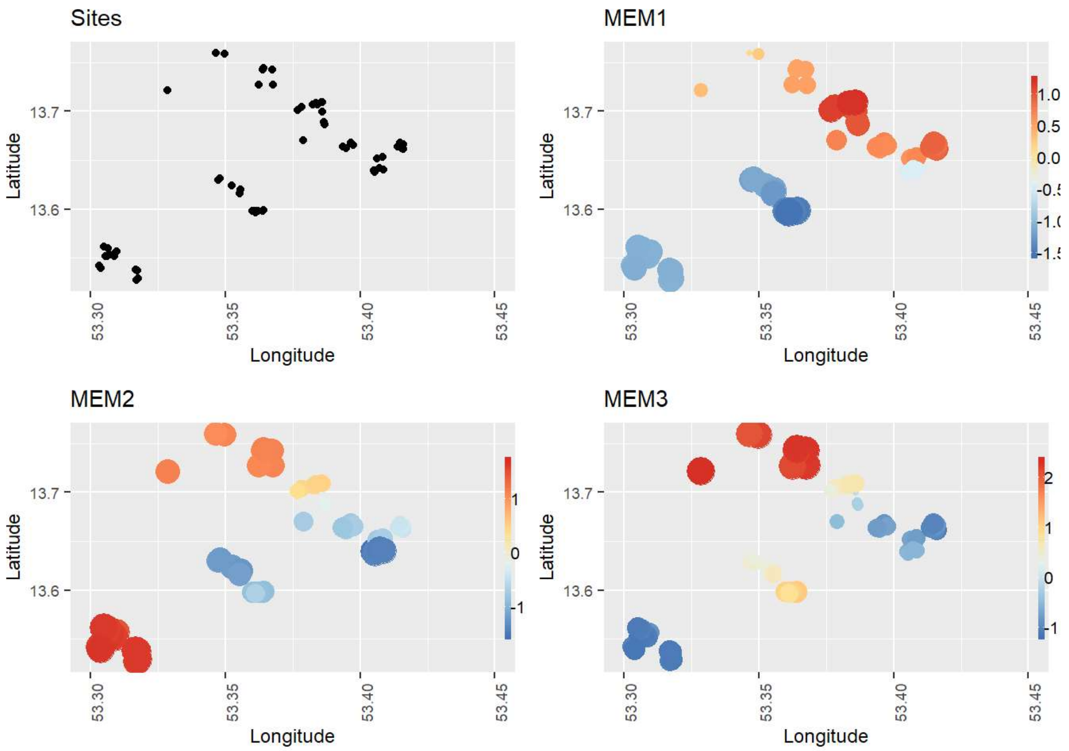


---

### 4. Regressing each SAC patterns (MEM vectors) using the taxa data

To explore which taxa reveal a strong association with each SAC pattern, we use a random forest algorithm. Since a relatively large instability is expected due to the excessive amount of predictors compared to the sample size, we use 10-fold subsampling to assess the relative importance of the predictors and model accuracy.

```
for(j in dbmem.selected){  #length(dbmem.selected)
    # selecting a MEM vector and combine it with the selected taxa data
    #j = dbmem.selected[1]
    df.2 = data.frame(dbmem.data0[,j],df_selected)
    colnames(df.2) = c(j, colnames(df_selected))

```

```
    # iteration group
    k_fold = 10
    set.seed(2)
    i_row = sample(c(1:k_fold), nrow(df.2), replace=T)
    R2.fitting    = numeric(0)
    R2.validation = numeric(0)
    vimp = matrix(NA, ncol = k_fold, nrow = length(df_selected))
    rownames(vimp) = colnames(df_selected)

    # parallelization
    parallelStartSocket(3)

    # RF modeling
    for(i_group in c(1:k_fold)){
        regr.task = makeRegrTask(data = df.2, target = j)
        regr.learner.cforest = makeLearner(cl="regr.cforest", predict.type
= "response", ntree = 1000)
        model.cforest = train(regr.learner.cforest,   regr.task, subset =
which(i_row!=i_group))
        fit.cforest   = predict(model.cforest, task = regr.task, subset =
which(i_row!=i_group))
        pred.cforest  = predict(model.cforest, task = regr.task, subset =
which(i_row==i_group))
        R2.fitting    = append(R2.fitting,    performance(fit.cforest,
measures = list(rsq)))
        R2.validation = append(R2.validation, performance(pred.cforest,
measures = list(rsq)))
        vimp[,i_group] = generateFilterValuesData(regr.task, method =
"cforest.importance")$data$cforest.importance
    }
    parallelStop()
    # calculating the mean and standard deviation for each taxon
    vimp_summary = data.frame(id = colnames(df_selected), ave =
apply(vimp,1,mean), stdev = apply(vimp,1,sd))
    vimp_summary = vimp_summary[order(vimp_summary$ave, decreasing = T),]

    # storing the R2 scores in the form of ggplot
    g = ggplot(data.frame(eval=append(rep("Fitted",
k_fold),rep("Validation", k_fold)),
                          score=append(R2.fitting, R2.validation)),
aes(x=eval, y=score)) +
        geom_boxplot() +
        labs(title="R-squared", x="", y="Score") +
        theme(text = element_text(size=9))
    eval(parse(text=(paste("g_eval_",j, "= g", sep=""))))

    # storing variable importance measure in the form of ggplot
    g = ggplot(vimp_summary[1:10, ], aes(x=reorder(id, -ave), y=ave)) +
        geom_bar(stat="identity") +
        geom_errorbar(aes(ymin=ave-stdev, ymax=ave+stdev), width=.2,
position=position_dodge(.9)) +
        labs(title="Relative variable importance", x="Taxa",
y="Importance") +
        theme(text = element_text(size=9), axis.text.x =
element_text(angle=90, hjust=1))

    eval(parse(text=(paste("g_vimp_",j, "= g", sep=""))))

}
## Starting parallelization in mode=socket with cpus=3.
## Stopped parallelization. All cleaned up.
```

```
## Starting parallelization in mode=socket with cpus=3.
## Stopped parallelization. All cleaned up.
## Starting parallelization in mode=socket with cpus=3.
## Stopped parallelization. All cleaned up.
print(g_1 / g_2 / g_3 | g_eval_MEM1 / g_eval_MEM2/ g_eval_MEM3 |
g_vimp_MEM1 / g_vimp_MEM2 / g_vimp_MEM3)
```

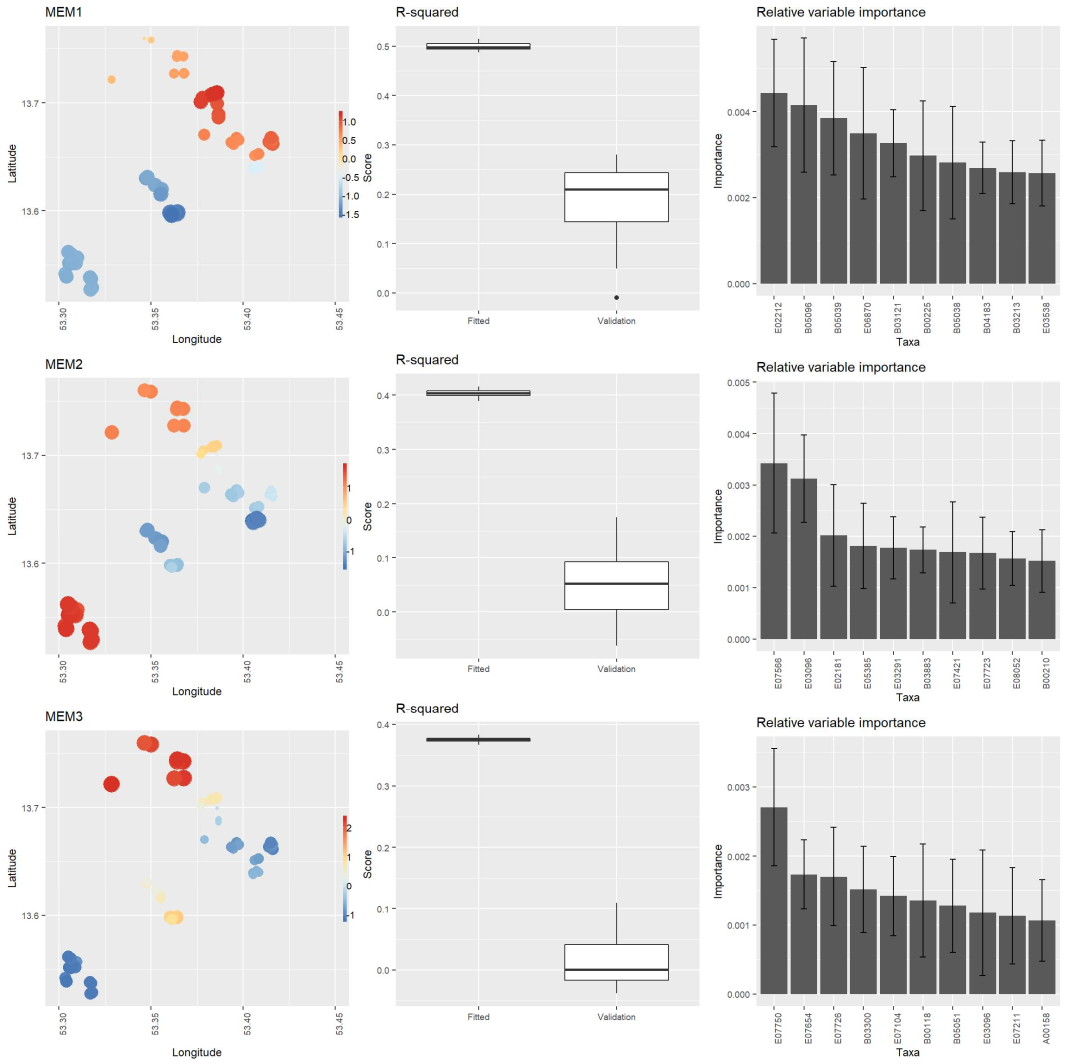


### Interpretation

None of the analyzed predictors revealed a strong match to any SAC patterns. Supporting this, cross Validation showed a strong decrease of R2 to nearly zero. Probably, no taxa have a strong spatial pattern in this region.
