## Supplementary material for "From microbes to mammals: pond biodiversity homogenization across different land-use types in an agricultural landscape": Table S1 sampling summary

| | Coordinates | | Water Collection | | | | | Sediment collection | | | | | In-Situ parameters | | | | | Water chemistry | | | | | Crops | | |
|---|---|---|---|---|---|---|---|---|---|---|---|---|---|---|---|---|---|---|---|---|---|---|---|---|---|
| #KH | Lat | Long | Dec '16 | Mar '17 | May '17 | Jun '17 | Oct '17 | Dec '16 | Mar '17 | May '17 | Jun '17 | Oct '17 | Dec '16 | Mar '17 | May '17 | Jun '17 | Oct '17 | Dec '16 | Mar '17 | May '17 | Jun '17 | Oct '17 | 2016 | 2017 | Landuse |
| 28 | 53.317754 | 13.528872 | dry | + | + | + | NA | + | - | - | - | NA | - | + | + | + | NA | NA | + | - | + | NA | Rye | Wheat | Arable |
| 40 | 53.317315 | 13.537215 | dry | + | + | dry | NA | + | - | - | - | NA | - | + | + | - | NA | NA | - | + | + | NA | Triticale | Corn | Arable |
| 133 | 53.367837 | 13.727254 | dry | dry | dry | dry | NA | - | + | - | - | NA | - | - | - | - | NA | NA | - | - | - | NA | NA | NA | Arable |
| 149 | 53.349858 | 13.758482 | dry | + | + | + | NA | - | + | - | - | NA | - | + | + | + | NA | NA | + | + | + | NA | Rape | Wheat | Arable |
| 183 | 53.367470 | 13.742717 | dry | dry | dry | dry | NA | x | + | - | - | NA | - | - | - | - | NA | NA | - | - | - | NA | NA | NA | Arable |
| 187 | 53.363995 | 13.742511 | dry | + | + | + | NA | + | + | - | - | NA | - | + | + | + | NA | NA | - | + | + | NA | Corn | Wheat | Arable |
| 190 | 53.362444 | 13.727117 | dry | + | + | dry | NA | + | + | - | - | NA | - | + | + | - | NA | NA | + | + | - | NA | Wheat | Barley | Arable |
| 235 | 53.327811 | 13.721543 | dry | dry | dry | dry | NA | - | - | - | - | NA | - | - | - | - | NA | NA | - | - | - | NA | NA | Rape | Arable |
| 236 | 53.328479 | 13.721488 | dry | dry | dry | dry | NA | - | + | - | - | NA | - | - | - | - | NA | NA | - | - | - | NA | NA | Rape | Arable |
| 258 | 53.382468 | 13.707032 | + | + | + | + | + | + | + | - | - | NA | - | + | + | + | + | NA | + | + | + | + | Barley | Rape | Arable |
| 259 | 53.384092 | 13.706979 | + | + | + | + | + | + | + | - | - | NA | - | + | + | + | + | NA | - | + | + | + | Barley | Rape | Arable |
| 265 | 53.378195 | 13.704492 | + | + | + | + | + | + | + | - | - | NA | - | + | + | + | + | NA | + | + | + | + | Wheat | Wheat | Arable |
| 269 | 53.376962 | 13.701001 | dry | + | + | + | + | + | + | - | - | NA | - | + | + | + | + | NA | - | + | + | + | Wheat | Corn | Arable |
| 275 | 53.385975 | 13.709487 | + | + | + | + | + | + | + | - | - | NA | - | + | + | + | + | NA | - | + | + | + | Rye | Barley | Arable |
| 287 | 53.385843 | 13.699214 | dry | + | + | + | + | + | + | - | - | NA | - | + | + | + | + | NA | + | + | + | + | Barley | Rape | Arable |
| 311 | 53.386548 | 13.689226 | dry | dry | + | dry | NA | - | + | - | - | NA | - | - | + | - | NA | NA | - | + | - | NA | Wheat | Rape | Arable |
| 312 | 53.386734 | 13.686318 | + | + | + | + | + | + | + | - | - | NA | - | + | + | + | + | NA | - | + | + | + | Wheat | Rape | Arable |
| 319 | 53.378829 | 13.670430 | dry | dry | + | dry | NA | x | - | - | + | NA | - | - | + | - | NA | NA | - | + | - | NA | NA | Rape | Arable |
| 606 | 53.347328 | 13.629944 | dry | + | + | + | NA | + | + | - | - | NA | - | + | + | + | NA | NA | - | + | + | NA | Corn | Corn | Arable |
| 607 | 53.347977 | 13.631487 | dry | + | + | dry | NA | + | + | - | - | NA | - | + | + | - | NA | NA | - | + | + | NA | Rape | Wheat | Arable |
| 608 | 53.348694 | 13.632055 | dry | dry | dry | + | NA | - | - | - | - | NA | - | - | - | + | NA | NA | - | - | - | NA | Rape | Wheat | Arable |
| 805 | 53.394665 | 13.662017 | dry | + | + | + | NA | + | + | - | - | NA | - | + | + | + | NA | NA | - | + | + | NA | Rape | Wheat | Arable |
| 807 | 53.397377 | 13.665798 | + | + | + | + | + | + | + | - | - | NA | - | + | + | + | + | NA | + | + | + | + | Rape | Wheat | Arable |
| 808 | 53.396518 | 13.667568 | dry | dry | dry | dry | NA | - | + | - | - | NA | - | - | - | - | NA | NA | - | - | - | NA | Rape | Wheat | Arable |
| 892 | 53.406355 | 13.651345 | + | + | + | + | + | + | + | - | - | NA | - | + | + | + | + | NA | + | - | + | + | Rape | Wheat | Arable |
| 893 | 53.408342 | 13.652892 | dry | + | + | + | NA | + | + | - | - | NA | - | + | + | + | NA | NA | + | - | + | NA | Rape | Wheat | Arable |
| 907 | 53.405524 | 13.638137 | + | + | + | + | + | + | + | - | - | NA | - | + | + | + | + | NA | - | + | + | + | Grassland | Grassland | Grassland |
| 908 | 53.405256 | 13.639553 | dry | dry | dry | dry | NA | x | + | - | - | NA | - | - | - | - | NA | NA | - | - | - | NA | Grassland | Grassland | Grassland |
| 910 | 53.407186 | 13.641936 | dry | + | + | dry | NA | + | + | - | + | NA | - | + | + | - | NA | NA | - | + | - | NA | Grassland | Grassland | Grassland |
| 911 | 53.408547 | 13.640171 | + | + | + | + | NA | + | + | - | - | NA | - | + | + | + | NA | NA | + | + | + | NA | Grassland | Grassland | Grassland |
| 939 | 53.415295 | 13.662426 | dry | + | + | + | NA | - | + | - | - | NA | - | + | + | + | NA | NA | - | + | + | NA | Grassland | Grassland | Grassland |
| 940 | 53.415256 | 13.664351 | dry | + | + | + | + | + | + | - | - | NA | - | + | + | + | + | NA | + | + | + | + | Grassland | Grassland | Grassland |
| 942 | 53.416030 | 13.665971 | dry | + | + | + | + | + | + | - | - | NA | - | + | + | + | + | NA | + | + | + | + | Grassland | Grassland | Grassland |
| 1189 | 53.353518 | 13.618345 | dry | + | + | + | NA | + | + | - | - | NA | - | + | + | + | NA | NA | + | + | + | NA | Barley | Corn | Arable |
| 1228 | 53.364171 | 13.598962 | dry | + | + | dry | NA | - | + | - | + | NA | - | + | + | - | NA | NA | + | + | - | NA | Forest | Corn | Arable |
| 1229 | 53.363931 | 13.597662 | dry | + | + | dry | NA | - | - | - | - | NA | - | + | + | - | NA | NA | + | + | - | NA | Forest | Corn | Arable |
| 1338 | 53.309612 | 13.556842 | dry | + | + | dry | NA | - | + | - | + | NA | - | + | + | - | NA | NA | + | + | - | NA | Wheat | Barley | Arable |
| 1510 | 53.395027 | 13.685878 | dry | dry | + | dry | NA | + | + | - | + | NA | - | - | + | - | NA | NA | - | + | - | NA | Wheat | Barley | Arable |
| 1590 | 53.303302 | 13.541943 | + | + | + | + | NA | + | + | - | - | NA | - | + | + | + | NA | NA | + | + | + | NA | Barley | Rape | Forest |
| 1598 | 53.308537 | 13.553018 | dry | + | + | + | NA | x | + | - | - | NA | - | + | + | + | NA | NA | + | + | + | NA | Wheat | Wheat | Arable |
| 1599 | 53.308914 | 13.551857 | + | + | + | dry | NA | x | - | - | + | NA | - | + | + | - | NA | NA | + | + | - | NA | Wheat | Wheat | Arable |
| 1604 | 53.306172 | 13.551442 | dry | + | + | + | NA | + | - | - | - | NA | - | + | + | + | NA | NA | + | + | + | NA | Wheat | Wheat | Arable |
| 2484 | 53.352275 | 13.623681 | dry | + | + | + | NA | + | + | - | - | NA | - | + | + | + | NA | NA | + | + | - | NA | Barley | Corn | Arable |
| 2489 | 53.355749 | 13.620038 | dry | dry | + | + | NA | - | + | - | - | NA | - | - | + | + | NA | NA | - | + | + | NA | Barley | Corn | Arable |
| 2565 | 53.306179 | 13.558765 | dry | + | + | + | NA | x | + | - | - | NA | - | + | + | + | NA | NA | - | + | + | NA | Rape | Wheat | Forest |
| A | 53.413945 | 13.663804 | NA | + | + | + | NA | NA | + | - | - | NA | NA | + | + | + | NA | NA | + | + | + | NA | Forest | Forest | Forest |
| B | 53.414943 | 13.667696 | NA | + | + | + | NA | NA | + | - | - | NA | NA | + | + | + | NA | NA | - | + | + | NA | Grassland | Grassland | Grassland |
| C | 53.415976 | 13.661671 | NA | dry | + | dry | NA | NA | + | - | - | NA | NA | - | + | - | NA | NA | - | - | - | NA | Grassland | Grassland | Grassland |
| D | 53.385634 | 13.709338 | NA | + | + | + | dry | NA | - | - | - | NA | NA | + | + | + | - | NA | + | + | + | - | Barley | Rape | Arable |
| E | 53.383527 | 13.708374 | NA | + | + | + | + | NA | - | - | - | NA | NA | + | + | + | + | NA | - | + | + | - | Barley | Rape | Arable |
| F | 53.393498 | 13.663738 | NA | + | + | + | NA | NA | - | - | - | NA | NA | + | + | + | NA | NA | + | + | + | NA | Rape | Wheat | Arable |
| G | 53.355291 | 13.615695 | NA | + | + | + | NA | NA | - | - | - | NA | NA | + | + | + | NA | NA | - | + | + | NA | Corn | Wheat | Arable |
| H | 53.346533 | 13.759664 | NA | + | + | dry | NA | NA | + | - | + | NA | NA | + | + | - | NA | NA | + | + | - | NA | Rape | Wheat | Arable |
| I | 53.364154 | 13.744220 | NA | + | + | dry | NA | NA | - | - | - | NA | NA | + | + | - | NA | NA | + | + | - | NA | Corn | Wheat | Arable |
| J | 53.306989 | 13.553221 | NA | + | + | + | NA | NA | + | - | - | NA | NA | + | + | + | NA | NA | + | - | + | NA | Wheat | Wheat | Arable |

| #KH | Coordinates | | Water Collection | | | | | Sediment collection | | | | | In-Situ parameters | | | | | Water chemistry | | | | | Crops | | Landuse |
|---|---|---|---|---|---|---|---|---|---|---|---|---|---|---|---|---|---|---|---|---|---|---|---|---|---|
| | Lat | Long | Dec '16 | Mar '17 | May '17 | Jun '17 | Oct '17 | Dec '16 | Mar '17 | May '17 | Jun '17 | Oct '17 | Dec '16 | Mar '17 | May '17 | Jun '17 | Oct '17 | Dec '16 | Mar '17 | May '17 | Jun '17 | Oct '17 | 2016 | 2017 | |
| K | 53.306335 | 13.559830 | NA | + | + | + | NA | NA | + | - | - | NA | NA | + | + | + | NA | NA | + | + | + | NA | Rape | Wheat | Forest |
| L | 53.305043 | 13.561817 | NA | + | + | + | NA | NA | - | - | - | NA | NA | + | + | + | NA | NA | + | + | + | NA | Rape | Wheat | Arable |
| M | 53.303807 | 13.539182 | NA | + | + | dry | NA | NA | - | - | + | NA | NA | + | + | - | NA | NA | + | + | - | NA | Barley | Triticale | Arable |
| N | 53.316667 | 13.538287 | NA | + | + | + | NA | NA | - | - | - | NA | NA | + | + | + | NA | NA | + | + | + | NA | Triticale | Corn | Arable |
| O | 53.360647 | 13.598382 | NA | + | + | + | NA | NA | + | - | - | NA | NA | + | + | + | NA | NA | - | + | + | NA | Forest | Forest | Forest |
| P | 53.360999 | 13.598467 | NA | + | + | + | NA | NA | + | - | - | NA | NA | + | + | + | NA | NA | + | + | + | NA | Forest | Forest | Forest |
| Q | 53.360195 | 13.598307 | NA | + | + | + | NA | NA | + | - | - | NA | NA | + | + | + | NA | NA | - | + | + | NA | Forest | Forest | Forest |
| R | 53.360190 | 13.598448 | NA | + | + | + | NA | NA | + | - | - | NA | NA | + | + | + | NA | NA | - | + | + | NA | Forest | Forest | Forest |
| S | 53.361987 | 13.597970 | NA | + | + | + | NA | NA | + | - | - | NA | NA | + | + | + | NA | NA | - | + | + | NA | Forest | Forest | Forest |
| T | 53.361270 | 13.596169 | NA | + | + | dry | NA | NA | + | - | + | NA | NA | + | + | - | NA | NA | + | - | - | NA | Forest | Forest | Forest |
