## Supplementary material for "From microbes to mammals: pond biodiversity homogenization across different land-use types in an agricultural landscape": Table S2 Physico-Chemical paramteres

| | | | General descriptors | | | | | | Physical and Chemical paramters | | | | | | | | | | | | | | | | | | | | | | | | | | | | |
|---|---|---|---|---|---|---|---|---|---|---|---|---|---|---|---|---|---|---|---|---|---|---|---|---|---|---|---|---|---|---|---|---|---|---|---|---|---|
| Sample Name | KH# | Month | Landuse | Reed | Canopy | Size | Depth | Perm. | pH | Cond. µS | Temp | Tiefe | O2 Sat | O2 mg/L | Cl | SO4 | TP | DOC | NO3-N | NH4-N | NO2 | TN | o-PO4-P | o-PO4 | TP | Na | K | Mg | Ca | Br | TFe | TOC | SAK | NO2-N | E0 | Eh | An-N |
| 1189W2 | 1189 | Mar | Arable | Y | N | M | S | 3 | 7.87 | 571 | 7.98 | 35.00 | 105.6 | 12.15 | 20.98 | 117.83 | 0.06 | 15.99 | 4.77 | 0.01 | 0.01 | 7.52 | 0.02 | 0.05 | 0.04 | 6.25 | 15.77 | 4.81 | 95.14 | 0.00 | 0.10 | 18.54 | 44.02 | 0.00 | NA | NA | 4.79 |
| 1189W3 | 1189 | May | Arable | Y | N | M | S | 3 | 7.19 | 459 | 17.92 | 35.00 | 98 | 8.92 | 12.47 | 60.20 | 0.16 | 21.31 | 0.00 | 0.00 | NA | 1.64 | 0.10 | 0.29 | 0.06 | 8.13 | 12.58 | 7.32 | 65.72 | 0.00 | 0.27 | 21.47 | NA | NA | NA | NA | NA |
| 1189W4 | 1189 | Jun | Arable | Y | N | M | S | 3 | 6.90 | 468 | 19.9 | 35.00 | 0.4 | 0.04 | 19.08 | 43.80 | 0.79 | 25.33 | 0.04 | 0.02 | NA | 2.32 | 0.55 | 1.70 | 0.23 | 7.42 | 17.35 | 6.43 | 62.87 | 0.00 | 0.12 | NA | 3.48 | NA | -22.00 | 188.00 | NA |
| 1228W2 | 1228 | Mar | Arable | N | Y | S | S | 2 | 6.63 | 580 | 9.36 | 30.00 | 86 | 9.57 | 19.92 | 79.15 | 0.21 | 21.23 | 11.15 | NA | 0.00 | 14.02 | 0.20 | 0.61 | 0.01 | 7.43 | 21.44 | 7.80 | 69.86 | 0.00 | NA | 22.14 | NA | 0.00 | NA | NA | NA |
| 1228W3 | 1228 | May | Arable | N | Y | S | S | 2 | 6.06 | 333 | 13.06 | 20.00 | 17.9 | 1.84 | 15.74 | 90.41 | 1.14 | 42.45 | 0.01 | 0.14 | NA | 2.85 | 0.81 | NA | NA | 8.84 | 8.11 | 7.09 | 44.96 | NA | 0.41 | 44.82 | 156.06 | NA | NA | NA | NA |
| 1229W2 | 1229 | Mar | Arable | N | Y | S | S | 2 | 7.65 | 270 | 8.74 | 30.00 | 34.8 | 3.9 | 11.33 | 62.92 | 0.55 | 25.34 | 0.75 | NA | 0.00 | 2.17 | 0.52 | 1.60 | 0.03 | 2.50 | 22.62 | 4.96 | 40.17 | 0.00 | NA | 26.94 | NA | 0.00 | NA | NA | NA |
| 1229W3 | 1229 | May | Arable | N | Y | S | S | 2 | 6.80 | 251 | 12.25 | 20.00 | 4.97 | 47.4 | 13.58 | 29.87 | 1.86 | 57.85 | 0.02 | 0.39 | NA | 3.29 | 1.20 | NA | NA | 4.96 | 22.57 | 5.17 | 22.94 | 0.05 | 0.34 | 58.56 | 224.04 | NA | NA | NA | NA |
| 1338W2 | 1338 | Mar | Arable | N | N | M | S | 2 | 7.80 | 207 | 10.92 | 20.00 | 98 | 10.5 | 11.07 | 32.32 | 0.21 | 12.68 | 2.79 | 1.21 | 0.12 | 6.37 | 0.02 | 0.06 | 0.19 | 3.72 | 10.95 | 3.04 | 40.76 | 0.00 | 0.81 | 15.74 | 49.75 | 0.04 | NA | NA | 4.03 |
| 1338W3 | 1338 | May | Arable | N | N | M | S | 2 | 7.99 | 195 | 21.09 | 15.00 | 70.3 | 6.14 | 13.16 | 18.44 | 0.90 | 32.48 | 0.05 | 0.17 | NA | 2.98 | 0.30 | NA | NA | 2.97 | 21.08 | 6.10 | 14.97 | 0.04 | 0.26 | 36.51 | 87.78 | NA | NA | NA | NA |
| 149W2 | 149 | Mar | Arable | N | Y | L | S | 3 | 7.60 | 1102 | 9.46 | 70.00 | 72.3 | 7.91 | 38.47 | 136.42 | 0.07 | 16.83 | 5.58 | NA | 0.00 | 7.31 | 0.08 | 0.24 | -0.01 | 17.16 | 9.76 | 16.50 | 130.26 | 0.00 | NA | 17.61 | NA | 0.00 | NA | NA | NA |
| 149W3 | 149 | May | Arable | N | Y | L | S | 3 | 7.74 | 1162 | 8.35 | 40.00 | 51.3 | 5.77 | 49.40 | 177.99 | 0.07 | 19.54 | 0.09 | 0.07 | NA | 1.88 | 0.07 | 0.21 | 0.00 | 24.26 | 8.35 | 22.73 | 143.97 | 0.00 | 0.09 | 23.55 | NA | NA | NA | NA | NA |
| 149W4 | 149 | Jun | Arable | N | Y | L | S | 3 | 8.26 | 1100 | 18.3 | 25.00 | 77.7 | 7.51 | 50.78 | 98.60 | 0.61 | 30.30 | 0.06 | 0.04 | NA | 1.96 | 0.55 | 1.70 | 0.05 | 20.57 | 9.58 | 16.70 | 151.08 | 0.40 | 0.05 | NA | 3.44 | NA | 115.90 | 325.90 | NA |
| 1510W3 | 1510 | May | Arable | N | N | S | S | 1 | 10.82 | 520 | 17.38 | 15.00 | 278.9 | 25.9 | 58.97 | 25.69 | 0.97 | 40.66 | 0.66 | 0.22 | 0.41 | 9.58 | 0.29 | 0.89 | 0.68 | 8.75 | 33.73 | 28.92 | 48.23 | 0.20 | 0.34 | 49.01 | 69.65 | 0.12 | NA | NA | 1.01 |
| 1590W2 | 1590 | Mar | Forest | N | Y | L | S | 3 | 7.90 | 588 | 9.83 | 50.00 | 52 | 5.7 | 26.20 | 84.87 | 0.05 | 26.65 | 4.34 | 0.09 | 0.25 | 7.19 | 0.02 | 0.05 | 0.03 | 8.29 | 8.10 | 7.52 | 86.29 | 0.00 | 0.09 | 27.88 | 81.22 | 0.08 | NA | NA | 4.50 |
| 1590W3 | 1590 | May | Forest | N | Y | L | S | 3 | 8.12 | 6.56 | 14.92 | 40.00 | 70.4 | 6.94 | NA | NA | NA | NA | NA | NA | NA | NA | NA | NA | NA | NA | NA | NA | NA | NA | NA | NA | NA | NA | NA | NA | NA |
| 1590W4 | 1590 | Jun | Forest | N | Y | L | S | 3 | 7.29 | 612 | 15.1 | 30.00 | 23 | 2.28 | 24.94 | 38.83 | 0.37 | 34.04 | 0.06 | 0.11 | NA | 2.29 | 0.17 | 0.52 | 0.20 | 5.60 | 5.01 | 3.69 | 30.57 | 0.10 | 0.04 | NA | 3.64 | NA | 117.20 | 327.20 | NA |
| 1598W2 | 1598 | Mar | Arable | Y | P | L | D | 3 | 8.21 | 208 | 6.98 | 150.00 | 57.7 | 6.92 | 17.06 | 33.82 | 0.52 | 20.78 | 0.02 | 0.08 | 0.00 | 2.03 | 0.39 | 1.19 | 0.13 | 4.47 | 10.72 | 2.87 | 41.30 | 0.00 | 0.06 | 21.57 | 71.70 | 0.00 | NA | NA | 0.10 |
| 1598W3 | 1598 | May | Arable | Y | P | L | D | 3 | 7.26 | 303 | 16.07 | 100.00 | 40.9 | 3.96 | 19.88 | 20.87 | 0.67 | 22.74 | 0.03 | 0.13 | NA | 1.51 | 0.53 | NA | NA | 7.63 | 15.54 | 6.39 | 33.48 | 0.05 | 0.19 | 24.77 | 67.80 | NA | NA | NA | NA |
| 1598W4 | 1598 | Jun | Arable | Y | P | L | D | 3 | 6.49 | 316 | 15.9 | 30.00 | 37.2 | 3.5 | 20.73 | 5.66 | 1.19 | 24.15 | 0.03 | 0.03 | NA | 1.89 | 1.20 | 3.68 | -0.01 | 8.00 | 14.18 | 5.95 | 34.48 | 0.01 | 0.06 | NA | 3.09 | NA | 172.90 | 382.90 | NA |
| 1599W2 | 1599 | Mar | Arable | Y | N | M | S | 2 | 7.70 | 340 | 6.98 | 30.00 | 75.1 | 8.94 | 15.70 | 61.83 | 0.08 | 20.73 | 0.21 | 0.06 | 0.00 | 1.90 | 0.04 | 0.13 | 0.04 | 3.09 | 6.51 | 4.40 | 40.92 | 0.00 | 0.05 | 22.75 | 72.00 | 0.00 | NA | NA | 0.27 |
| 1599W3 | 1599 | May | Arable | Y | N | M | S | 2 | 7.97 | 550 | 15.16 | 15.00 | 18.5 | 1.82 | NA | NA | NA | NA | NA | NA | NA | NA | NA | NA | NA | NA | NA | NA | NA | NA | NA | NA | NA | NA | NA | NA | NA |
| 1604W2 | 1604 | Mar | Arable | Y | P | L | S | 3 | 8.10 | 638 | 8.99 | 30.00 | 125 | 14 | 61.19 | 78.70 | 0.05 | 13.21 | 1.20 | 0.00 | 0.00 | 2.66 | 0.04 | 0.12 | 0.01 | 18.57 | 5.57 | 6.83 | 93.08 | 0.06 | 0.17 | 14.70 | 40.77 | 0.00 | NA | NA | 1.21 |
| 1604W3 | 1604 | May | Arable | Y | P | L | S | 3 | 8.19 | 775 | 14.99 | 50.00 | 18 | 1.78 | 45.71 | 55.07 | 0.34 | 15.28 | 0.03 | 0.14 | NA | 1.38 | 0.07 | NA | NA | 24.91 | 12.18 | 11.71 | 62.00 | 0.03 | 0.02 | 15.31 | 42.00 | NA | NA | NA | NA |
| 1604W4 | 1604 | Jun | Arable | Y | P | L | S | 3 | 7.37 | 795 | 18.2 | 50.00 | 33.7 | 3.13 | 38.53 | 15.67 | 0.60 | 16.60 | 0.05 | 0.01 | NA | 1.33 | 0.21 | 0.66 | 0.39 | 7.82 | 5.04 | 3.81 | 38.25 | 0.75 | 0.07 | NA | 3.13 | NA | 166.70 | 376.70 | NA |
| 187W3 | 187 | May | Arable | Y | N | M | S | 3 | 7.49 | 1084 | 11.06 | 60.00 | 33.8 | 3.57 | 33.86 | 213.88 | 0.16 | 18.49 | 0.00 | 0.01 | NA | 1.98 | 0.13 | 0.39 | 0.03 | 15.28 | 13.73 | 15.98 | 146.03 | 0.00 | 0.08 | 21.81 | NA | NA | NA | NA | NA |
| 187W4 | 187 | Jun | Arable | Y | N | M | S | 3 | 7.72 | 1133 | 19.6 | 50.00 | 31 | 2.84 | 32.53 | 147.84 | 0.96 | 22.82 | 0.05 | 0.03 | NA | 2.37 | 0.89 | 2.72 | 0.07 | 16.97 | 9.24 | 16.75 | 190.45 | 0.63 | 0.02 | NA | 2.83 | NA | 43.30 | 253.30 | NA |
| 190W2 | 190 | Mar | Arable | Y | N | M | S | 2 | 6.10 | 1604 | 14.23 | 30.00 | 67 | 6.6 | 40.91 | 609.05 | 0.11 | 15.48 | 11.49 | NA | 0.00 | 15.60 | 0.05 | 0.14 | 0.06 | 30.80 | 25.45 | 14.62 | 237.90 | 0.00 | NA | 16.78 | NA | 0.00 | NA | NA | NA |
| 190W3 | 190 | May | Arable | Y | N | M | S | 2 | 7.10 | 779 | 13.53 | 50.00 | 113.1 | 11.3 | 10.63 | 268.92 | 0.24 | 30.80 | 0.00 | 0.09 | NA | 3.53 | 0.17 | 0.52 | 0.07 | 6.13 | 16.09 | 10.14 | 141.14 | 0.00 | 0.80 | 35.68 | NA | NA | NA | NA | NA |
| 2484W2 | 2484 | Mar | Arable | Y | N | L | D | 3 | 8.97 | 449 | 7.43 | 170.00 | 135 | 9.37 | 19.52 | 33.88 | 0.08 | 16.62 | 3.87 | 0.04 | 0.30 | 6.20 | 0.02 | 0.06 | 0.06 | 7.10 | 4.17 | 6.05 | 93.46 | 0.00 | 0.19 | 16.76 | 29.11 | 0.09 | NA | NA | 4.00 |
| 2484W3 | 2484 | May | Arable | Y | N | L | D | 3 | 9.04 | 448 | 13.45 | 200.00 | 141 | 14.12 | 16.21 | 33.07 | 0.39 | 14.90 | 1.16 | 0.05 | NA | 3.22 | 0.11 | 0.33 | 0.29 | 10.18 | 9.67 | 8.46 | 47.82 | 0.00 | 0.54 | 18.98 | NA | NA | NA | NA | NA |
| 2489W3 | 2489 | May | Arable | Y | N | S | S | 2 | 8.16 | 574 | 17.19 | 40.00 | 241.4 | 22.27 | 12.87 | 15.28 | 0.97 | 23.54 | 0.01 | 0.04 | NA | 2.49 | 0.90 | 2.75 | 0.08 | 5.64 | 24.83 | 6.28 | 46.04 | 0.10 | 0.22 | 30.27 | NA | NA | NA | NA | NA |
| 2489W4 | 2489 | Jun | Arable | Y | N | S | S | 2 | 6.77 | 656 | 21.1 | 30.00 | 79.2 | 2.58 | 24.77 | 1.38 | 2.67 | 51.52 | 0.09 | 0.18 | NA | 5.11 | 0.24 | 0.74 | 2.43 | 4.75 | 22.89 | 5.36 | 41.87 | 0.17 | 10.15 | NA | 3.60 | NA | -1.00 | 209.00 | NA |
| 2565W3 | 2565 | May | Forest | N | Y | L | S | 3 | 7.78 | 275 | 15.75 | 90.00 | 31.9 | 3.11 | 20.28 | 22.74 | 0.65 | 41.19 | 0.09 | 0.55 | NA | 3.56 | 0.35 | NA | NA | 6.87 | 13.22 | 5.37 | 31.07 | 0.02 | 0.85 | 49.87 | 184.83 | 0.00 | NA | NA | NA |
| 2565W4 | 2565 | Jun | Forest | N | Y | L | S | 3 | 6.28 | 234 | 15.4 | 90.00 | 18.3 | 1.8 | 17.06 | 13.02 | 0.35 | 43.25 | 0.02 | 0.02 | NA | 3.23 | 0.26 | 0.80 | 0.08 | 5.49 | 8.25 | 4.33 | 25.23 | 0.10 | 0.17 | NA | 3.99 | NA | 162.30 | 372.30 | NA |
| 258W2 | 258 | Mar | Arable | N | N | M | D | 3 | 7.63 | 743 | 6.92 | 100.00 | 106.9 | 12.7 | 25.60 | 153.61 | 0.05 | 12.59 | 4.81 | 0.01 | 0.08 | 6.96 | 0.02 | 0.06 | 0.03 | 9.04 | 9.40 | 9.92 | 104.18 | 0.00 | 0.05 | 13.46 | 24.70 | 0.02 | NA | NA | 4.84 |
| 258W3 | 258 | May | Arable | N | N | M | D | 3 | 7.89 | 606 | 14.61 | 150.00 | 95.5 | 9.34 | 21.03 | 151.41 | 0.07 | 13.80 | 0.00 | 0.18 | NA | 1.59 | 0.04 | 0.12 | 0.03 | 10.61 | 11.84 | 13.41 | 83.49 | 0.00 | 0.14 | 14.91 | NA | NA | NA | NA | NA |
| 258W4 | 258 | Jun | Arable | N | N | M | D | 3 | 7.65 | 676 | 21.56 | 110.00 | 46.1 | 3.95 | 22.68 | 129.49 | 0.30 | 14.72 | 0.00 | 0.03 | NA | 1.77 | 0.22 | 0.69 | 0.07 | 12.86 | 11.73 | 14.20 | 94.05 | 0.00 | 0.06 | NA | 2.33 | NA | NA | NA | NA |
| 258W5 | 258 | Oct | Arable | N | N | M | D | 3 | 7.67 | 627 | 11.9 | NA | NA | NA | 29.03 | 90.59 | 0.43 | 17.40 | 0.04 | 1.58 | NA | 3.18 | 0.35 | 1.07 | 0.08 | 13.70 | 16.37 | 14.12 | 95.81 | 0.00 | 0.20 | 18.62 | 48.06 | NA | 172.40 | 382.40 | NA |
| 259W3 | 259 | May | Arable | Y | N | L | S | 3 | 6.99 | 646 | 11.48 | 80.00 | 125.2 | 1.11 | 26.99 | 149.12 | 0.08 | 22.36 | 0.12 | 0.08 | NA | 2.61 | 0.05 | 0.14 | 0.03 | 13.09 | 6.66 | 8.05 | 102.69 | 0.00 | 0.16 | 22.95 | NA | NA | NA | NA | NA |
| 259W4 | 259 | Jun | Arable | Y | N | L | S | 3 | 7.10 | 739 | 21.4 | NA | 79.5 | 7.15 | 37.75 | 99.28 | 1.64 | 37.51 | 0.04 | 0.05 | NA | 3.60 | 0.54 | 1.66 | 1.10 | 5.68 | 3.22 | 2.36 | 29.01 | 0.02 | 0.09 | NA | 2.94 | NA | 155.20 | 365.20 | NA |
| 259W5 | 259 | Oct | Arable | Y | N | L | S | 3 | 7.19 | 625 | 11 | NA | NA | NA | 40.35 | 25.11 | 1.36 | 36.17 | 0.04 | 0.02 | NA | 3.13 | 1.85 | 5.67 | 0.49 | 17.18 | 12.28 | 9.40 | 105.71 | 0.11 | 0.50 | 37.61 | 101.36 | NA | 141.20 | 351.20 | NA |
| 265W2 | 265 | Mar | Arable | Y | N | L | D | 3 | 7.21 | 1000 | 5.8 | 75.00 | 112 | 13.84 | 37.82 | 249.84 | 0.03 | 15.63 | 9.59 | 0.01 | 0.03 | 14.75 | 0.02 | 0.06 | 0.01 | 11.33 | 12.66 | 8.43 | 138.15 | 0.00 | 0.07 | 15.85 | 31.06 | 0.01 | NA | NA | 9.61 |
| 265W3 | 265 | May | Arable | Y | N | L | D | 3 | 8.25 | 890 | 11.95 | 120.00 | 119.8 | 12.4 | 37.21 | 257.66 | 0.03 | 18.86 | 1.63 | 0.00 | NA | 17.20 | 0.01 | 0.03 | 0.02 | 11.71 | 17.20 | 10.99 | 141.46 | 0.00 | 0.06 | 19.58 | NA | NA | NA | NA | NA |
| 265W4 | 265 | Jun | Arable | Y | N | L | D | 3 | 7.90 | 923 | 21.55 | 100.00 | 33.9 | 2.93 | 38.07 | 220.57 | 0.41 | 25.63 | 0.00 | 0.01 | NA | 2.01 | 0.35 | 1.06 | 0.06 | 13.06 | 17.46 | 11.66 | 110.97 | 0.00 | 0.08 | NA | 2.85 | NA | NA | NA | NA |
| 265W5 | 265 | Oct | Arable | Y | N | L | D | 3 | 7.22 | 737 | 12.6 | NA | 50.5 | 5.29 | 38.79 | 126.88 | 0.71 | 20.52 | 0.04 | 0.10 | NA | 1.96 | 0.63 | 1.92 | 0.08 | 13.11 | 22.80 | 10.95 | 125.64 | 0.00 | 0.15 | 21.67 | 53.31 | NA | 177.60 | 387.60 | NA |
| 269W3 | 269 | May | Arable | N | P | M | S | 3 | 7.73 | 1074 | 11.73 | 90.00 | 57.2 | 5.94 | 48.63 | 283.83 | 0.82 | 32.26 | 0.39 | 0.17 | NA | 4.05 | 0.75 | 2.31 | 0.06 | 29.40 | 22.26 | 16.21 | 158.98 | 0.00 | 0.10 | 33.26 | NA | NA | NA | NA | NA |
| 269W4 | 269 | Jun | Arable | N | P | M | S | 3 | 8.20 | 1235 | 22.6 | NA | 91 | 7.8 | 52.06 | 196.24 | 1.78 | 35.98 | 0.07 | 0.02 | NA | 3.46 | 0.73 | 2.23 | 1.06 | 8.01 | 21.03 | 6.34 | 57.86 | 0.43 | 0.04 | NA | 3.06 | NA | 166.40 | 376.40 | NA |
| 269W5 | 269 | Oct | Arable | N | P | M | S | 3 | 7.87 | 861 | 12.7 | NA | 29.2 | 288 | 30.74 | 61.94 | 1.75 | 29.92 | 0.04 | 0.00 | NA | 2.37 | 1.85 | 5.68 | -0.10 | 16.66 | 37.44 | 17.70 | 147.32 | 0.00 | 0.28 | 30.89 | 80.02 | NA | 165.10 | 375.10 | NA |
| 275W3 | 275 | May | Arable | Y | N | L | D | 3 | 8.10 | 438 | 13.35 | 50.00 | 124.8 | 12.53 | 15.84 | 105.42 | 0.16 | 20.01 | 0.00 | 0.02 | NA | 2.08 | 0.08 | 0.24 | 0.08 | 7.34 | 8.99 | 8.45 | 54.76 | NA | 0.10 | 20.59 | NA | NA | NA | NA | NA |
| 275W4 | 275 | Jun | Arable | Y | N | L | D | 3 | 7.91 | 447 | 21 | NA | 78.5 | 7 | 14.13 | 85.75 | 0.12 | 21.14 | 0.02 | 0.02 | NA | 2.13 | 0.06 | 0.18 | 0.06 | 7.54 | 6.99 | 7.91 | 53.92 | 0.10 | 0.04 | NA | 2.78 | NA | 149.60 | 359.60 | NA |
| 275W5 | 275 | Oct | Arable | Y | N | L | D | 3 | 7.88 | 418 | 11 | NA | NA | NA | 15.75 | 15.12 | 0.43 | 23.34 | 0.06 | 0.01 | NA | 1.79 | 0.40 | 1.23 | 0.03 | 8.82 | 17.64 | 9.68 | 60.46 | 0.00 | 0.21 | 24.11 | 63.92 | NA | 153.50 | 363.50 | NA |
| 287W2 | 287 | Mar | Arable | Y | P | M | S | 3 | 7.76 | 790 | 5.47 | 30.00 | 63.1 | 7.76 | 26.06 | 184.61 | 0.41 | 31.45 | 19.01 | 1.21 | 0.65 | 28.00 | 0.18 | 0.56 | 0.22 | 7.48 | 24.37 | 8.97 | 82.40 | 0.00 | 0.06 | 32.90 | 73.06 | 0.20 | NA | NA | 20.42 |
| 287W3 | 287 | May | Arable | Y | P | M | S | 3 | 6.92 | 434 | 16.42 | 60.00 | 73.21 | 6.89 | 31.14 | 103.76 | 0.13 | 41.72 | 0.00 | 0.28 | NA | 4.75 | 1.68 | 5.14 | -1.54 | 11.00 | 24.64 | 9.56 | 74.41 | 0.00 | 0.40 | 49.20 | NA | NA | NA | NA | NA |
| 287W4 | 287 | Jun | Arable | Y | P | M | S | 3 | 6.64 | 458 | 21.9 | NA | 51.7 | 4.46 | 23.58 | 31.27 | 2.13 | 51.23 | 0.05 | 0.02 | NA | 4.08 | 2.49 | 7.63 | -0.37 | 12.30 | 18.99 | 9.69 | 77.88 | 0.05 | 2.49 | NA | 3.39 | NA | 116.80 | 326.80 | NA |
| 287W5 | 287 | Oct | Arable | Y | P | M | S | 3 | 5.70 | 804 | 11.9 | 30.00 | 41.1 | 4.36 | 21.50 | 312.70 | 0.17 | 25.84 | 1.91 | 0.06 | NA | 4.22 | 0.07 | 0.21 | 0.10 | 12.16 | 16.93 | 12.90 | 135.17 | 0.00 | 0.55 | 28.06 | 107.15 | NA | 190.40 | 400.40 | NA |
| 28W2 | 28 | Mar | Arable | Y | P | M | S | 3 | 7.50 | 213 | 9.72 | 30.00 | 44 | 4.82 | 15.18 | 38.14 | 0.39 | 37.88 | 0.11 | 0.04 | 0.00 | 3.07 | 0.25 | 0.77 | 0.14 | 4.63 | 10.95 | 3.07 | 30.59 | 0.00 | 0.17 | 39.74 | 163.96 | 0.00 | NA | NA | 0.15 |
| 28W3 | 28 | Mar | Arable | Y | P | M | S | 3 | 7.84 | 305 | 14.35 | 15.00 | 13.2 | 1.32 | NA | NA | NA | NA | NA | NA | NA | NA | NA | NA | NA | NA | NA | NA | NA | NA | NA | NA | NA | NA | NA | NA | NA |
| 28W4 | 28 | Jun | Arable | Y | P | M | S | 3 | 6.19 | 234 | 17.3 | NA | 20.8 | 2 | 12.48 | 3.71 | 1.69 | 51.20 | 0.00 | 0.02 | NA | 3.26 | 0.73 | 2.24 | 0.96 | 3.12 | 8.00 | 1.63 | 9.29 | 0.00 | 0.70 | NA | 4.07 | NA | 97.20 | 307.20 | NA |

| | | | General descriptors | | | | | | Physical and Chemical paramters | | | | | | | | | | | | | | | | | | | | | | | | | | | | |
|---|---|---|---|---|---|---|---|---|---|---|---|---|---|---|---|---|---|---|---|---|---|---|---|---|---|---|---|---|---|---|---|---|---|---|---|---|---|
| Sample Name | KH# | Month | Landuse | Reed | Canopy | Size | Depth | Perm. | pH | Cond. µS | Temp | Tiefe | O2 Sat | O2 mg/L | Cl | SO4 | TP | DOC | NO3-N | NH4-N | NO2 | TN | o-PO4-P | o-PO4 | TP | Na | K | Mg | Ca | Br | TFe | TOC | SAK | NO2-N | E0 | Eh | An-N |
| 311W3 | 311 | May | Arable | N | P | L | S | 1 | 6.34 | 389 | 12.73 | 25.00 | 64.7 | 6.64 | 3.29 | 144.95 | 0.27 | 18.07 | 0.00 | 0.08 | NA | 1.87 | 0.22 | 0.66 | 0.06 | 1.97 | 13.97 | 6.80 | 40.44 | 0.00 | 0.23 | 19.44 | NA | NA | NA | NA | NA |
| 312W3 | 312 | May | Arable | Y | N | L | S | 3 | 8.64 | 647 | 14.83 | 25.00 | 84.4 | 8.27 | 24.41 | 141.38 | 0.06 | 23.06 | 1.25 | 0.10 | 0.07 | NA | 0.02 | 0.06 | 0.04 | 10.57 | 9.18 | 11.05 | 107.93 | 0.03 | 0.25 | 23.43 | 50.60 | 0.02 | NA | NA | 1.37 |
| 312W4 | 312 | Jun | Arable | Y | N | L | S | 3 | 7.17 | 611 | 19.9 | NA | 52 | 4.32 | 22.93 | 115.23 | 0.14 | 25.98 | 0.03 | 0.10 | NA | 2.16 | 0.04 | 0.11 | 0.11 | 10.84 | 5.26 | 10.15 | 106.48 | 0.09 | 0.90 | NA | 2.76 | NA | 161.90 | 371.90 | NA |
| 312W5 | 312 | Oct | Arable | Y | N | L | S | 3 | 9.08 | 252 | 13.2 | NA | NA | NA | 15.86 | 4.38 | 0.26 | 26.35 | 0.00 | 0.00 | NA | 2.18 | 0.09 | 0.28 | 0.17 | 10.35 | 4.90 | 4.04 | 39.05 | 0.00 | 3.34 | 29.29 | 68.16 | NA | 115.60 | 325.60 | NA |
| 319W3 | 319 | May | Arable | N | Y | S | S | 1 | 6.30 | 298 | 14.2 | 15.00 | 40.1 | 3.95 | NA | NA | NA | NA | NA | NA | NA | NA | NA | NA | NA | NA | NA | NA | NA | NA | NA | NA | NA | NA | NA | NA | NA |
| 40W3 | 40 | May | Arable | N | Y | M | S | 2 | 8.22 | 450 | 14.18 | 20.00 | 71 | 7.13 | 34.93 | 37.20 | 2.93 | 33.70 | 0.03 | 0.27 | NA | 2.74 | 1.51 | NA | NA | 10.13 | 29.40 | 8.90 | 51.60 | 0.02 | 0.30 | 35.61 | 109.74 | NA | NA | NA | NA |
| 40W4 | 40 | Jun | Arable | N | Y | M | S | 2 | 7.16 | 465 | 15 | NA | 8.9 | 0.9 | 31.96 | 26.58 | 1.40 | 33.43 | 0.02 | 1.70 | NA | 4.13 | 1.80 | 5.53 | -0.40 | 10.79 | 19.31 | 7.62 | 59.39 | 0.03 | 0.24 | NA | 3.86 | NA | 33.20 | 243.20 | NA |
| 606W3 | 606 | May | Arable | N | P | M | S | 3 | 9.50 | 540 | 19.99 | 50.00 | 133 | 11.59 | 17.21 | 66.20 | 0.67 | 24.91 | 0.00 | 0.15 | NA | 2.90 | 0.53 | 1.61 | 0.14 | 10.39 | 27.93 | 11.87 | 56.13 | 0.00 | 0.18 | 27.30 | NA | NA | NA | NA | NA |
| 606W4 | 606 | Jun | Arable | N | P | M | S | 3 | 7.43 | 674 | 21.3 | NA | 11.5 | 1.01 | 23.75 | 9.97 | 1.47 | 73.14 | 0.03 | 3.94 | NA | 8.97 | 2.68 | 8.21 | -1.21 | 10.04 | 33.34 | 10.69 | 71.65 | 0.48 | 0.84 | NA | 1.57 | NA | 106.90 | 316.90 | NA |
| 607W3 | 607 | May | Arable | Y | N | M | S | 3 | 7.79 | 688 | 15.09 | 30.00 | 68.6 | 6.61 | 20.16 | 83.41 | 0.09 | 37.55 | 0.00 | 0.10 | NA | 1.81 | 0.06 | 0.18 | 0.03 | 11.84 | 6.67 | 9.06 | 86.80 | 0.00 | 0.28 | 39.50 | NA | NA | NA | NA | NA |
| 607W4 | 607 | Jun | Arable | Y | N | M | S | 3 | 7.22 | 623 | 22.3 | NA | 35.6 | 3.05 | 20.37 | 15.82 | 0.66 | 55.53 | 0.03 | 0.07 | NA | 2.35 | 0.57 | 1.73 | 0.09 | 12.06 | 6.80 | 8.27 | 99.85 | 0.58 | 0.10 | NA | 1.77 | NA | 171.50 | 381.50 | NA |
| 805W3 | 805 | May | Arable | Y | Y | L | D | 3 | 7.34 | 572 | 9.45 | 50.00 | 61.2 | 6.77 | 33.58 | 184.38 | 0.09 | 21.39 | 0.06 | 0.03 | NA | 1.80 | 0.03 | 0.10 | 0.06 | 15.12 | 7.17 | 8.67 | 84.56 | 0.00 | 0.73 | 22.59 | NA | NA | NA | NA | NA |
| 805W4 | 805 | Jun | Arable | Y | Y | L | D | 3 | 6.60 | 540 | 18.4 | NA | 29.8 | 2.79 | 30.73 | 109.12 | 0.21 | 27.55 | 0.04 | 0.01 | NA | 1.73 | 0.16 | 0.48 | 0.06 | 16.29 | 4.03 | 9.79 | 87.90 | 0.03 | 0.12 | NA | 2.93 | NA | -38.70 | 171.30 | NA |
| 807W2 | 807 | Mar | Arable | Y | N | L | D | 3 | 7.88 | 301 | 7.19 | 100.00 | 102 | 11.99 | 15.94 | 31.54 | 0.05 | 13.71 | 0.18 | 0.00 | 0.00 | 1.37 | 0.01 | 0.03 | 0.04 | 3.81 | 10.53 | 3.42 | 26.07 | 0.00 | 0.51 | 14.41 | 33.07 | 0.00 | NA | NA | 0.18 |
| 807W3 | 807 | May | Arable | Y | N | L | D | 3 | 7.61 | 313 | 12.4 | 150.00 | 69 | 7.13 | 9.82 | 9.73 | 0.07 | 15.82 | 0.00 | 0.00 | NA | 1.27 | 0.02 | 0.06 | 0.05 | 3.46 | 14.77 | 5.67 | 33.58 | 0.00 | 0.85 | 15.98 | NA | NA | NA | NA | NA |
| 807W4 | 807 | Jun | Arable | Y | N | L | D | 3 | 7.16 | 320 | 18.1 | NA | 48.2 | 4.51 | 20.42 | 53.73 | 0.18 | 17.99 | 0.03 | 0.01 | NA | 1.19 | 0.07 | 0.21 | 0.11 | 3.93 | 16.99 | 6.47 | 45.97 | 0.02 | 1.20 | NA | 2.92 | NA | 138.80 | 348.80 | NA |
| 807W5 | 807 | Oct | Arable | Y | N | L | D | 3 | 6.94 | 271 | 10.9 | NA | NA | NA | 12.20 | 0.34 | 0.33 | 14.64 | 0.00 | 0.00 | NA | 1.33 | 0.08 | 0.24 | 0.25 | 4.45 | 16.42 | 5.62 | 34.82 | 0.00 | 4.09 | 16.75 | 37.61 | NA | 168.60 | 378.60 | NA |
| 892W2 | 892 | Mar | Arable | Y | N | L | D | 3 | 8.59 | 415 | 6.18 | 150.00 | 158.6 | 19.18 | 40.31 | 17.19 | 0.11 | 14.60 | 0.09 | NA | 0.00 | 2.39 | 0.08 | 0.24 | 0.03 | 18.68 | 24.04 | 11.52 | 66.97 | 0.00 | NA | 15.31 | 19.78 | 0.00 | NA | NA | NA |
| 892W4 | 892 | Jun | Arable | Y | N | L | D | 3 | 8.45 | 484 | 16.8 | 170.00 | 78.1 | 7.55 | 30.45 | 9.34 | 0.11 | 13.42 | 0.02 | 0.01 | NA | 1.23 | 0.11 | 0.34 | 0.00 | 9.67 | 10.37 | 9.51 | 57.45 | 0.01 | 0.03 | NA | 2.05 | NA | 161.90 | 371.90 | NA |
| 892W5 | 892 | Oct | Arable | Y | N | L | D | 3 | 7.15 | 389 | 10.5 | NA | 42.2 | 2.4 | 26.32 | 6.94 | 0.12 | 11.99 | 0.06 | 0.39 | NA | 1.53 | 0.12 | 0.35 | 0.01 | 9.80 | 12.72 | 9.95 | 49.90 | 0.00 | 0.02 | 13.26 | 33.41 | NA | 219.40 | 429.40 | NA |
| 893W2 | 893 | Mar | Arable | N | N | L | S | 3 | 7.70 | 757 | 7.08 | 30.00 | 103 | 12.2 | 29.54 | 51.65 | 0.05 | 9.65 | 10.76 | 0.28 | NA | 13.14 | 0.02 | 0.07 | 0.03 | 9.83 | 3.71 | 11.40 | 79.15 | 0.00 | 0.16 | 10.47 | NA | 0.00 | NA | NA | NA |
| 893W3 | 893 | Mar | Arable | N | N | L | S | 3 | 7.85 | 701 | 11.66 | 40.00 | 75.2 | 7.91 | NA | NA | NA | NA | NA | NA | NA | NA | NA | NA | NA | NA | NA | NA | NA | NA | NA | NA | NA | NA | NA | NA | NA |
| 893W4 | 893 | Jun | Arable | N | N | L | S | 3 | 8.45 | 484 | 16.8 | 10.00 | 78.1 | 7.55 | 36.79 | 50.10 | 0.16 | 12.00 | 2.91 | 0.01 | NA | 3.52 | 0.01 | 0.04 | 0.15 | 11.96 | 3.21 | 10.44 | 58.29 | 0.04 | 0.27 | NA | 57.30 | NA | 161.90 | 371.90 | NA |
| 907W3 | 907 | May | Grassland | N | N | S | D | 3 | 7.85 | 319 | 11.53 | 150.00 | 39.9 | 4.21 | 1.80 | 9.58 | 0.33 | 17.14 | 0.00 | 0.17 | NA | 2.05 | 0.24 | 0.73 | 0.10 | 3.09 | 8.18 | 4.36 | 43.83 | NA | 0.32 | 17.92 | NA | NA | NA | NA | NA |
| 907W4 | 907 | Jun | Grassland | N | N | S | D | 3 | 6.89 | 309 | 21.1 | NA | 38.9 | 3.45 | 3.27 | 8.19 | 0.34 | 14.40 | 0.01 | 0.01 | NA | 1.28 | 0.25 | 0.76 | 0.09 | 4.31 | 7.94 | 4.85 | 51.92 | 0.10 | 0.37 | NA | 51.30 | NA | 101.70 | 311.70 | NA |
| 907W5 | 907 | Oct | Grassland | N | N | S | D | 3 | 6.44 | 199.5 | 10.1 | NA | 16.3 | 1.8 | 2.21 | 1.74 | 0.51 | 19.37 | 0.00 | 0.01 | NA | 1.45 | 0.47 | 1.45 | 0.03 | 2.96 | 10.94 | 3.46 | 30.95 | 0.00 | 0.96 | 21.13 | 31.73 | NA | 202.30 | 412.30 | NA |
| 910W3 | 910 | May | Grassland | Y | N | L | S | 2 | 7.83 | 719 | 10.87 | 40.00 | 86.2 | 9.22 | 19.65 | 90.57 | 0.06 | 13.98 | 5.47 | 0.03 | NA | 7.24 | 0.04 | 0.13 | 0.02 | 9.81 | 2.23 | 11.56 | 114.82 | 0.00 | 0.20 | 16.36 | NA | NA | NA | NA | NA |
| 911W2 | 911 | Mar | Grassland | N | P | L | D | 3 | 8.40 | 776 | 5.71 | 125.00 | 73 | 9.02 | 47.69 | 76.85 | 1.32 | 20.34 | 0.92 | NA | 0.14 | 5.83 | 1.23 | 3.77 | 0.09 | 23.15 | 31.41 | 21.42 | 106.58 | 0.05 | NA | 21.32 | NA | 0.04 | NA | NA | NA |
| 911W3 | 911 | May | Grassland | N | P | L | D | 3 | 8.49 | 748 | 10.77 | 160.00 | 38.4 | 4.12 | 41.42 | 47.21 | 0.89 | 21.95 | 0.05 | 0.04 | 0.00 | 1.77 | 0.87 | 2.66 | 0.02 | 20.28 | 22.44 | 17.93 | 115.74 | 0.02 | 0.06 | 22.25 | 46.74 | 0.00 | NA | NA | 0.09 |
| 911W4 | 911 | Jun | Grassland | N | P | L | D | 3 | 7.43 | 839 | 19.8 | NA | 39.1 | 3.35 | 45.31 | 31.07 | 1.48 | 19.71 | 0.02 | 0.02 | NA | 1.66 | 1.17 | 3.59 | 0.31 | 18.09 | 20.35 | 14.87 | 99.89 | 0.07 | 0.11 | NA | 55.30 | NA | 166.20 | 376.20 | NA |
| 939W3 | 939 | May | Grassland | Y | N | M | S | 3 | 7.42 | 388 | 10.01 | 30.00 | 36.3 | 3.97 | 13.81 | 63.28 | 1.35 | 46.83 | 0.00 | 0.40 | NA | 4.10 | 1.35 | 4.15 | -0.01 | 8.69 | 18.55 | 6.79 | 55.76 | 0.00 | 0.14 | 47.19 | NA | NA | NA | NA | NA |
| 939W4 | 939 | Jun | Grassland | Y | N | M | S | 3 | 6.70 | 388 | 18.7 | NA | 27.2 | 2.48 | 12.27 | 6.99 | 1.49 | 46.06 | 0.04 | 0.02 | NA | 3.82 | 0.85 | 2.59 | 0.64 | 3.94 | 5.19 | 5.19 | 21.60 | 0.02 | 0.59 | NA | 3.75 | NA | 49.70 | 259.70 | NA |
| 940W2 | 940 | Mar | Grassland | Y | N | M | S | 3 | 8.64 | 390 | 0.94 | 15.00 | 61.8 | 8.6 | 17.44 | 76.05 | 0.36 | 19.87 | 2.05 | NA | 0.00 | 2.96 | 0.29 | 0.89 | 0.07 | 12.54 | 9.49 | 5.77 | 72.27 | 0.00 | NA | 21.72 | NA | 0.00 | NA | NA | NA |
| 940W3 | 940 | May | Grassland | Y | N | M | S | 3 | 7.38 | 451 | 8.81 | 40.00 | 8.6 | 0.96 | 16.46 | 46.30 | 0.33 | 31.96 | 0.04 | 0.33 | 0.00 | 2.73 | 0.22 | 0.67 | 0.11 | 9.45 | 3.33 | 6.90 | 95.46 | 0.00 | 0.16 | 33.17 | 97.03 | 0.00 | NA | NA | 0.37 |
| 940W4 | 940 | Jun | Grassland | Y | N | M | S | 3 | 6.96 | 509 | 16.6 | NA | 24.6 | 2.38 | 9.64 | 39.73 | 0.99 | 27.79 | 0.04 | 0.01 | NA | 2.01 | 0.49 | 1.49 | 0.50 | 6.15 | 2.25 | 3.09 | 40.71 | 0.10 | 0.04 | NA | 3.29 | NA | 76.10 | 286.10 | NA |
| 940W5 | 940 | Oct | Grassland | Y | N | M | S | 3 | 6.75 | 370 | 10.4 | NA | NA | NA | 8.11 | 2.41 | 0.35 | 22.09 | 0.00 | 0.00 | NA | 1.37 | 0.23 | 0.71 | 0.11 | 7.59 | 4.96 | 4.83 | 67.15 | 0.00 | 0.77 | 25.03 | 68.61 | NA | -79.10 | 130.90 | NA |
| 942W2 | 942 | Mar | Grassland | N | N | M | S | 3 | 8.68 | 486 | 9.51 | 30.00 | 90.9 | 11.5 | 20.73 | 76.96 | 0.02 | 9.17 | 0.08 | NA | 0.00 | 0.87 | 0.01 | 0.03 | 0.00 | 7.82 | 26.55 | 15.08 | 64.08 | 0.00 | NA | 10.58 | NA | 0.00 | NA | NA | NA |
| 942W3 | 942 | May | Grassland | N | N | M | S | 3 | 8.04 | 573 | 11.7 | 60.00 | 69.8 | 7.34 | 9.76 | 144.08 | 0.09 | 10.97 | 0.00 | 0.06 | NA | 1.01 | 0.07 | 0.21 | 0.02 | 5.42 | 15.64 | 11.43 | 77.65 | 0.00 | 0.12 | 12.88 | NA | NA | NA | NA | NA |
| 942W4 | 942 | Jun | Grassland | N | N | M | S | 3 | 7.65 | 562 | 17.5 | NA | 35.7 | 3.37 | 11.60 | 134.76 | 0.23 | 12.03 | 0.04 | 0.08 | NA | 1.40 | 0.08 | 0.24 | 0.16 | 3.56 | 8.09 | 4.90 | 33.98 | 0.10 | 0.23 | NA | 3.61 | NA | 45.20 | 255.20 | NA |
| 942W5 | 942 | Oct | Grassland | N | N | M | S | 3 | 8.23 | 397 | 10.4 | NA | NA | NA | 8.48 | 36.42 | 0.59 | 11.52 | 0.03 | 0.00 | NA | 1.18 | 0.58 | 1.76 | 0.01 | 7.39 | 13.35 | 8.22 | 60.83 | 0.00 | 0.05 | 12.66 | 27.95 | NA | 163.60 | 373.60 | NA |
| AW2 | A | Mar | Forest | N | Y | L | S | 3 | 8.65 | 665 | 4.34 | 30.00 | 63 | 8.02 | 28.11 | 132.83 | 0.05 | 19.65 | 3.70 | NA | 0.00 | 5.16 | 0.03 | 0.10 | 0.02 | 13.87 | 5.06 | 10.76 | 106.20 | 0.00 | NA | 21.27 | NA | 0.00 | NA | NA | NA |
| AW3 | A | May | Forest | N | Y | L | S | 3 | 7.60 | 659 | 8.18 | 40.00 | 43.4 | 4.96 | 29.30 | 101.12 | 0.05 | 31.28 | 0.37 | 0.07 | 0.00 | 2.56 | 0.02 | 0.06 | 0.03 | 18.55 | 11.88 | 13.50 | 121.55 | 0.00 | 0.10 | 32.13 | 77.15 | 0.00 | NA | NA | 0.44 |
| AW4 | A | Jun | Forest | N | Y | L | S | 3 | 7.58 | 602 | 19.32 | 30.00 | 17 | 1.52 | 25.89 | 66.52 | 0.14 | 32.04 | 0.01 | NA | NA | 2.43 | 0.05 | 0.15 | 0.09 | 16.30 | 0.95 | 10.36 | 70.83 | 0.03 | 0.13 | 34.00 | 95.10 | NA | NA | NA | NA |
| BW3 | B | May | Grassland | Y | N | M | S | 3 | 7.21 | 658 | 9.11 | 30.00 | 53.5 | 6.01 | 15.75 | 28.76 | 0.56 | 27.84 | 0.05 | 0.01 | 0.00 | 2.11 | 0.52 | 1.59 | 0.05 | 10.63 | 6.18 | 8.01 | 109.72 | 0.00 | 0.17 | 28.10 | 82.14 | 0.00 | NA | NA | 0.06 |
| BW4 | B | Jun | Grassland | Y | N | M | S | 3 | 7.31 | 581 | 22.17 | 15.00 | 14.6 | 1.23 | 9.65 | 2.47 | 1.87 | 42.25 | 0.01 | NA | NA | 3.40 | 0.54 | 1.65 | 1.33 | 10.35 | 4.60 | 8.33 | 84.85 | 0.04 | 1.05 | 44.95 | 125.00 | NA | NA | NA | NA |
| DW2 | D | Mar | Arable | Y | N | S | D | 3 | 7.73 | 173 | 9.63 | 70.00 | 100 | 11.19 | 22.66 | 142.73 | 0.04 | 15.28 | 5.18 | 0.38 | 0.05 | 7.69 | 0.01 | 0.02 | 0.03 | 9.60 | 9.55 | 8.05 | 76.45 | 0.00 | 0.09 | 15.54 | NA | 0.02 | NA | NA | 5.58 |
| DW3 | D | May | Arable | Y | N | S | D | 3 | 7.30 | 244 | 14.86 | 90.00 | 60.3 | 5.86 | 8.09 | 25.17 | 1.35 | 23.66 | 0.00 | 0.07 | NA | 2.78 | 1.25 | 3.82 | 0.11 | 2.98 | 17.84 | 5.10 | 25.18 | 0.00 | 0.51 | 25.08 | NA | NA | NA | NA | NA |
| DW4 | D | Jun | Arable | Y | N | S | D | 3 | 7.49 | 230 | 27.48 | 230.00 | 121.1 | 9.3 | 7.86 | 13.39 | 1.48 | 37.82 | 0.01 | NA | NA | 3.60 | 0.64 | 1.96 | 0.84 | 5.84 | 9.98 | 5.99 | 26.86 | 0.04 | 1.98 | 114.00 | 114.00 | NA | NA | NA | NA |
| EW3 | E | May | Arable | Y | N | M | S | 3 | 8.98 | 611 | 15.32 | 100.00 | 129.5 | 12.46 | 19.08 | 153.29 | 0.06 | 18.66 | 0.51 | 0.26 | NA | 2.67 | 0.03 | 0.09 | 0.03 | 12.62 | 4.58 | 11.39 | 91.79 | NA | 0.53 | 19.28 | NA | NA | NA | NA | NA |
| EW4 | E | Jun | Arable | Y | N | M | S | 3 | 7.58 | 695 | 25.1 | 30.00 | 59.8 | 4.78 | 28.48 | 89.85 | 0.55 | 32.43 | 0.00 | 0.15 | NA | 3.48 | 0.43 | 1.32 | 0.12 | 15.01 | 4.22 | 11.69 | 91.24 | 0.00 | 0.25 | NA | NA | NA | NA | NA | NA |
| FW2 | F | Mar | Arable | N | Y | S | S | 3 | 7.78 | 980 | 7.58 | 50.00 | 90 | 10.56 | 19.76 | 63.29 | 0.09 | 10.16 | 10.04 | NA | 0.41 | 12.29 | 0.07 | 0.21 | 0.02 | 8.75 | 21.62 | 6.14 | 97.17 | 0.00 | NA | 11.28 | NA | 0.12 | NA | NA | NA |
| FW3 | F | May | Arable | N | Y | S | S | 3 | 8.05 | 809 | 10.35 | 80.00 | 98 | 10.62 | 23.42 | 65.79 | 0.08 | 20.87 | 12.38 | 0.02 | 0.69 | 14.15 | 0.04 | 0.12 | 0.04 | 14.71 | 37.68 | 11.06 | 106.49 | 0.00 | 0.04 | 22.06 | 54.92 | 0.21 | NA | NA | 12.61 |
| FW4 | F | Jun | Arable | N | Y | S | S | 3 | 7.81 | 878 | 18.83 | 100.00 | 98 | 10.62 | 21.79 | 54.81 | 0.29 | 22.05 | 0.00 | 0.09 | NA | 2.09 | 0.29 | 0.90 | 0.00 | 13.16 | 30.47 | 9.99 | 91.99 | 0.00 | 0.05 | NA | NA | NA | NA | NA | NA |
| GW3 | G | May | Arable | N | N | M | S | 3 | 7.63 | 570 | 12 | 40.00 | 121.7 | 12.56 | NA | NA | NA | NA | NA | NA | NA | NA | NA | NA | NA | NA | NA | NA | NA | NA | NA | NA | NA | NA | NA | NA | NA |
| GW4 | G | Jun | Arable | N | N | M | S | 3 | 6.96 | 520 | 17.7 | 70.00 | 25.4 | 2.34 | 26.48 | 20.52 | 1.33 | 31.10 | 0.00 | 0.08 | NA | 2.59 | 1.10 | 3.37 | 0.23 | 12.02 | 28.43 | 8.28 | 53.33 | 0.00 | 0.20 | NA | NA | NA | NA | NA | NA |
| HW2 | H | Mar | Arable | N | Y | L | S | 2 | 7.54 | 1387 | 9.19 | 50.00 | 44 | 4.91 | 52.75 | 206.56 | 0.07 | 15.96 | 3.84 | NA | 0.00 | 5.04 | 0.04 | 0.12 | 0.03 | 23.71 | 6.15 | 22.73 | 152.29 | 0.00 | NA | 16.37 | NA | 0.00 | NA | NA | NA |
| HW3 | H | May | Arable | N | Y | L | S | 2 | 7.50 | 1635 | 8.13 | 30.00 | 21.4 | 2.42 | 42.69 | 162.69 | 0.12 | 20.16 | 0.21 | 0.40 | NA | 2.63 | 0.10 | 0.31 | 0.02 | 25.18 | 8.21 | 26.70 | 162.89 | 0.00 | 0.46 | 22.41 | NA | NA | NA | NA | NA |
| IW2 | I | Mar | Arable | Y | N | L | S | 2 | 8.00 | 874 | 10.91 | 30.00 | 99.8 | 10.57 | 24.95 | 79.16 | 0.25 | 11.39 | 7.16 | NA | 0.00 | 9.30 | 0.18 | 0.54 | 0.07 | 15.35 | 10.60 | 6.96 | 91.71 | 0.00 | NA | 12.94 | NA | 0.00 | NA | NA | NA |

| | | | General descriptors | | | | | | Physical and Chemical paramters | | | | | | | | | | | | | | | | | | | | | | | | | | | | |
|---|---|---|---|---|---|---|---|---|---|---|---|---|---|---|---|---|---|---|---|---|---|---|---|---|---|---|---|---|---|---|---|---|---|---|---|---|---|
| Sample Name | KH# | Month | Landuse | Reed | Canopy | Size | Depth | Perm. | pH | Cond. µS | Temp | Tiefe | O2 Sat | O2 mg/L | Cl | SO4 | TP | DOC | NO3-N | NH4-N | NO2 | TN | o-PO4-P | o-PO4 | TP | Na | K | Mg | Ca | Br | TFe | TOC | SAK | NO2-N | E0 | Eh | An-N |
| IW3 | I | May | Arable | Y | N | L | S | 2 | 7.35 | 700 | 8.25 | 30.00 | 20.1 | 3.41 | 27.19 | 95.48 | 0.85 | 33.22 | 0.73 | 0.31 | NA | 3.77 | 0.55 | 1.69 | 0.30 | 15.11 | 20.19 | 14.32 | 107.29 | 0.00 | 0.55 | 33.87 | NA | NA | NA | NA | NA |
| JW2 | J | Mar | Arable | Y | N | M | S | 3 | 8.25 | 403 | 6.92 | 40.00 | 28 | 3.3 | 13.82 | 70.28 | 0.08 | 15.07 | 0.15 | NA | 0.00 | 1.61 | 0.04 | 0.11 | 0.05 | 3.41 | 23.69 | 7.10 | 56.53 | 0.00 | NA | 15.85 | NA | 0.00 | NA | NA | NA |
| JW4 | J | Jun | Arable | Y | N | M | S | 3 | 6.76 | 246 | 22.22 | 5.00 | 11.8 | 0.99 | 11.38 | 2.10 | 0.21 | 17.93 | 0.00 | 0.01 | NA | 0.90 | 0.18 | 0.56 | 0.03 | 8.24 | 2.80 | 6.48 | 65.91 | 0.00 | 0.12 | NA | NA | NA | NA | NA | NA |
| KW2 | K | Mar | Forest | N | Y | M | S | 3 | 7.80 | 263 | 10.06 | 50.00 | 116 | 12.7 | 17.56 | 35.88 | 0.06 | 29.16 | 3.56 | 0.07 | 0.52 | 6.29 | 0.01 | 0.03 | 0.05 | 4.18 | 9.02 | 4.57 | 17.70 | 0.00 | 0.09 | 30.36 | 67.75 | 0.16 | NA | NA | 3.79 |
| KW3 | K | May | Forest | N | Y | M | S | 3 | 7.50 | 268 | 16.19 | 30.00 | 24.9 | 2.41 | NA | NA | NA | NA | NA | NA | NA | NA | NA | NA | NA | NA | NA | NA | NA | NA | NA | NA | NA | NA | NA | NA | NA |
| KW4 | K | Jun | Forest | N | Y | M | S | 3 | 6.86 | 236 | 20.03 | 15.00 | 2.9 | 0.79 | 12.92 | 0.33 | 1.33 | 58.46 | 0.00 | 1.09 | NA | 6.82 | 1.48 | 4.54 | -0.15 | 5.92 | 14.50 | 6.85 | 24.82 | 0.00 | 0.82 | NA | NA | NA | NA | NA | NA |
| LW2 | L | Mar | Arable | N | P | S | D | 3 | 7.60 | 443 | 9.63 | 70.00 | 132 | 14.6 | 26.56 | 65.81 | 0.05 | 8.41 | 12.90 | 0.02 | 0.00 | 14.12 | 0.03 | 0.09 | 0.02 | 8.25 | 7.40 | 6.78 | 71.67 | 0.00 | 0.23 | 9.33 | NA | 0.00 | NA | NA | 12.92 |
| LW3 | L | May | Arable | N | P | S | D | 3 | 8.33 | 405 | 13.94 | 60.00 | 34.5 | 3.5 | 13.59 | 78.90 | 1.57 | 27.68 | 0.05 | 0.17 | NA | 3.09 | 0.96 | NA | NA | 3.39 | 29.09 | 8.33 | 51.42 | NA | 0.15 | 33.67 | 92.14 | 0.00 | NA | NA | NA |
| LW4 | L | Jun | Arable | N | P | S | D | 3 | 7.39 | 304 | 19.64 | 25.00 | 38 | 3.38 | 19.96 | 31.69 | 1.33 | 27.85 | 0.00 | 0.36 | NA | 3.36 | 2.75 | 8.43 | -1.42 | 6.65 | 25.15 | 6.88 | 35.30 | 0.00 | 0.21 | NA | NA | NA | NA | NA | NA |
| MW2 | M | Mar | Arable | N | Y | M | S | 2 | 7.73 | 611 | 10.64 | 40.00 | 88 | 9.5 | 37.77 | 94.48 | 0.49 | 32.01 | 2.84 | 0.07 | 0.02 | 5.92 | 0.35 | 1.08 | 0.14 | 9.22 | 26.04 | 9.04 | 74.33 | 0.00 | 0.14 | 34.52 | 89.66 | 0.01 | NA | NA | 2.92 |
| MW3 | M | May | Arable | N | Y | M | S | 2 | 8.35 | 811 | 13.93 | 30.00 | 95.3 | 9.96 | 17.68 | 52.18 | 0.13 | 12.97 | 15.87 | 0.14 | NA | 18.08 | 0.05 | NA | NA | 13.39 | 2.61 | 15.64 | 75.01 | 0.06 | -0.02 | 14.15 | 37.40 | 0.07 | NA | NA | NA |
| NW2 | N | Mar | Arable | N | Y | M | S | 3 | 7.46 | 451 | 8.71 | 50.00 | 85 | 9.6 | 14.65 | 53.22 | 0.11 | 18.43 | 0.07 | 0.43 | 0.00 | 1.56 | 0.07 | 0.21 | 0.04 | 6.45 | 3.90 | 5.18 | 70.25 | 0.00 | 0.15 | 19.52 | 48.02 | 0.00 | NA | NA | 0.50 |
| NW3 | N | May | Arable | N | Y | M | S | 3 | 7.93 | 455 | 14.37 | 30.00 | 32.8 | 3.29 | 28.24 | 66.95 | 0.41 | 27.61 | 1.36 | 0.27 | NA | 4.20 | 0.25 | NA | NA | 12.12 | 16.79 | 8.78 | 55.92 | 0.02 | 0.12 | 29.84 | 84.38 | 0.03 | NA | NA | NA |
| NW4 | N | Jun | Arable | N | Y | M | S | 3 | 7.35 | 475 | 15.07 | 20.00 | 32.8 | 3.29 | 33.39 | 1.87 | 1.24 | 37.53 | 0.00 | 2.67 | NA | 6.84 | 3.94 | 12.09 | -2.71 | 10.03 | 27.78 | 7.52 | 54.80 | 0.00 | 1.04 | NA | NA | NA | NA | NA | NA |
| OW3 | O | May | Forest | Y | Y | L | S | 3 | 7.70 | 402 | 18.86 | 20.00 | 11.5 | 1.17 | 10.25 | 58.58 | 0.81 | 37.24 | 0.02 | 0.81 | NA | 3.03 | 0.47 | NA | NA | 5.75 | 5.61 | 5.29 | 56.74 | 0.05 | 0.47 | 38.73 | 137.49 | NA | NA | NA | NA |
| OW4 | O | Jun | Forest | Y | Y | L | S | 3 | 7.13 | 245 | 18.03 | 40.00 | 21.9 | 2.46 | 5.07 | 5.84 | 0.55 | 30.19 | 0.00 | 0.40 | NA | 2.65 | 0.46 | 1.41 | 0.09 | 4.38 | 1.05 | 3.70 | 40.82 | 0.00 | 0.39 | NA | NA | NA | NA | NA | NA |
| PW2 | P | Mar | Forest | Y | Y | L | S | 3 | 7.84 | 414 | 5.65 | 30.00 | 62.1 | 7.57 | 14.63 | 51.04 | 0.05 | 13.95 | 0.03 | 0.04 | 0.00 | 1.06 | 0.02 | 0.05 | 0.04 | 6.02 | 3.30 | 5.32 | 65.39 | 0.00 | 0.11 | 14.54 | 33.60 | 0.00 | NA | NA | 0.07 |
| PW3 | P | May | Forest | Y | Y | L | S | 3 | 7.79 | 494 | 13.6 | 10.00 | 15.8 | 1.61 | 16.24 | 35.74 | 0.46 | 16.55 | 0.03 | 0.19 | NA | 1.39 | 0.17 | NA | NA | 9.52 | 3.64 | 6.97 | 59.98 | 0.02 | 0.10 | 16.63 | 48.58 | NA | NA | NA | NA |
| PW4 | P | Jun | Forest | Y | Y | L | S | 3 | 7.36 | 295 | 17.13 | 10.00 | 20.6 | 1.93 | 11.61 | 7.20 | 0.27 | 23.10 | 0.00 | 0.78 | NA | 2.83 | 0.21 | 0.63 | 0.06 | 8.18 | 1.55 | 5.47 | 65.80 | 0.00 | 0.25 | NA | NA | NA | NA | NA | NA |
| QW3 | Q | May | Forest | N | Y | M | S | 3 | 7.66 | 296 | 13.37 | 30.00 | 24.06 | 2.51 | 8.62 | 31.79 | 0.88 | 38.91 | NA | 0.10 | NA | 1.94 | 0.49 | NA | NA | 4.58 | 7.28 | 5.20 | 48.22 | 0.04 | 0.57 | 42.02 | 145.71 | NA | NA | NA | NA |
| QW4 | Q | Jun | Forest | N | Y | M | S | 3 | 7.10 | 239 | 19.19 | 30.00 | 23.1 | 2.07 | 14.62 | 8.35 | 0.67 | 44.12 | 0.00 | 0.09 | NA | 2.72 | 0.56 | 1.71 | 0.11 | 13.20 | 4.44 | 3.89 | 38.67 | 0.00 | 0.68 | NA | NA | NA | NA | NA | NA |
| RW3 | R | May | Forest | N | Y | M | D | 3 | 7.17 | 214 | 13.34 | 70.00 | 21 | 2.15 | 5.86 | 47.77 | 0.31 | 32.57 | 0.01 | 0.23 | NA | 1.74 | 0.13 | NA | NA | 2.96 | 4.37 | 3.17 | 33.82 | 0.04 | 0.69 | 34.28 | 127.62 | NA | NA | NA | NA |
| RW4 | R | Jun | Forest | N | Y | M | D | 3 | 6.71 | 108 | 21.34 | 80.00 | 24.5 | 2.1 | 4.05 | 18.14 | 0.38 | 34.07 | 0.00 | 0.14 | NA | 2.44 | 0.25 | 0.77 | 0.13 | 2.86 | 3.08 | 2.24 | 30.91 | 0.00 | 1.17 | NA | NA | NA | NA | NA | NA |
| SW3 | S | May | Forest | N | P | M | S | 3 | 8.39 | 523 | 14.03 | 20.00 | 32.8 | 3.31 | 15.31 | 27.15 | 0.12 | 14.62 | 0.03 | 0.07 | NA | 0.95 | 0.07 | NA | NA | 9.30 | 3.32 | 7.37 | 56.30 | 0.02 | 0.05 | 14.93 | 43.51 | 0.00 | NA | NA | NA |
| SW4 | S | Jun | Forest | N | P | M | S | 3 | 7.63 | 485 | 17.51 | 40.00 | 27.8 | 2.58 | NA | NA | NA | NA | NA | NA | NA | NA | NA | NA | NA | NA | NA | NA | NA | NA | NA | NA | NA | NA | NA | NA | NA |
| TW2 | T | Mar | Forest | N | P | M | S | 2 | 7.23 | 86 | 6.14 | 25.00 | 46.6 | 5.61 | 11.14 | 25.96 | 0.33 | 26.92 | 0.04 | 0.04 | 0.00 | 1.67 | 0.26 | 0.78 | 0.07 | 3.67 | 5.14 | 2.24 | 38.70 | 0.00 | 0.28 | 27.82 | 96.79 | 0.00 | NA | NA | 0.08 |
