## Supplementary material for "From microbes to mammals: pond biodiversity homogenization across different land-use types in an agricultural landscape": Table S3 Sequence counts

| Sample | Eukaria | Bacteria | Archaea |
|---|---|---|---|
| 0W4 | 183149 | 338735 | 376775 |
| 1189L2 | 264926 | 356406 | 310251 |
| 1189S2 | 473555 | 192563 | 201826 |
| 1189W2 | 79664 | 641175 | 0 |
| 1189W3 | 418463 | 490723 | 292341 |
| 1189W4 | 247519 | 314464 | 252281 |
| 1228L2 | 357232 | 217493 | 0 |
| 1228L4 | 316891 | 603971 | 306007 |
| 1228S2 | 295147 | 143914 | 0 |
| 1228S4 | 285187 | 569201 | 318048 |
| 1228W2 | 84237 | 509013 | 0 |
| 1228W3 | 273798 | 339618 | 295498 |
| 1229W2 | 114986 | 330676 | 106123 |
| 1229W3 | 287072 | 330186 | 46517 |
| 1338L4 | 437740 | 593038 | 193432 |
| 1338S2 | 212965 | 268163 | 174973 |
| 1338S4 | 265544 | 679512 | 638271 |
| 1338W2 | 288036 | 511674 | 314124 |
| 1338W3 | 383168 | 371250 | 159852 |
| 133L2 | 326985 | 282754 | 374922 |
| 133S2 | 291248 | 295123 | 450404 |
| 135AW3 | 265588 | 335773 | 295120 |
| 135BW3 | 432613 | 449180 | 64302 |
| 136AW3 | 246815 | 378401 | 373324 |
| 136BW3 | 524813 | 299960 | 132990 |
| 149L2 | 407293 | 216184 | 66705 |
| 149S2 | 334391 | 256427 | 290813 |
| 149W2 | 292889 | 468639 | 0 |
| 149W3 | 534207 | 330883 | 500242 |
| 149W4 | 364482 | 361183 | 356957 |
| 1510L2 | 234439 | 221779 | 246924 |
| 1510S2 | 379384 | 283675 | 113956 |
| 1510S4 | 260522 | 329435 | 329739 |
| 1510SL1 | 351633 | 232938 | 93377 |
| 1510W3 | 378975 | 599303 | 135532 |
| 1590L2 | 462713 | 200119 | 265897 |
| 1590S2 | 402720 | 245094 | 63791 |
| 1590W1 | 512893 | 327063 | 281182 |
| 1590W2 | 714437 | 288542 | 0 |
| 1590W3 | 342331 | 353901 | 241933 |
| 1590W4 | 247443 | 367296 | 212478 |
| 1598L2 | 200174 | 276622 | 428245 |
| 1598S2 | 288308 | 240946 | 478666 |
| 1598W2 | 1119992 | 401137 | 705278 |
| 1598W3 | 324216 | 276077 | 323870 |
| 1598W4 | 336114 | 358914 | 544019 |

| Sample | Eukaria | Bacteria | Archaea |
|---|---|---|---|
| 1599L4 | 354118 | 515201 | 404216 |
| 1599S4 | 238243 | 572844 | 362520 |
| 1599W1 | 290174 | 380651 | 300701 |
| 1599W2 | 2079697 | 479829 | 133590 |
| 1599W3 | 329034 | 276676 | 197780 |
| 1604W2 | 229028 | 430989 | 376895 |
| 1604W3 | 375649 | 341514 | 55851 |
| 1604W4 | 343778 | 404210 | 373376 |
| 183L2 | 356618 | 227248 | 442879 |
| 183S2 | 421463 | 203169 | 444963 |
| 187L2 | 357014 | 236318 | 417785 |
| 187S2 | 274663 | 211830 | 422253 |
| 187W2 | 308160 | 544476 | 0 |
| 187W3 | 463911 | 660483 | 422959 |
| 187W4 | 296308 | 437210 | 416603 |
| 190L2 | 298559 | 206849 | 374334 |
| 190S2 | 261257 | 259680 | 414894 |
| 190W2 | 991539 | 474921 | 458619 |
| 190W3 | 265892 | 276410 | 176439 |
| 236L2 | 335904 | 206649 | 184816 |
| 236S2 | 500100 | 251229 | 476713 |
| 2484L2 | 224273 | 428316 | 251687 |
| 2484S2 | 227688 | 403560 | 226060 |
| 2484W1 | 297337 | 298185 | 335385 |
| 2484W2 | 399639 | 659539 | 215003 |
| 2484W3 | 353745 | 427094 | 285030 |
| 2484W4 | 389024 | 315637 | 0 |
| 2489L2 | 356627 | 450917 | 280778 |
| 2489S2 | 199450 | 412430 | 280528 |
| 2489W3 | 328785 | 314491 | 223684 |
| 2489W4 | 254198 | 351772 | 174576 |
| 2565S2 | 365406 | 187849 | 111766 |
| 2565SL2 | 270245 | 204899 | 90194 |
| 2565W2 | 916607 | 308667 | 0 |
| 2565W3 | 280644 | 262116 | 79539 |
| 2565W4 | 319301 | 289817 | 135979 |
| 258L1 | 277560 | 301863 | 214291 |
| 258L2 | 206800 | 447749 | 373061 |
| 258S1 | 182626 | 339007 | 247337 |
| 258S2 | 386985 | 349053 | 339714 |
| 258W1 | 221679 | 351380 | 0 |
| 258W2 | 231281 | 434472 | 38355 |
| 258W3 | 272973 | 358970 | 431725 |
| 258W4 | 243303 | 299508 | 293707 |
| 258W5 | 385828 | 444153 | 447905 |
| 259S1 | 439691 | 346108 | 161456 |

**Name Structure**

KH(L/S/W)(1-5)

| | |
|---|---|
| KH | KH name or number |
| L | Deep sediment |
| S | Surface sediment |
| W | Water |
| (1-5) | Campaign number |
| 1 | Dec-16 |
| 2 | Mar-17 |
| 3 | May-17 |
| 4 | Jun-17 |
| 5 | Oct-17 |

| Sample | Eukaria | Bacteria | Archaea |
|---|---|---|---|
| 259S2 | 211813 | 321458 | 0 |
| 259W1 | 281696 | 291738 | 379859 |
| 259W2 | 70492 | 528050 | 0 |
| 259W3 | 337910 | 383182 | 244911 |
| 259W4 | 251217 | 331156 | 354375 |
| 259W5 | 380735 | 321106 | 434528 |
| 265L1 | 431606 | 432670 | 167739 |
| 265L2 | 185675 | 416857 | 153590 |
| 265S1 | 386907 | 387317 | 216485 |
| 265S2 | 238149 | 353043 | 233711 |
| 265W1 | 297849 | 354152 | 188839 |
| 265W2 | 371670 | 278068 | 362514 |
| 265W3 | 378322 | 479131 | 308037 |
| 265W4 | 381971 | 408263 | 325794 |
| 265W5 | 280524 | 442300 | 388637 |
| 269L1 | 632470 | 427077 | 205113 |
| 269L2 | 246224 | 402854 | 36125 |
| 269S1 | 428488 | 358603 | 0 |
| 269S2 | 285739 | 507634 | 0 |
| 269W2 | 414677 | 428347 | 0 |
| 269W3 | 584085 | 453696 | 168527 |
| 269W4 | 234492 | 360926 | 244529 |
| 269W5 | 330024 | 325051 | 443067 |
| 275L1 | 559108 | 312068 | 239970 |
| 275L2 | 229433 | 379797 | 220372 |
| 275S1 | 554937 | 465907 | 240303 |
| 275S2 | 181560 | 430916 | 289687 |
| 275W1 | 318948 | 386314 | 0 |
| 275W2 | 512789 | 319469 | 0 |
| 275W3 | 284924 | 417404 | 501208 |
| 275W4 | 303349 | 368540 | 160311 |
| 275W5 | 277757 | 314591 | 106310 |
| 287L2 | 290696 | 367572 | 121593 |
| 287S2 | 189224 | 340318 | 0 |
| 287SL1 | 399805 | 177205 | 101701 |
| 287W2 | 196689 | 368073 | 42877 |
| 287W3 | 333981 | 531043 | 230380 |
| 287W4 | 238084 | 198302 | 307673 |
| 287W5 | 320263 | 274298 | 317817 |
| 28W2 | 660848 | 361595 | 130000 |
| 28W3 | 234312 | 199109 | 341231 |
| 28W4 | 228830 | 289283 | 78751 |
| 311L1 | 486525 | 499396 | 261332 |
| 311L2 | 337113 | 383799 | 136261 |
| 311S1 | 493278 | 325439 | 145678 |
| 311S2 | 223868 | 384853 | 232004 |

| Sample | Eukaria | Bacteria | Archaea |
|---|---|---|---|
| 311W3 | 377454 | 476585 | 147710 |
| 312L1 | 568667 | 545052 | 58585 |
| 312L2 | 299190 | 286620 | 311707 |
| 312S1 | 525172 | 338237 | 195854 |
| 312S2 | 281651 | 239836 | 231302 |
| 312W2 | 294347 | 378703 | 0 |
| 312W3 | 387087 | 290682 | 325281 |
| 312W4 | 365449 | 246398 | 221160 |
| 312W5 | 606677 | 339632 | 322078 |
| 319S4 | 319050 | 595081 | 327819 |
| 319W3 | 256145 | 277799 | 221407 |
| 40W2 | 355258 | 358129 | 0 |
| 40W3 | 340034 | 414249 | 201015 |
| 40W4 | 219936 | 279747 | 311032 |
| 606L2 | 435809 | 195944 | 217316 |
| 606S2 | 331334 | 215183 | 316302 |
| 606W2 | 357868 | 190596 | 0 |
| 606W3 | 397068 | 479383 | 209925 |
| 606W4 | 344229 | 313681 | 380486 |
| 607L2 | 371634 | 204865 | 97585 |
| 607S2 | 301478 | 207300 | 348383 |
| 607W2 | 106863 | 622818 | 0 |
| 607W3 | 586880 | 344971 | 204646 |
| 607W4 | 252556 | 0 | 445777 |
| 805L1 | 349006 | 260556 | 0 |
| 805L2 | 373286 | 132612 | 0 |
| 805S1 | 300618 | 247406 | 0 |
| 805S2 | 252989 | 278605 | 150119 |
| 805W2 | 486024 | 631476 | 0 |
| 805W3 | 476119 | 322146 | 66958 |
| 805W4 | 217902 | 223017 | 207575 |
| 807L1 | 475557 | 367732 | 215039 |
| 807L2 | 313467 | 239281 | 230930 |
| 807S1 | 284179 | 429728 | 231487 |
| 807S2 | 259918 | 210609 | 262900 |
| 807W1 | 198223 | 383037 | 59258 |
| 807W2 | 280062 | 692500 | 402968 |
| 807W3 | 503361 | 485805 | 309091 |
| 807W4 | 500903 | 343551 | 217001 |
| 807W5 | 278968 | 453337 | 313178 |
| 808L2 | 403040 | 353952 | 145647 |
| 808S2 | 426238 | 420507 | 279746 |
| 892L1 | 205892 | 283068 | 225037 |
| 892L2 | 205037 | 209518 | 0 |
| 892S1 | 206205 | 324965 | 133166 |
| 892S2 | 193282 | 254854 | 128018 |

| Sample | Eukaria | Bacteria | Archaea | Sample | Eukaria | Bacteria | Archaea |
|---|---|---|---|---|---|---|---|
| 311W1 | 214936 | 326657 | 244988 | 892W1 | 211794 | 358763 | 313336 |
| 892W2 | 376321 | 487602 | 336148 | 893W3 | 373548 | 334934 | 353821 |
| 892W4 | 250401 | 278730 | 287045 | 893W4 | 269581 | 366905 | 458827 |
| 892W5 | 417543 | 320562 | 343395 | 907L1 | 180172 | 304910 | 220986 |
| 893L1 | 276895 | 404318 | 284479 | 907L2 | 323638 | 246414 | 0 |
| 893L2 | 245185 | 424897 | 224924 | 907S1 | 265182 | 388376 | 236879 |
| 893S1 | 216757 | 518490 | 246710 | 907S2 | 343366 | 148445 | 229927 |
| 893S2 | 353355 | 471252 | 381808 | 907W1 | 343167 | 354975 | 104614 |
| 893W1 | 231711 | 367551 | 253338 | 907W2 | 87191 | 610365 | 0 |
| 893W2 | 289182 | 298105 | 381014 | 907W3 | 269650 | 316300 | 299671 |
| 907W4 | 351078 | 273289 | 267202 | BW3 | 329900 | 343906 | 365051 |
| 907W5 | 385399 | 347848 | 294922 | BW4 | 342067 | 191940 | 164218 |
| 908L2 | 244968 | 325862 | 299353 | C11 | 340416 | 1E+06 | 338266 |
| 908S2 | 233907 | 363026 | 275043 | C12 | 310506 | 758218 | 265870 |
| 910L1 | 401867 | 338135 | 200883 | C13 | 370843 | 578882 | 233536 |
| 910L2 | 271203 | 337281 | 203902 | C14 | 370858 | 890708 | 91991 |
| 910S1 | 287216 | 134094 | 0 | C15 | 304455 | 741392 | 288578 |
| 910S2 | 240006 | 332775 | 211760 | C16 | 468494 | 785913 | 487243 |
| 910S4 | 278497 | 514597 | 333652 | CL2 | 194058 | 400186 | 0 |
| 910W2 | 93185 | 113331 | 0 | CS2 | 283444 | 387627 | 41593 |
| 910W3 | 272236 | 370088 | 368302 | DW2 | 99837 | 606738 | 154797 |
| 911L1 | 241951 | 365755 | 161410 | DW3 | 396814 | 490912 | 327586 |
| 911L2 | 152957 | 319036 | 153312 | DW4 | 389067 | 366322 | 268890 |
| 911S1 | 217950 | 365935 | 0 | EW2 | 344805 | 341115 | 0 |
| 911S2 | 175631 | 182843 | 89921 | EW3 | 246368 | 311229 | 263703 |
| 911W1 | 277829 | 358714 | 265592 | EW4 | 339326 | 257446 | 274561 |
| 911W2 | 354337 | 393003 | 316938 | EW5 | 397598 | 333740 | 250071 |
| 911W3 | 409040 | 338847 | 387078 | FW2 | 78124 | 575989 | 153282 |
| 911W4 | 290262 | 245737 | 326831 | FW3 | 265940 | 130344 | 256970 |
| 939L1 | 335301 | 268892 | 0 | FW4 | 224255 | 256594 | 237965 |
| 939L2 | 257515 | 269519 | 0 | GW2 | 67715 | 84341 | 0 |
| 939S1 | 376597 | 259373 | 88240 | GW3 | 511799 | 545806 | 462906 |
| 939S2 | 236262 | 410127 | 0 | GW4 | 370597 | 338800 | 80918 |
| 939W2 | 112659 | 517061 | 0 | HL2 | 404642 | 239382 | 315759 |
| 939W3 | 311588 | 507629 | 150657 | HL4 | 229003 | 281448 | 313411 |
| 939W4 | 301645 | 249858 | 430558 | HS2 | 346933 | 220205 | 272708 |
| 940L1 | 241341 | 292903 | 0 | HS4 | 205672 | 277857 | 216487 |
| 940L2 | 253446 | 360684 | 178024 | HW2 | 299607 | 352836 | 446196 |
| 940S1 | 338930 | 310238 | 0 | HW3 | 266690 | 225957 | 660923 |
| 940S2 | 163322 | 287598 | 171191 | IW2 | 481013 | 465829 | 158467 |
| 940W2 | 472725 | 394150 | 51912 | IW3 | 308992 | 490914 | 358100 |
| 940W3 | 292122 | 314779 | 417520 | JS2 | 247398 | 237962 | 221749 |
| 940W4 | 199125 | 290439 | 242956 | JW2 | 926638 | 532683 | 63627 |
| 940W5 | 388999 | 339411 | 300202 | JW4 | 261153 | 306556 | 374640 |
| 942L1 | 332300 | 374229 | 185449 | KL2 | 359793 | 172345 | 90889 |
| 942L2 | 238601 | 365465 | 269257 | KS2 | 168107 | 210090 | 194075 |

| Sample | Eukaria | Bacteria | Archaea |
|---|---|---|---|
| 942S1 | 163635 | 294491 | 104520 |
| 942S2 | 193654 | 225092 | 261216 |
| 942W1 | 283158 | 304949 | 259565 |
| 942W2 | 386950 | 419576 | 399797 |
| 942W3 | 388869 | 397349 | 295183 |
| 942W4 | 397831 | 372577 | 443733 |
| 942W5 | 237704 | 304035 | 372861 |
| AL2 | 282275 | 343564 | 0 |
| AS2 | 323057 | 397459 | 101195 |
| AW2 | 365831 | 403015 | 95631 |
| AW3 | 377738 | 456835 | 389756 |
| AW4 | 232282 | 272324 | 253424 |
| BL2 | 340407 | 403227 | 40683 |
| BS2 | 229348 | 368709 | 218329 |
| BW2 | 303453 | 452424 | 0 |
| OW4 | 338704 | 329013 | 203091 |
| PL2 | 169992 | 421391 | 283086 |
| PS2 | 158404 | 474188 | 0 |
| PW2 | 1835623 | 424640 | 118091 |
| PW3 | 269534 | 337916 | 398867 |
| PW4 | 324612 | 281190 | 658385 |
| QL2 | 127540 | 218451 | 284333 |
| QS2 | 144208 | 409089 | 0 |
| QW2 | 1341696 | 355988 | 0 |
| QW3 | 289699 | 348351 | 281474 |
| QW4 | 320150 | 356004 | 168763 |
| RL2 | 155295 | 461678 | 187994 |
| RS2 | 136903 | 457300 | 0 |
| RW2 | 907197 | 285416 | 0 |
| RW3 | 384083 | 370668 | 278491 |
| RW4 | 287807 | 323440 | 545586 |
| SL2 | 178714 | 543781 | 308942 |
| SS2 | 126776 | 423256 | 273081 |
| SW2 | 263282 | 263282 | 0 |
| SW3 | 264469 | 284462 | 110911 |
| SW4 | 348624 | 501879 | 1008 |
| TL4 | 420817 | 464739 | 373322 |
| TS4 | 420583 | 632183 | 210343 |
| TW2 | 513741 | 345270 | 67914 |
| UW4 | 216831 | 359421 | 380273 |
| 607W4A | 0 | 359785 | 0 |
| 1484W4 | 0 | 0 | 95084 |

| Sample | Eukaria | Bacteria | Archaea |
|---|---|---|---|
| KW2 | 261090 | 382956 | 104605 |
| KW3 | 400973 | 355405 | 255064 |
| KW4 | 364754 | 241511 | 362172 |
| LW2 | 318196 | 602138 | 138879 |
| LW3 | 410901 | 349727 | 93975 |
| LW4 | 394068 | 368961 | 197530 |
| MW2 | 1617097 | 429502 | 223529 |
| MW3 | 450032 | 170856 | 240035 |
| NW2 | 499916 | 357920 | 176774 |
| NW3 | 301281 | 289462 | 119955 |
| NW4 | 282939 | 360392 | 336395 |
| OL2 | 191277 | 238386 | 237656 |
| OS2 | 129673 | 216269 | 0 |
| OW2 | 1308380 | 327183 | 0 |
| OW3 | 348915 | 283925 | 90979 |
