## Supplementary material for "From microbes to mammals: pond biodiversity homogenization across different land-use types in an agricultural landscape": Table S4 Indicator Species

**Table S4.** Top 5 bacterial and eukaryotic indicator species. Taxa were sorted by decreasing significance (see color legend) and association coefficient (Co.). The full results of the indicator species analysis are given in Supplementary dataset 2.

| | | Arable | | Forest | | Grassland | | Arable & Forest | | Arable & Grassland | | Forest & Grassland | |
|---|---|---|---|---|---|---|---|---|---|---|---|---|---|
| | | **Taxa** | **Co.** | **Taxa** | **Co.** | **Taxa** | **Co.** | **Taxa** | **Co.** | **Taxa** | **Co.** | **Taxa** | **Co.** |
| Presence Absence | *Bacteria* | *Acidovorax* sp. | 0.39 | *Rhodopseudomonas* sp. | 0.73 | *Cyanobium* sp. | 0.64 | *Alicyclobacillaceae* sp. | 0.57 | *Geitlerinema* sp. | 0.71 | unc. *Myxococcales* | 0.77 |
| | | *Comamonas* sp. | 0.39 | Unc. Forest soil bacterium (*Acetobacteraceae)* | 0.72 | *Commamonas* sp. | 0.61 | *Pseudahrensia sp.* | 0.63 | Cyanobium sp. | 0.66 | *Ramlibacter* sp. | 0.74 |
| | | *Pedobacter* sp. | 0.36 | *Burkholderia* sp. | 0.70 | *Nostoc* sp. | 0.60 | *Bacillus* sp. | 0.60 | *Erythromicrobium* sp. | 0.54 | *Cand.* Kaiserbacteria; | 0.74 |
| | | | | *Nocardioides* sp. | 0.67 | *Methanococcoides* sp. | 0.60 | *Gemmatimonas* sp. | 0.58 | *Nostoc* sp. | 0.65 | unc. *Rhizobiaceae* | 0.71 |
| | | | | Unc. *Sphingomonadaceae* | 0.66 | *Elusimicrobium* sp. | 0.58 | *Flavobacterium* sp. | 0.57 | *Pedobacter* sp. | 0.63 | *Mucinivorans* sp. | 0.68 |
| | Eukaryptes | | | *Helicoon* sp. | 0.67 | *Myxobolus sp.* | 0.48 | *Coccomyxa* sp. | 0.61 | *Chroomonas* sp. | 0.59 | *Chaetonotus* sp. | 0.66 |
| | | | | *Hydryphantes* sp. | 0.62 | *Ptychogastria* sp. | 0.48 | *Xironogiton* sp. | 0.49 | *Fistulifera* sp. | 0.52 | *Chondrostereum* sp. | 0.62 |
| | | | | *Ochrolechia* sp. | 0.61 | *Pterula* sp. | 0.45 | *Castrada sp.* | 0.49 | *Sigara* sp. | 0.51 | *Sorghum* sp. | 0.60 |
| | | | | *Macrostomum* sp. | 0.60 | *Nephroma* sp. | 0.41 | *Closterium* sp. | 0.48 | *Ankistrodesmus* sp. | 0.52 | *Coccophagus* sp. | 0.59 |
| | | | | *Chloroscypha* sp. | 0.59 | *Andreaea* sp. | 0.56 | *Oocystella* sp. | 0.47 | *Chlorella* sp. | 0.50 | *Marchandiomyces* sp. | 0.59 |
| Quantiattive | *Bacteria* | Comamonas sp. | 0.39 | *Rhodoluna* sp. | 0.95 | Unc. *Parcubacteria* | 0.99 | *Beijerinckia* sp. | 0.93 | *Polynucleobacter* sp. | 0.94 | *Novosphingobium* sp. | 0.93 |
| | | | | *Novosphingobium* sp. | 0.92 | *Calothrix* sp | 0.89 | *Pragia* sp. | 0.76 | *Tabrizicola* sp. | 0.88 | *Crenothrix* sp. | 0.90 |
| | | | | *Novosphingobium* sp. | 0.86 | Unc. Cyanobacterium | 0.85 | *Sphingomonas* sp. | 0.64 | *Hydrogenophaga* sp. | 0.83 | Unc. *Gammaproteobacteria* | 0.89 |
| | | | | *Microbacteriaceae* sp. | 0.84 | *Cyanobium* sp. | 0.79 | Unc. *Pseudahrensia* | 0.63 | Unc. *Sporichthyaceae* | 0.87 | *Sphingomonas* sp. | 0.89 |
| | | | | *Sphaerotilus* sp. | 0.84 | *Methanogranum* sp. | 0.78 | *Phyllobacterium* sp. | 0.60 | *Persicitalea* sp. | 0.78 | *Phenilobacterium* sp. | 0.89 |
| | Eukaryptes | Unc. *Leucocryptos* | 0.44 | *Helicascus* sp. | 0.94 | *Pterospora* sp. | 0.44 | *Aspergillus* sp. | 0.84 | Unc. *Cryptomycota* | 0.86 | *Pseudodiaptomus* sp. | 0.73 |
| | | | | *Chromophyton* sp. | 0.87 | *Roya* sp. | 0.41 | *Castrella* sp. | 0.86 | *Rhabditida* sp. | 0.76 | *Isotomiella* sp. | 0.68 |
| | | | | *Laevapex* sp. | 0.82 | *Eucyclops* sp. | 0.83 | *Daphnia* sp. | 0.70 | Unc. *Pedinellales* | 0.67 | *Pinnularia* sp. | 0.64 |
| | | | | *Trapelia* sp. | 0.79 | *Diplodinium* sp. | 0.57 | *Vorticella* sp. | 0.81 | *Cryptomonas* sp. | 0.85 | Unc. *Harpacticoida* sp. | 0.63 |
| | | | | *Helicoon* sp. | 0.74 | *Haplosporidium* sp. | 0.54 | *Pinnularia* sp. | 0.71 | *Amphora* sp. | 0.65 | *Coccophagus* sp. | 0.59 |

| | | |
|---|---|---|
| P<=0.001 | 0.001<P<=0.01 | 0.01<P<=0.05 |
